# AID/APOBEC3 Dynamic Catalytic Pockets and AID-α7 Regulation Mechanisms

**DOI:** 10.1101/2025.09.29.679391

**Authors:** David Nicolas Giuseppe Huebert, Justin J. King, Mani Larijani

**Affiliations:** Structural Biology and Immunology Program, Department of Molecular Biology and Biochemistry, Faculty of Science, Simon Fraser University, Burnaby, BC V5A 1S6, Canada; Immunology and Infectious Diseases Program, Division of Biomedical Sciences, Faculty of Medicine, Memorial University of Newfoundland, St. John’s, NL A1C 5S7, Canada

**Keywords:** AID/APOBEC3, Molecular Dynamics, Catalytic Pocket, AID-α7, Protein Regulation

## Abstract

AID/APOBEC3s mutate dC to dU in ssDNA in the restriction of viruses, creation of antibodies, and when off-target affecting cancers. Little was known of the catalytic pocket dynamics of these enzymes in real time. As such, we utilised molecular dynamics analyses to determine catalytic pocket volumes and states. We found that most AID/APOBECs remain predominantly in either a closed or indeterminate state to varying degrees while A3G-CD2 is predominantly open. This occurs chiefly from the catalytic residues themselves and the loop 1 serine/threonine in the “floor” of the catalytic pocket while other residues of the secondary catalytic loops 1, 3, 5, and 7 aid in these states. Surprisingly, we found that the presupposed stable position of AID’s unique α7 (β-state) in contact with the β-sheet and α6 was unstable while the α-state in contact with α6 and loop7 was stable and caused pocket closure. Mutations could cause the destabilisation of the α-state (R171Y and R178D) or stabilisation of the β-state (R190D and R194A). Thus, pocket states are relatively similar in overall cause; however, multiple residues from secondary catalytic loops as well as AID-α7 determine the rates at which these states occur, potentially affecting the chances any AID/APOBEC is able to have catalysis.

## 1. Introduction

AID is part of the apolipoprotein B mRNA editing enzyme catalytic polypeptide (AID/APOBEC) family of Zn-dependent cytidine deaminases (CDA), comprising 11 human members: AID, APOBEC1, APOBEC2, APOBEC4, and APOBEC3 subfamily members A through H (excluding E) [1–3]. These enzymes deaminate deoxycytidine (dC) to deoxyuridine (dU), primarily in single-stranded DNA (ssDNA) [1, 4–17]. AID acts on immunoglobulin (Ig) loci to initiate somatic hypermutation (SHM) and class-switch recombination (CSR), thereby enhancing antibody affinity and isotype diversity [1, 4, 18, 19]. AID deficiency causes a mild to severe, treatable form of hyper-IgM syndrome (HIGM), marked by absence of high-affinity, class-switched antibodies [4, 19, 20]. APOBEC3 (A3) enzymes function both in an innate and adaptive manner as anti-retroviral restriction factors, mutating viral and retroelement genomes [21–23]. A3C, A3D, A3F, A3G, and A3H (haplotypes II, V, VII) operate individually or together to block cross-species transmission [23–32]. However, off-target mutations by AID, A3A, A3B, and A3H (haplotype I) can trigger genome-wide mutations and double-strand breaks, contributing to tumorigenesis [33–54]. Due to their dual role in immunity and cancer, considerable effort has gone into understanding the structure, dynamics, and function of AID/APOBEC3s, particularly in the last decade.

All AID/APOBEC3s share a conserved core: a central β-sheet of five strands flanked by six or seven α-helices, connected by 12-13 variable-length loops (Figure 2.A,C) [10, 28, 54–62]. The catalytic pocket features a Zn ion coordinated by two cysteines and a histidine, along with a glutamate that serves as a proton shuttle, typically located at the N-termini of α2 and α3 (Figure 2.A,C). In 2015, our lab identified 21 “secondary catalytic” residues forming the pocket’s “walls” and “floor,” which stabilise dC binding [55]. These residues reside on four highly flexible loops (ℓ1, ℓ3, ℓ5, and ℓ7) collectively called secondary catalytic loops. Their importance was subsequently confirmed in AID crystal structures, and identical/homologous residues were identified in APOBEC3s [10, 28, 54, 56–58, 60–63].

**Figure 1.**
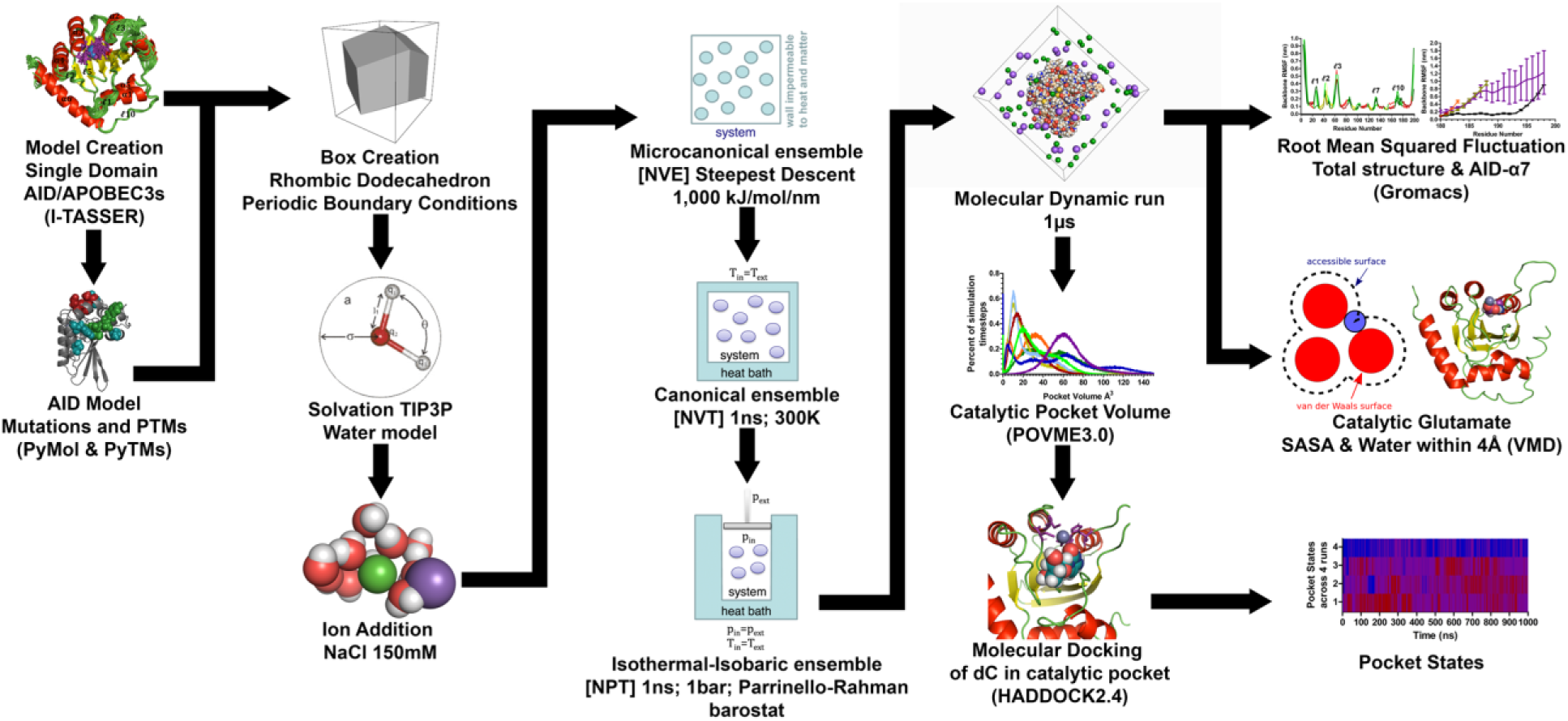
Computational workflow of primate A3s in MD using Gromacs.

**Figure 2.**
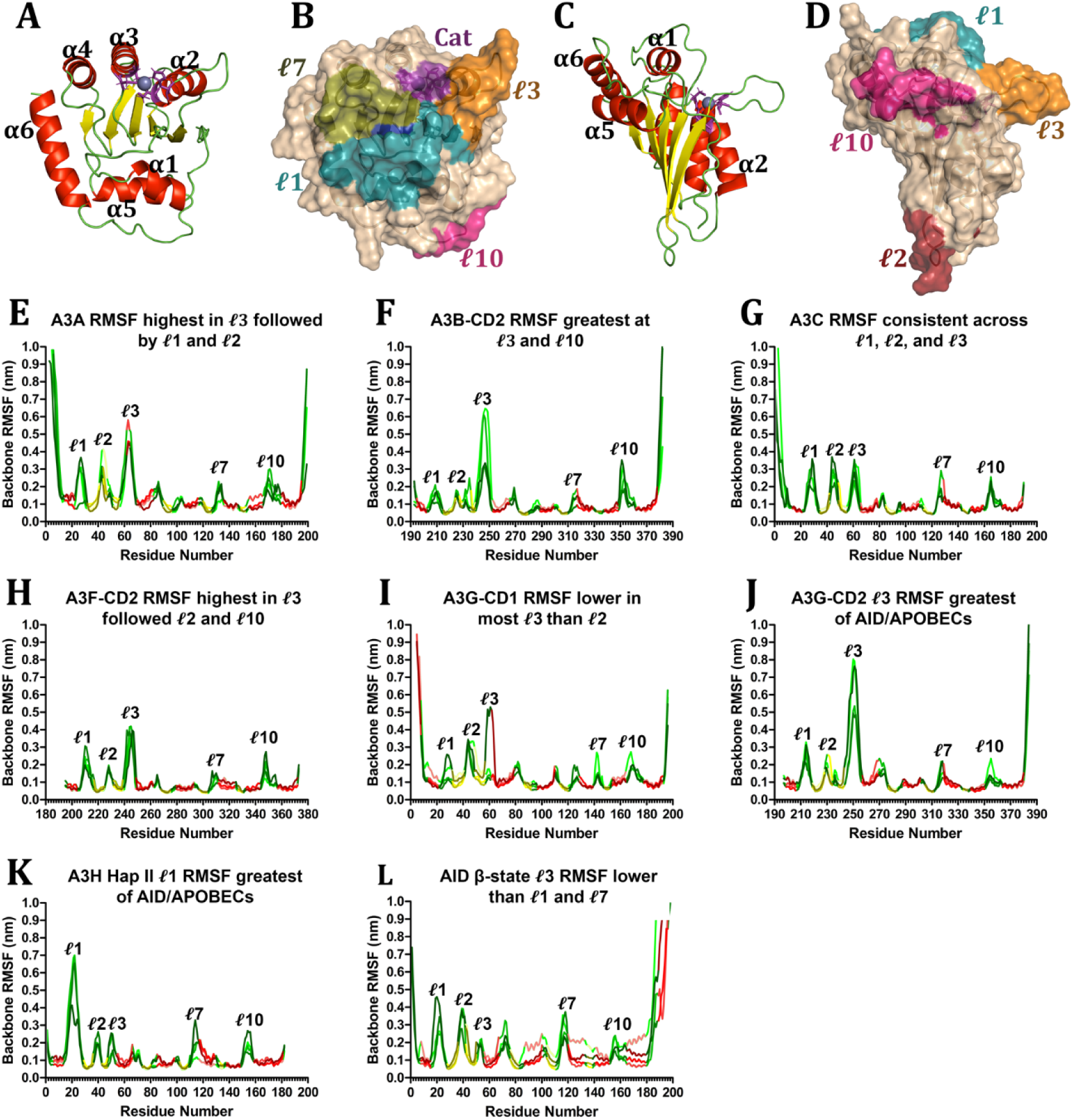
RMSF of AID/APOBEC3. 3D structural homology model of A3C shown from (A-B) over the pocket and (C-D) side on view showing either (A,C) secondary structure or (B,D) highlighting of the catalytic residues (purple), loop ℓ1 (deepteal), loop ℓ2 (firebrick), loop ℓ3 (orange), loop ℓ5 (blue), loop ℓ7 (deepolive), and loop ℓ10 (hotpink). (E-L) RMSF of multiple runs of AID/APOBEC3 proteins plotted highlighting loops (green), α-helices (red), and β-strands (yellow) with different shades representing different runs. Graphs are of (E) A3A, (F) A3B-CD2, (G) A3C, (H) A3F-CD2, (I) A3G-CD1, (J) A3G-CD2, (K) A3H haplotype II, and (L) AID with the α7 in the β-state.

Driven by loop flexibility, we found AID’s catalytic pocket shifts between open (25%) and closed (75%) conformations [55, 64]. Altering this dynamic through constructing AID variants modulated AID activity, establishing a direct structure-function correlation [55]. We termed this equilibrium “Schrödinger’s CATalytic pocket” mediated by secondary catalytic loops [64]. This dynamic was later validated via NMR and crystallography in AID and tumorigenic relatives A3A and A3B [11, 13, 65, 66]. These findings suggest that catalytic pocket closure serves evolutionarily as an auto-inhibitory mechanism to limit genomic damage.

The secondary catalytic loops also underlie many functional distinctions among AID/APOBEC3s. Loops ℓ1, ℓ3, and ℓ7 vary most in sequence, length, and compaction relative to the enzyme core (Figure 2.A-C) [11, 17, 23, 28, 55, 56, 58, 66, 67]. These loops contribute to dC binding, ssDNA interaction, and overall activity. Substrate specificity is dictated by loops ℓ1 and ℓ7 [13, 16, 67–74]. Loop ℓ1 contains a conserved “gatekeeper” residue (R25 in AID, positive or polar in A3s), which orchestrates ssDNA backbone interactions at the catalytic pocket mouth [10, 55, 66]. Loop ℓ3 is implicated in dimerization interfaces across several family members including AID, A3A, A3F, A3G, and A3H [29, 53, 62, 75–77].

Beyond these loops, several regions outside the catalytic pocket are essential for AID-specific functions. The assistant patch comprised of four arginine residues (R171, R174, R177, R178) on α6 is unique to AID and essential for CSR; its mutation abrogates CSR [10]. Positioned on the “front” of AID, the assistant patch extends the DNA-binding groove and enables binding to branched DNA and G-quadruplexes, but not linear ssDNA. CSR requires AID localization to immunoglobulin via switch-regions G4 quadruplexes [10, 78]. AID also possesses a unique 18-residue C-terminal region predicted to form α-helix (α7) not found in other APOBEC3s [55, 64, 75]. Although two crystal structures of AID have been solved [10, 79], neither resolved α7’s structure or position. Nonetheless, α7 is essential for CSR, as shown by both experimental variants and human HIGM cases with α7 deletions [20, 80]. To address this structural gap, our lab and others developed 3D homology models, predicting α7 to lie on the “back” of AID, opposite the catalytic pocket [55, 81]. Alternative models propose α7 on the “front,” near the assistant patch and interacting with ℓ7 [55]. In addition to DNA-G4 assistant patch binding, AID also binds RNA G-quadruplexes derived from immunoglobulin switch-region transcripts via a putative RNA-binding domain (residues 130–138) [82–84]. This region, located on the enzyme’s “back,” stabilises the dominant “back” conformation of α7 and overlaps with several CSR-disruptive HIGM mutations [20, 80, 85].

## 2. Materials and methods

### 2.1. Free adenine-derived nucleo -base/-side deaminases

I-TASSER (http://zhanglab.ccmb.med.umich.edu/I-TASSER/) was used to generate the AID/APOBEC 3D structural homology models in the same manner as our previous publication [86–89]. Due to AID/APOBEC enzymes often requiring truncations or mutations for crystallization, computational modeling was the most suitable method for producing wildtype structures. Structural templates were selected from the Protein Data Bank (http://www.rcsb.org) and visualised using PyMOL v2.5.4 (http://www.pymol.org/). For each protein, four structural models were generated from which those with open catalytic pockets were prioritised, except in one group of AID and A3G-CD1. These models contained Zn²⁺ ions placed within the catalytic pocket using I-TASSER’s COACH program [90, 91]. Templates for model construction included: A3A (5KEG, 2M65, 5SWW, 4XXO) [11, 53, 54, 66], A3B-CD2 (6NFK, 5CQK, 5CQI) [65, 92], A3C (5ZVA, 6NIL, 5W2M) [14, 26, 63], A3F-CD2 (5HX5, 5W2M, 3WUS, 5ZVB) [14, 59, 63, 93], A3G-CD1 (5K81, 5K82, 5K83) [28], A3G-CD2 (4ROV, 3E1U, 3IR2) [15, 23, 77], A3H Hap II (5Z98, 6B0B, 5W45) [17, 62, 94], and several AID conformations: β-state (5W0R, 5W0Z) [10], α-state (5W0R, 5W0Z, 5W1C, 5W0U) [10], and closed pocket β-state (5W0U, 5W0Z, 5W1C, 5JJ4) [10, 79].

All models were manually corrected to ensure catalytic cysteines were deprotonated to the thiolate form. Thiolates are critical for coordinating Zn²⁺ and for avoiding steric clashes caused by hydrogen atoms overlapping with the metal’s Van der Waals radius. Notably, some α7s in AID were unstable, despite being built from the same templates and dynamics parameters. To explore α7 dynamics, we introduced mutations and post-translational modifications (PTMs) using PyMOL’s mutagenesis wizard and PyTMs plugin, respectively [95]. R-group rotamers were preserved unless steric clashes occurred, in which case alternate rotamers with minimal overlap were selected, assuming equilibration would resolve any minor deviations.

### 2.2. Molecular dynamics setup

Molecular dynamics (MD) simulations were performed using the GROMACS 2020.2 suite [96, 97] in combination with the CHARMM36 all-atom force field [98]. GROMACS was selected for its flexibility, extensive plugin support, and capacity to customise simulations, while CHARMM36 was used to ensure compatibility with previous lab work and for its strong reputation in modeling protein systems in aqueous environments. Each simulation began by positioning the holoenzyme 3D model in the centre of a rhombic dodecahedron box. This shape was selected because it reduces the simulation volume to approximately 71% of that of a cubic box, thereby improving computational efficiency. Periodic boundary conditions (PBCs) were applied to replicate the system across all faces of the box, ensuring that each dodecahedron (with 12 faces, 14 vertices, and 24 edges) connects with its mirror images. The box boundaries were placed 2 nm away from the holoenzymes initial position to prevent steric interference between the original structure and its periodic images, as nonbonded interaction cutoffs were set to 1.2 nm and periodic images were 4 nm apart.

The system was solvated using the TIP3P water model, which is compatible with CHARMM36 due to its inclusion of Lennard-Jones potentials on both oxygen and hydrogen atoms [99, 100]. To neutralise system charge and replicate physiological ionic strength, Na⁺ and Cl⁻ ions were added to a final concentration of 150 mM [101]. The final solvated systems contained ∼20,000 solvent and ion molecules and approximately 70,000 atoms, depending on the AID/APOBEC variant modelled. The specifics of the equilibrations and simulations were done through the modifications of GROMACs molecular dynamics parameter files; however, many variables were determined by the use of the CHARMM36 forcefield. Long-range Coulomb interactions were done through the use of the particle mesh Ewald (PME) method [102].

Energy minimization was performed using the steepest descent algorithm within the microcanonical (NVE) ensemble until the maximum force on any atom was less than 1,000 kJ/mol/nm or until 50,000 steps were completed with a minimum step size of 0.01 nm. This step helped identify the system’s local energy minimum by optimizing the geometry of both protein and solvent. Following minimization, the system underwent a 1 ns equilibration under the canonical (NVT) ensemble using 2 fs time steps at 300 K. A subsequent 1 ns equilibration under the isothermal-isobaric (NPT) ensemble was performed using the Parrinello-Rahman barostat to stabilise pressure at 1 bar [103]. These temperature and pressure conditions are consistent with those used in the literature and previous AID/APOBEC simulations. The final production MD simulation was run for 1 μs, with trajectory data saved every 10 ps, to capture detailed protein dynamics and assess catalytic pocket breathing over time. Visualization was post-processed to centre the protein and remove rotational drift for clearer analysis.

### 2.3. Molecular dynamics analysis

Each molecular dynamics (MD) simulation was analysed for structural stability using key metrics: radius of gyration, root mean square deviation (RMSD), and root mean square fluctuation (RMSF), all calculated via the GROMACS suite. These parameters assessed both overall protein conformation and localised fluctuations at the secondary structure level. Visual Molecular Dynamics (VMD 1.9.3) was used to inspect trajectories, facilitate visual interpretation, and perform additional calculations requiring precise selection algebra [104]. To assess the accessibility of the catalytic pocket, we used VMD to measure the solvent-accessible surface area (SASA) and to count the number of water molecules within 4 Å of the catalytic glutamate’s two R-group oxygen atoms. This provided insights into the proton shuttle’s hydration and accessibility (Figure 5). Pocket volumes were calculated using POVME 3.0, a program that computes volume by filling enzyme-free space with voxels, using a grid spacing of 0.5 Å³ per voxel [105]. We placed two 2.5 Å diameter seed spheres to define the core catalytic region of each AID/APOBEC variant. These were surrounded by inclusion spheres, defining the following maximum volume limits: set at 4.0 Å for A3A and A3G-CD2; 3.5 Å for A3B-CD2 and AID; and 3.0 Å for A3C, A3F-CD2, A3G-CD1, and A3H Hap II. The seed spheres established regions that should always remain open, while the inclusion spheres captured potentially open conformations. POVME identified all contiguous, unoccupied voxels connected to the seed spheres from which the total pocket volume at each simulation timepoint was computed.

To classify pocket states, representative timepoints were selected at even 5 Å³ volume intervals. At each of these timepoints, the corresponding 3D structural homology model was extracted and docked with dC using HADDOCK 2.4 under default parameters in a similar manner to our previous paper [86, 106, 107]. dC and the catalytic glutamate were defined as docking targets to evaluate whether the substrate could enter the catalytic pocket across all 4 simulations per protein. The resulting docking data allowed us to classify each pocket state as "open," "closed," or "indeterminate" based on whether docking was consistently successful, inconsistent, or unsuccessful across all four homology models for each volume. After establishing preliminary boundaries between these pocket states using 5 Å³ intervals, we refined them by conducting additional dockings at 1 Å³ intervals within the identified transition zones. This approach allowed us to pin-point more precisely the specific pocket volumes associated with each conformational state, providing a detailed view of how pocket accessibility fluctuates during simulations.

## 3. Results

### 3.1. Fluctuations of AID/APOBEC3 secondary structures consistent in DNA-binding groove

Each member of the AID/APOBEC family shares the characteristic APOBEC fold, which consists of a stable core structure flanked by variable secondary elements in terms of sequence length and diversity [79]. We aimed to explore the extent of fluctuation within these structural elements across family members with a focus on how these variations might impact the DNA-binding groove and surrounding protein surfaces. Our initial hypothesis was that the core fold would remain largely conserved due to its structural stability, while dynamic changes would be localised to the surface-exposed loops, particularly those involved in substrate binding (ℓ1, ℓ3, ℓ5, and ℓ7). The APOBEC fold’s central β-sheet and its six α-helices exhibited the highest stability across all proteins examined, as expected (Figure 2). In contrast, loops (especially ℓ1, ℓ2, ℓ3, and ℓ10) showed the most significant fluctuations. Loops ℓ1 and ℓ3 are central to the DNA-binding groove architecture, directly contributing to catalysis, whereas loops ℓ2 and ℓ10 are positioned further from the catalytic core (Figure 2.A-D). Specifically, ℓ1 contacts the DNA backbone and helps position the target cytidine within the catalytic site while also facilitating interactions with the other groove-forming loops. Loop ℓ3 plays a role in DNA stabilisation at the loop 1/3 interface and also has known functions in dimerization [66, 76]. Meanwhile, ℓ2 resides opposite the catalytic pocket, nestled between two β-strands and exposed to solvent with minimal catalytic involvement. Loop ℓ10, which in double-domain deaminases can interact with the linker, is situated between α5 and α6 and is also solvent-exposed but to a lesser degree.

Among the groove-associated loops, ℓ1 and ℓ3 are markedly more dynamic than ℓ5 and ℓ7 (Figure 2.E-L). The extent of fluctuation in these loops varies between family members. In A3A, A3B-CD2, A3F-CD2, and A3G-CD2, ℓ3 displays more fluctuation than ℓ1. Conversely, in A3C, both loops exhibit roughly equal fluctuation, while in AID, A3H hap II, and A3G-CD1 (except one simulation run), ℓ1 is more dynamic. The variability in ℓ3’s movement correlates with amino acid length which ranges across the following proteins: A3G-CD1 (4 residues), AID (5), A3H hap II (6), A3F-CD2 (9), and A3C (10). Interestingly, while the A3Z1 subtypes all share a ℓ3 length of 13 residues, A3G-CD2 has significantly higher fluctuations than A3A and A3B-CD2. Loop ℓ1’s flexibility also generally corresponds with its length. Yet, there are exceptions; A3G-CD1 and A3B-CD2, each with ℓ1 lengths of 10 and 12 residues, respectively, exhibit the lowest fluctuations. Meanwhile, A3F-CD2, A3G-CD2, and AID (all with 12 residues) and A3C with 13 residues, show higher flexibility. Notably, A3H hap II has the most dynamic ℓ1 despite its 15-residue length, followed unexpectedly by A3A, which only has a 9-residue loop.

Loop ℓ5, which supports ℓ7 and helps define the catalytic pocket’s architecture, is highly conserved and stable among all family members. Only A3F-CD2 shows marginally more stability, while A3A is slightly increased in flexibility. Loop ℓ7, critical for substrate specificity, has a well-conserved RL or RI motif at its N-terminal end that interacts with ℓ1. However, its C-terminal region is more solvent-exposed as it transitions into α4 and eventually toward α6. Despite ℓ7 lengths ranging from 7 to 9 residues, fluctuations are fairly consistent with A3F-CD2 being the most stable and A3C and A3H hap II being the most dynamic. Interestingly, AID in the α7 β-state, where α7 does not interact with ℓ7, shows even greater ℓ7 fluctuations than any APOBEC3 member. Together, these data suggest that while the structural framework of ℓ5 and ℓ7 is relatively conserved, ℓ1 and ℓ3 contribute significantly to the dynamic flexibility of the DNA-binding groove.

Beyond the catalytic groove, fluctuations in ℓ2 and ℓ10 further illustrate structural variability. Loop ℓ2, found between β1 and β2 on the “back side” of the protein, is notably solvent exposed, a condition exacerbated by variations in β-strand lengths, presence of ℓ4, and α2 length and positioning. Fluctuations are highest in A3A, due to its extended β-strands, and in AID, which features a prominent ℓ4 and a five-residue ℓ2. In contrast, A3B-CD2 and A3F-CD2 display minimal movement in this region. Loop ℓ10 lies near the DNA-binding groove beyond the loop 1/3 interface. The N-terminal portion of ℓ10, exiting α5 and moving toward α1 before transitioning into α6, shows moderate fluctuations across all proteins. Though solvent-exposed, ℓ10 never reaches the dynamic levels seen in ℓ1 and ℓ3. Finally, A3G-CD1’s ℓ8 stands out with greater flexibility than in any other AID/APOBEC tested due to its unusual length of 8 residues compared to the typical 2-3 residues seen in other family members. This extended loop may influence interactions with the CD2 domain’s β2 β-bulge.

### 3.2. AID/APOBEC3 pocket volumes shown to be greatest in AID, A3A, and A3G-CD2

To better understand the behavior of the catalytic pocket in AID/APOBEC proteins, we examined several parameters across the following 1 µs molecular dynamics simulations: the SASA of the oxygens of the catalytic glutamate, the number of water molecules within 4 Å of those atoms, and the volume of the catalytic pocket. These metrics provide insight into how available the catalytic glutamate is to serve as a proton shuttle, how accessible the pocket is to water molecules that may be hydrolyzed, and how much spatial accommodation exists for nucleotide binding, as well as the variability of the pocket’s size over time. We hypothesised that the A3Z1 subtypes, known to have the highest catalytic activity, would display the greatest SASA values and water accessibility for their proton shuttles. In contrast, we expected lower values for A3Z2 and A3Z3 subtypes, and minimal accessibility in AID and A3G-CD1 due to their low or absent catalytic activities. Consistent with our hypothesis, we observed that, although the catalytic glutamate’s SASA could occasionally reach higher values, 75% of the data across all proteins showed SASA values below 15 Å² with the most frequently occurring SASA being below 10 Å² (Figure 3.A-B). This indicates that the SASA distributions for the catalytic glutamates are positively-skewed.

**Figure 3.**
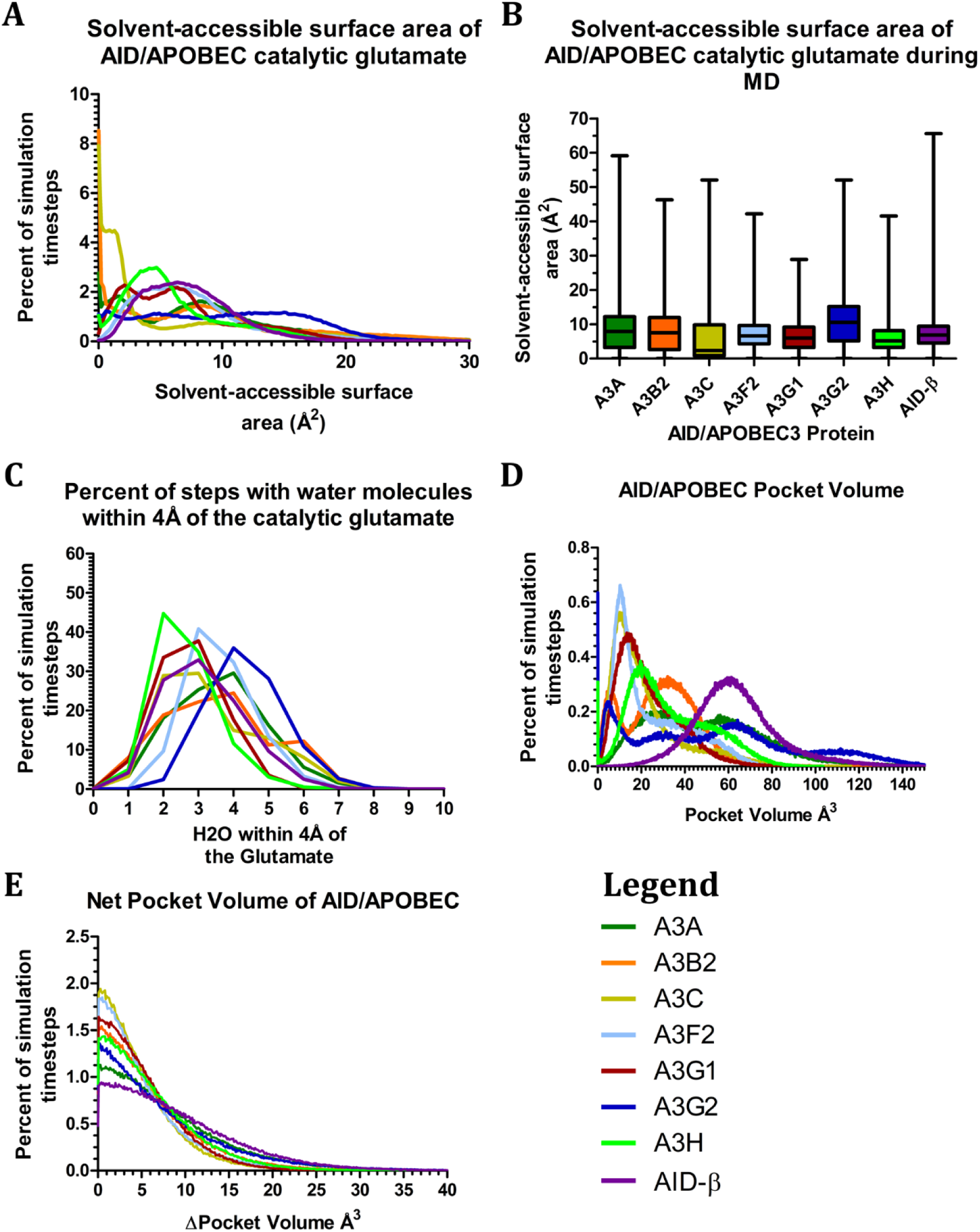
Pocket dynamics of AID/APOBEC3. Solvent-accessible surface area shown as a (A) histogram and (B) box and whisker plot of quartiles, and (C) number of water molecules within 4Å of the oxygens of AID/APOBEC3’s catalytic glutamates. (D) Pocket volume and (E) net pocket volume between timesteps of AID/APOBEC3 dependent on the number of steps.

Each of the A3Z subtypes and AID were different in SASA and water availability. Among the A3Z1 proteins, A3A and A3B-CD2 exhibited distinct SASA profiles. A3A had a non-symmetric trimodal distribution with peaks at 8.4 Å², 1.7 Å², and 0 Å², while A3B-CD2 showed a bimodal distribution with a major peak at 8.6 Å² and a minor one at 0 Å². A3G-CD2 had a more uniform, flat plateau from 0 Å² to 17.0 Å², lacking sharp peaks. Regarding water content in the catalytic pocket, all A3Z1s most commonly housed four water molecules. However, A3A and A3B-CD2 tended to spend more time with fewer than four waters, forming negatively-skewed distributions (Figure 3.C). A3G-CD2, in contrast, showed a positively-skewed distribution and often contained more than four waters. Interestingly, A3B-CD2 was the only A3Z1 subtype to display a non-symmetric bimodal water distribution with an additional peak at six waters in the pocket. The A3Z2 subtypes (A3C, A3F-CD2, and A3G-CD1) presented distinct patterns. A3C and A3F-CD2 showed positively-skewed unimodal SASA distributions, peaking at 0 Å² and 6.2 Å², respectively (Figure 3.A-B). A3G-CD1 exhibited a positively-skewed, non-symmetric bimodal distribution with peaks at 2.1 Å² and 6.4 Å². These three proteins typically maintained three waters in the catalytic pocket (Figure 3.C). A3G-CD1 again showed a negatively-skewed unimodal water distribution, spending more time with fewer than three waters, while A3C and A3F-CD2 exhibited positively-skewed distributions with more than three waters. The A3Z3 representative, A3H hap II, had a SASA peak at 4.7 Å² and most often accommodated two water molecules in its catalytic pocket (Figure 3.A-C). Both measurements formed positively-skewed distributions. AID itself presented a broad, unimodal SASA peak centered at 6.4 Å² and had a typical water content of three molecules, forming a positively-skewed SASA and a normally distribution water profile. Taken together, these data support our hypothesis that A3Z1 proteins have the most accessible catalytic environments, followed by a spectrum of intermediate values across A3Z2, A3Z3, and AID, with A3H hap II having the least water availability for hydrolysis.

Beyond SASA and water accessibility, we also analysed the catalytic pocket volume and its variability over short intervals (10 ps timesteps). Here, we predicted that the A3Z1 proteins would again exhibit the largest pocket volumes and greatest temporal fluctuations due to their enzymatic activity and structural dynamics with A3G-CD1 and AID displaying smaller and more rigid pockets. Across all proteins, we found at least some distribution density at 0 Å³; although, in most cases this occurred as a single point with minimal surrounding distribution. Within the A3Z1 group, A3A and A3B-CD2 showed non-symmetric bimodal distributions with A3A positively skewed and A3B-CD2 negatively-skewed (Figure 3.D). A3G-CD2 had a more complex positively-skewed multi-modal distribution with four distinct peaks. All three A3Z1 proteins shared a central volume peak around 26.6–32.1 Å³. Additional overlapping peaks were observed between A3A and A3G-CD2 at 56.3 Å³ and 64.3 Å³, respectively, and between A3B-CD2 and A3G-CD2 at lower volumes of 6.4 Å³ and 4.5 Å³. A3G-CD2 also displayed a smaller peak at 111.6 Å³. Maximum observed volumes were greatest in A3A at 210 Å³, followed by A3G-CD2 at 194.4 Å³, and lowest in A3B-CD2 at 134.9 Å³. In contrast, the A3Z2 proteins (A3C, A3F-CD2, and A3G-CD1) demonstrated consistent positively-skewed unimodal distributions with peaks at 10.1 Å³, 10.3 Å³, and 13.4 Å³, respectively. A3H hap II of the A3Z3 group exhibited a similar positively-skewed unimodal distribution peaking at 19.6 Å³. Among these, A3C had the highest maximum pocket volume at 116.3 Å³, followed by A3H hap II (103.5 Å³), A3G-CD1 (102.8 Å³), and A3F-CD2 (95.5 Å³). Interestingly, AID had a normally distributed pocket volume centered at 61.4 Å³ and a surprisingly high maximum volume of 172 Å³, overlapping in range with A3Z1 proteins.

To examine dynamic flexibility, we quantified the net volume change of the catalytic pocket over 10 ps intervals. Median pocket changes ranged from 3.63 Å³ to 7.38 Å³ across all AID/APOBECs (Figure 3.E). AID showed the highest median fluctuation followed by A3A, A3G-CD2, A3H hap II, A3B-CD2, A3G-CD1, A3F-CD2, and A3C. Maximum changes per 10 ps timestep revealed that A3G-CD2 had the greatest single interval fluctuation at 71.0 Å³. This was followed closely by A3A (68.63 Å³), AID (67.5 Å³), and A3B-CD2 (63.63 Å³). The A3H hap II and A3Z2 proteins trailed with moderate maximum changes with 57.25Å^3^ and 53.88-52.88Å^3^, respectively, while A3G-CD1 had the least at 41.25 Å³.

In conclusion, the A3Z1 proteins, particularly A3A and A3G-CD2, demonstrate the highest catalytic pocket accessibility, hydration, and volume fluctuation, correlating with their enzymatic activity. AID surprisingly mirrors some of this flexibility, though with less efficient catalysis. A3Z2 subtypes tend to have more restricted and less dynamic catalytic pocket volumes, while A3H hap II occupies an intermediate position among the AID/APOBEC family.

### 3.3. AID/APOBEC3 pocket states caused by catalytic residues and loop ℓ1 threonine/serine

Once we had determined the range of pocket volumes that the AID/APOBEC family proteins could adopt during simulation, the next critical step was to investigate how these volumetric variations impacted their functional states, specifically their ability to bind ligands. To assess this, representative 3D structural homology models were extracted at random time points, corresponding to a wide array of observed volumes. These models were then subjected to ligand docking simulations using dC as a probe. The goal was to determine whether the catalytic pocket was in a closed (ligand-inaccessible), open (ligand-accessible), or indeterminate state (where ligand accessibility varied across different models with similar volumes). We initially hypothesised that structural and functional differences between the A3Z subtypes would result in subtype-specific boundaries between these functional states. Moreover, given the historically low catalytic activity of AID and A3G-CD1, we posited that these enzymes would possess broader closed-volume ranges, reflecting a structural basis for their limited activity.

Indeed, all the catalytically active AID/APOBEC family members exhibited a closed pocket conformation at volumes from 0 Å³ to a maximum range that varied from 18 Å³ to 30 Å³ (Figure 4.A). At the lower end of this range were A3H hap II, AID, and A3C, while A3A exhibited the highest maximum volume at which closure still occurred. Notably, A3G-CD1 had the most extensive range of closed volumes, reaching up to 54 Å³, consistent with its known inactivity. The transition zones, defined as volume ranges where the pocket’s accessibility to the ligand was inconsistent (i.e., indeterminate state), varied widely among the proteins. AID and A3G-CD1 again stood out with the broadest ranges of indeterminate volumes at 48 Å³ and 45 Å³, respectively. In contrast, A3F-CD2 and A3G-CD2 had narrower indeterminate volume ranges of 28 Å³ and 30 Å³. The volumes associated with the open pocket conformations were equally diverse. These ranged from 39 Å³ in A3F-CD2 to 145 Å³ in A3A with other values including 45 Å³ for A3H hap II, 52 Å³ for A3C, 66 Å³ for A3B-CD2, and 135 Å³ for A3G-CD2. Notably, A3G-CD1 never adopted a conformation that was consistently open, and AID displayed a surprisingly broad range of open volumes (up to 89 Å³), which was unexpected given its low observed activity.

**Figure 4.**
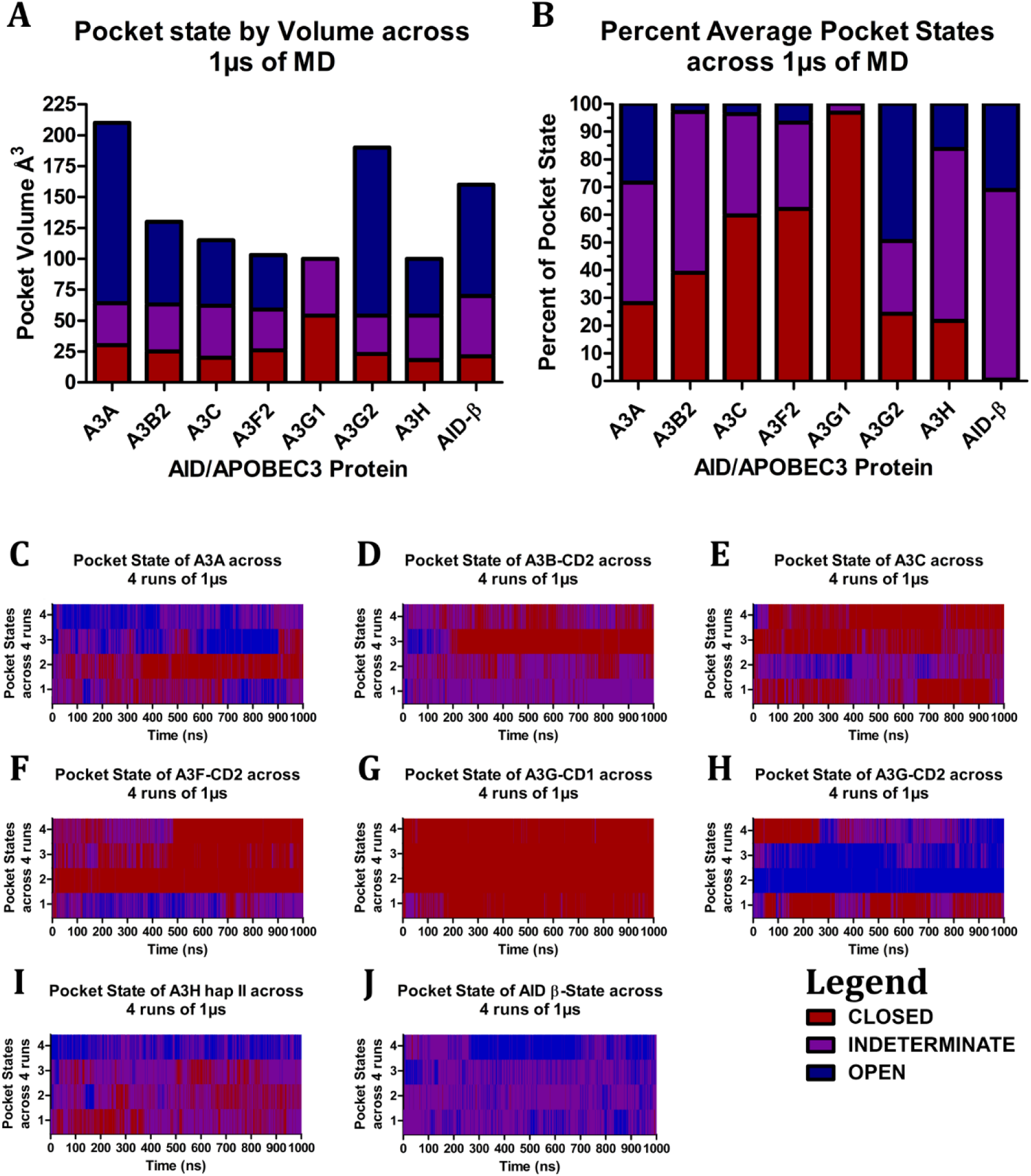
Pocket states of AID/APOBEC3. (A) Pocket state volume ranges and (B) pocket states percentages across multiple runs. Pocket states of 4 simulation runs of 1µs each for (C) A3A, (D) A3B-CD2, (E) A3C, (F) A3F-CD2, (G) A3G-CD1, (H) A3G-CD2, (I) A3H haplotype II, and (J) AID with the α7 in the β-state. The pocket states are registered as open (blue), indeterminate (purple), or closed (red) as determined by ability to dock substrate in the catalytic pocket.

**Figure 5.**
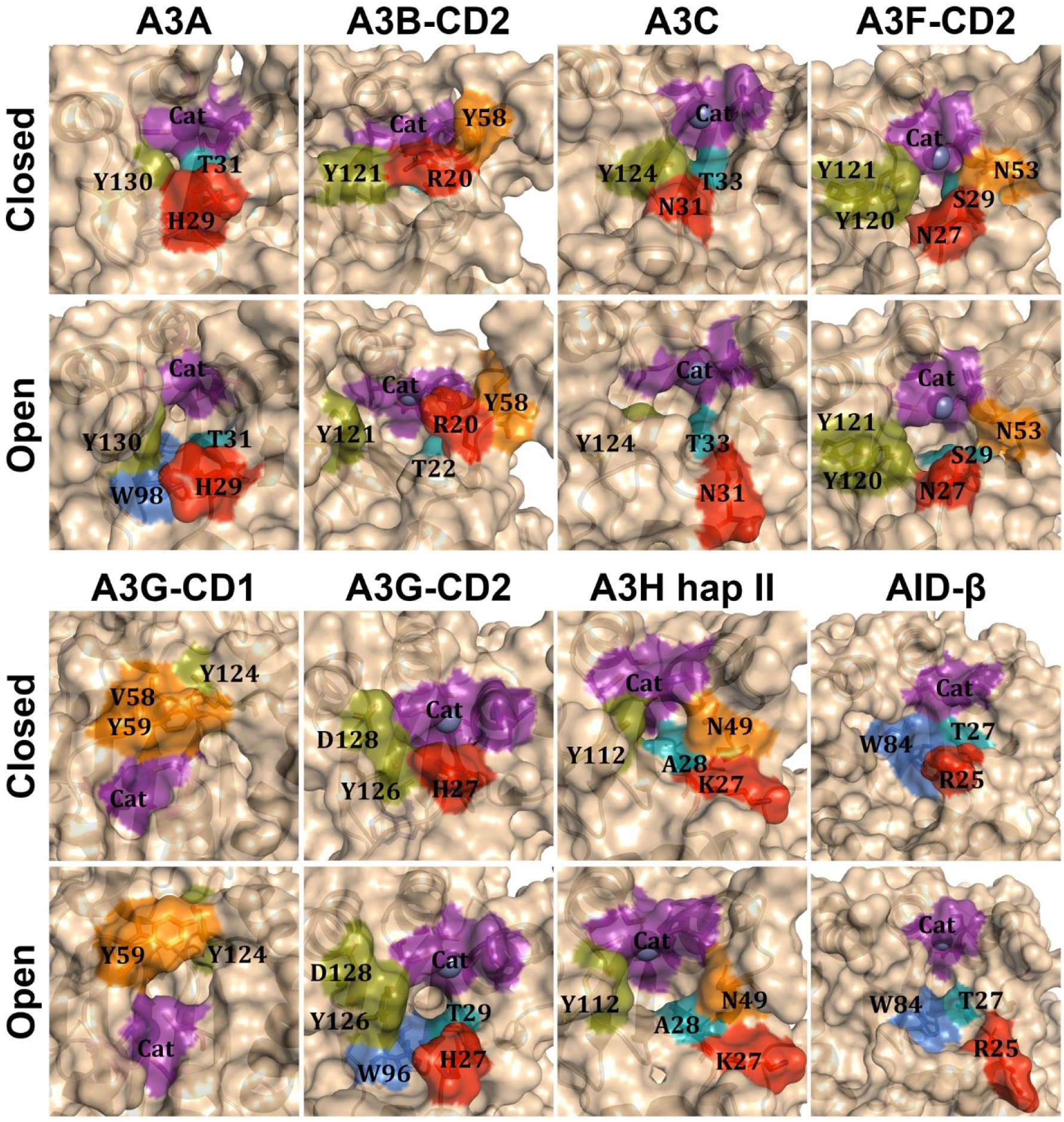
Structures of AID/APOBEC3 pocket states. AID/APOBEC3 proteins in both closed and open states shown as surface structure with select residues highlighted, and zinc as a sphere. The residues that are highlighted are the catalytic residues (purple), gatekeeper (red), and residues of the DNA-binding groove being loop ℓ1 (deepteal), loop ℓ3 (orange), loop ℓ5 (marine) and loop ℓ7 (deepolive).

To complement this volume-based analysis, we next examined how frequently each enzyme adopted the closed, open, or indeterminate pocket states throughout the course of its molecular dynamics simulation (Figure 4.B). Among the A3Z1s, A3A and A3B-CD2 primarily occupied the indeterminate state 43.6% and 58.1% of the time, respectively. A3G-CD2, in contrast, was mostly in the open state (49.5%). For A3A and A3G-CD2, time spent in the closed and open states was fairly balanced for the remainder of the simulation. However, A3B-CD2 was more likely to enter a closed conformation (39.0%). The A3Z2 subfamily displayed a different trend: they were closed most of the time. Specifically, A3C, A3F-CD2, and A3G-CD1 were in the closed state for 59.8%, 62.1%, and 96.9% of the simulation, respectively. A3C and A3F-CD2 also spent about a third of their simulation time in the indeterminate state, leaving minimal time in the open conformation. A3G-CD1, consistent with its maximum closed volume range, never entered the open state and only rarely adopted an indeterminate one. Among the A3Z3s, A3H hap II and AID were mostly in the indeterminate state with A3H hap II occupying this state for 62.1% and AID for 68.38% of the simulation. However, while A3H hap II split the remainder of the time fairly evenly between open and closed states, AID shifted nearly all remaining time into the open conformation. These data suggest that the A3Z1s and A3Z3s have evolved to more frequently adopt open or partially open (indeterminate) pocket conformations, potentially enhancing their catalytic readiness. Conversely, A3Z2s appear to be structurally biased toward closure which aligns with their generally lower enzymatic activities. The surprising finding was the large fraction of time that AID remained in an open or indeterminate state despite its poor catalytic efficiency.

To further explore the dynamics of state transitions and their temporal patterns, we analysed the trajectories of each protein to determine how frequently they moved between the open, closed, and indeterminate states over time. We hypothesised that certain proteins, particularly those that spent the most time in the indeterminate state, would exhibit more frequent transitions between the extremes of open and closed. In the A3Z1 group, A3A demonstrated continual switching between all three states in most simulations, except for one trajectory where it remained predominantly closed (Figures 4.C and S1–S4). A3B-CD2, by contrast, mainly cycled between the indeterminate and closed states with the indeterminate state being dominant (Figures 4.D and S5–S8). In one notable trajectory, A3B-CD2 alternated between open and indeterminate states for 190 ns before adopting a sustained closed conformation. A3G-CD2 showed high variability consistent with its broad volume distribution. In two simulations, it remained open with only brief detours into the indeterminate state or closure. Another trajectory showed extensive switching, and in a fourth run, it was predominantly closed, entered indeterminate and even open conformations sporadically, then returned to closure (Figures 4.H and S21–S24). Among A3Z2s, A3C remained mostly closed in all but one trajectory, where it entered the indeterminate state for short intervals (∼50 ns), and another where it alternated between all three states (Figures 4.E and S9–S12). A3F-CD2 behaved similarly; although one simulation featured intermittent transitions between open and indeterminate states (Figures 4.F and S13–S16). A3G-CD1 never adopted an open conformation and only entered the indeterminate state briefly in two runs (Figures 4.G and S17–S20). In one of these, the state lasted under 2 ns, and in the other, it persisted for ∼35 ns, briefly recurring after 160 ns. A3H hap II (A3Z3) mostly alternated between indeterminate and closed states, resembling the behavior of A3B-CD2 (Figure 4.I and S25–S28). However, in one simulation, it remained open for the entire trajectory, occasionally shifting to the indeterminate state in short bursts of 5–10 ns. AID was predominantly indeterminate but frequently switched to the open state for short periods, except in one run where it remained open for 400 ns with only brief (10–20 ns) excursions into the indeterminate state (Figure 4.J and S29–S32). These patterns reinforce the conclusion that A3Z2s tend to remain structurally closed with rare conformational transitions to states that might permit activity. A3G-CD1 stands out as the only protein we tested to be incapable of opening. Conversely, A3B-CD2, A3H hap II, and AID favour the indeterminate state and display dynamic switching behavior, but the direction of those transitions differs: A3B-CD2 and A3H hap II switch toward closure, while AID alternates into openness. A3A stands out for its rapid and frequent cycling between all three states, whereas A3G-CD2 enters all three but tends to stabilise in one once adopted.

To identify the molecular determinants of these conformational dynamics, we investigated which residues most directly contributed to pocket state changes. We hypothesised that catalytic and gatekeeper residues would be primary contributors. Consistently across all proteins, closure of the pocket was marked by inward movement of the catalytic residues, drawing them closer to the central β-sheet (Figure 5). In proteins like AID, A3A, and A3B-CD2, where ℓ1 includes arginine, histidine, and arginine as the gatekeeper, respectively, the repositioning of this residue over the catalytic pocket was frequently associated with closure. For A3F-CD2 and A3G-CD2 (with gatekeeper asparagine and histidine, respectively), this movement was occasionally observed but not required for closure. In A3C, A3G-CD1, and A3H hap II (asparagine, asparagine, and lysine gatekeepers, respectively), gatekeeper residues did not significantly obstruct the pocket. Another residue that consistently influenced closure was the terminal threonine of ℓ1 in A3A, A3B-CD2, A3C, A3G-CD2, and AID, which often participated via its sidechain hydroxyl. In A3F-CD2, a homologous serine performed the same role. However, this interaction was absent in A3G-CD1 (despite also containing a threonine) and in A3H hap II (where alanine replaced the threonine). Additional influential residues were found in ℓ3, ℓ5, and ℓ7. Loop ℓ3 residues such as N240 and Y250 (A3B-CD2), V58 and Y59 (A3G-CD1), and N247 (A3F-CD2) affected pocket dynamics. Aromatic side chains in ℓ5, including W281, W287, F285 (A3B-CD2), W284 (A3F-CD2), W292 (A3G-CD2), and W84 (AID), modulated pocket walls. Less frequent effects came from W82 (A3H hap II) and W98 (A3A). Hydroxyl-bearing residues like S293 (A3G-CD2) and S85 (AID) occasionally influenced volume. Loop ℓ7 also contributed to pocket closures via residues such as Y112 (A3H hap II), Y124 (A3C), and Y314/Y315 (A3F-CD2). Additional contributors include Y130 (A3A), Y313 (A3B-CD2), Y124 (A3G-CD1), and Y322/D324 (A3G-CD2). Notably, A3G-CD1’s unique ℓ3 geometry causes bound cytidine to orient away from the catalytic proton shuttle and Zn-OH, likely making catalysis infeasible. Together, these observations suggest that five residues are most critical in determining pocket state: the four catalytic residues and the ℓ1 threonine/serine. Secondary roles are played by residues from ℓ3, ℓ5, and ℓ7, while gatekeeper residues are functionally significant in only a subset of APOBEC proteins.

### 3.4. AID α7’s α-state more stable than β-state but can decrease pocket volume

While analysing the conformational behavior of AID’s catalytic pocket, we observed that AID consistently adopted an indeterminate conformation far more frequently than anticipated. Surprisingly, it occupied a closed conformation significantly less often and an open conformation significantly more often than expected. In our initial simulation set, all of the 3D structural homology models were constructed to begin in an open conformation, except A3G-CD1. In AID models, the α7 helix began in contact with the central β-sheet and α6 helix (Figure 6.A-B). We termed this arrangement the β-state. Across all simulations, α7 consistently moved away from its starting β-state position, indicating a lack of stability in this conformation. To investigate the stability and functional relevance of different α7 conformations, we developed two additional simulation sets. The first set consisted of AID models initialised in a closed catalytic pocket conformation to evaluate whether starting in this state increased the likelihood of maintaining closure throughout the trajectory. The second set involved an alternate α7 orientation, termed the α-state, in which α7 interacts with residues from the assistant patch on α6 (specifically R171, R174, R177, and R178) and ℓ7. This was designed to ascertain whether the α-state provides greater structural stability than the β-state.

**Figure 6.**
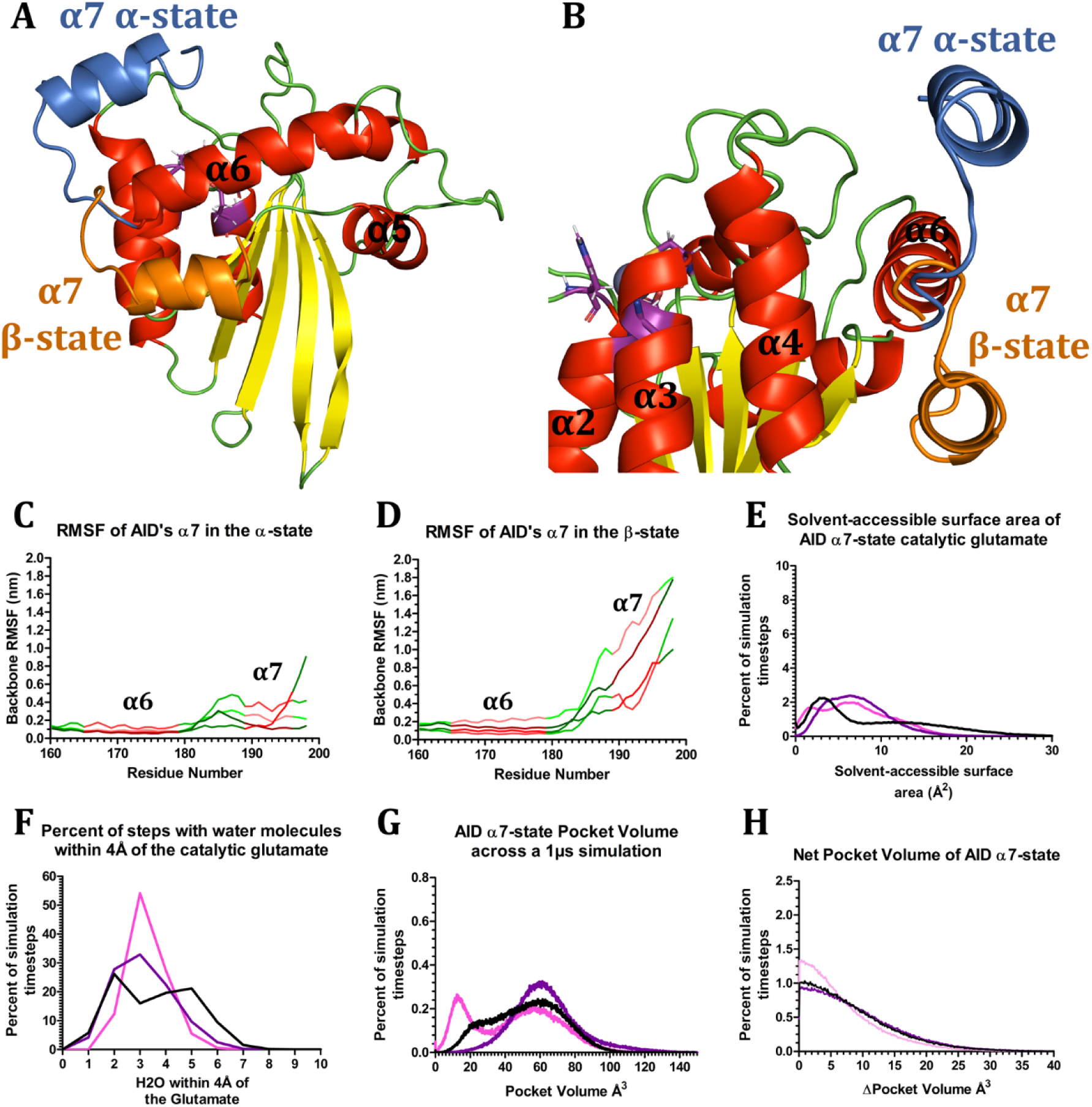
AID RMSF and pocket dynamics across α7. (A-B) Structure of AID from 2 angles coloured by secondary structure with the catalytic residues in purple. The loop ℓ11, α7 and the C-terminal loop are highlighted by the conformation as α-state (skyblue) and β-state (orange). RMSF of AID’s α7 dependent on the (C) α-state or (D) β-state coloured by secondary structure where α-helices (red) and loops (green) are shaded for different runs. (E) Solvent-accessible surface area and (F) number of water molecules within 4Å of the oxygens of AID/APOBEC3’s catalytic glutamates. (G) Pocket volume and (H) net pocket volume between steps of AID/APOBEC3 dependent on the number of steps. (E-F) Graphs of AID are coloured by their initial states with α-state (pink), β-state (purple), and β-state initially with a closed catalytic pocket (black).

When comparing the RMSF values among the different conformations, we found that the regions with the highest flexibility were generally consistent across simulations. However, a key distinction emerged in the behavior of the α7 helix itself (Figure 6.C-D). In the β-state, α7 exhibited elevated fluctuations and frequently drifted away from its starting position to interact with various other regions of the protein. In contrast, α7 in the α-state consistently remained stably positioned between α6 and ℓ7. These results suggest that the α-state, which overlays AID’s assistant patch, represents a more structurally stable configuration than the β-state. Furthermore, the apparent dynamic interplay between these α7 conformations likely modulates DNA binding, particularly in contexts involving structured versus unstructured DNA substrates.

We next sought to understand how these α7 conformations influence the structural and functional characteristics of the catalytic pocket. We examined key parameters such as the SASA of the proton shuttle, the water occupancy within the pocket, and the overall pocket volume across the different states. For β-state simulations starting with an open pocket, the SASA distribution of the proton shuttle followed a positively-skewed unimodal pattern peaking at 6.4 Å², as previously described (see section 1.4.2). When simulations began in a closed conformation, however, this peak shifted to 2.8 Å². Intriguingly, the α-state simulations produced a positively-skewed, non-symmetric bimodal distribution with peaks at 1.9 Å² and 5.8 Å² (Figure 6.E). Regarding water occupancy in the catalytic pocket, both β-state and α-state models initiated in the open conformation showed positively-skewed unimodal distributions, peaking at approximately three water molecules (Figure 6.F). Notably, this peak was more pronounced in the α-state. In contrast, models that began in the closed conformation showed a non-symmetric bimodal distribution with peaks at two and five waters. Pocket volume distributions provided further insights: β-state simulations initiated from an open conformation yielded a unimodal peak at 61.4 Å³ with a normal distribution, whereas those started in the closed conformation peaked at 58.8 Å³ with a negatively-skewed distribution (Figure 6.G). The α-state simulations produced a non-symmetric bimodal distribution with peaks at 57.0 Å³ and 12.4 Å³. While the net volume change was comparable between the β-state simulations that began in open or closed conformations, the α-state simulations consistently showed lower net changes, aligning more closely with results from A3H hap II and A3G-CD2 (Figure 6.H).

To contextualise these structural fluctuations, we examined how pocket conformational states (open, closed, or indeterminate) varied across simulation sets. While the specific volumes associated with each state remained constant regardless of α7 positioning, the proportion of time spent in each state varied markedly (Figure 7.A-B). The indeterminate state dominated across all simulations, occurring 68.38% and 73.05% of the time in the β-state initially open and closed sets, respectively, and 58.05% of the time in the α-state. Notably, AID in the β-state occupied the open state 31.04% of the time, compared to 21.07% and 18.30% in the initially closed β-state and α-state sets, respectively. Strikingly, while the β-state model rarely adopted the closed state (0.58% of the time), the initially closed conformation achieved this state 5.87% of the time. The α-state conformation achieved closure even more frequently, spending 23.65% of the simulation time in the closed conformation.

**Figure 7.**
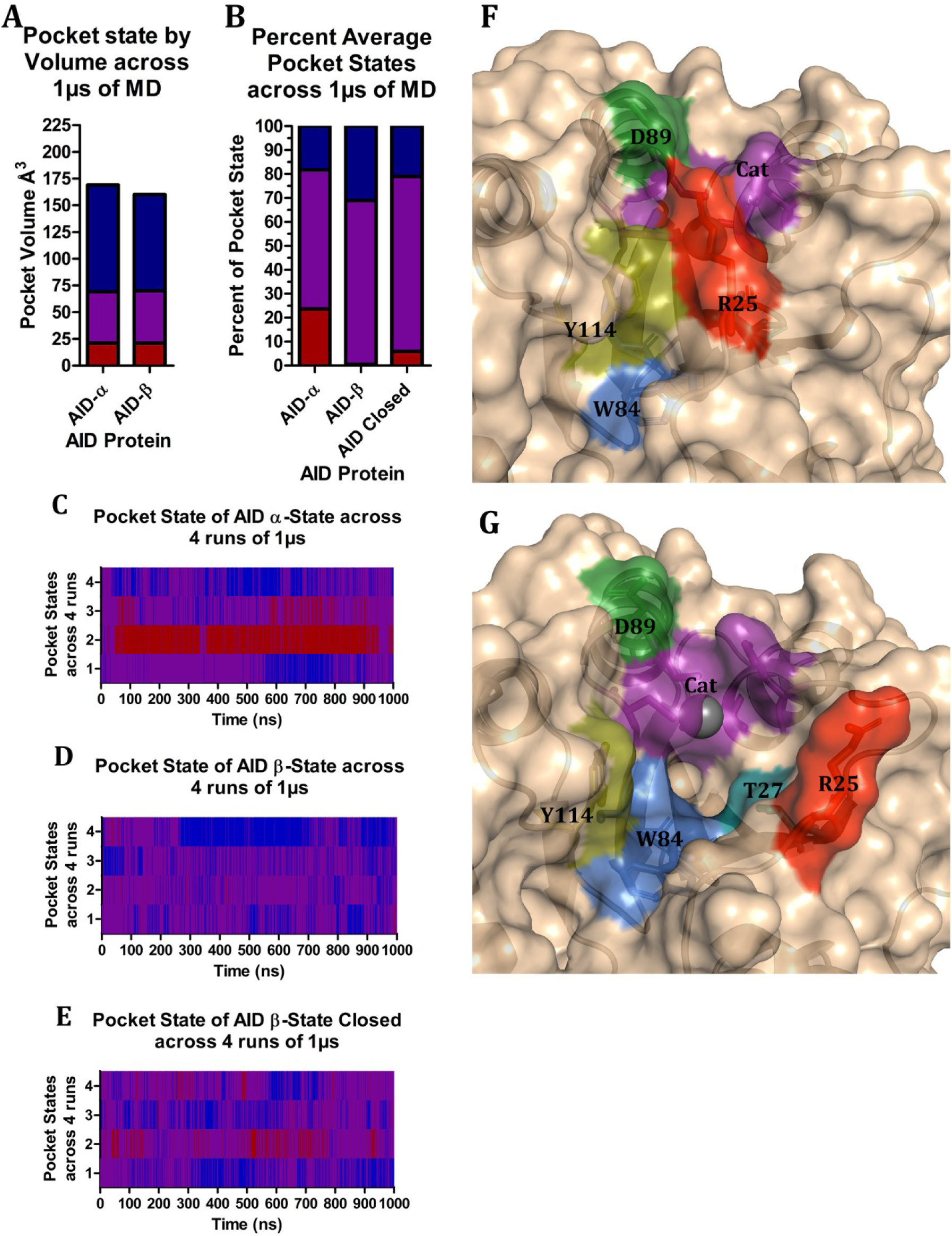
AID pocket states across α7. (A) Pocket state volume ranges and (B) pocket state percentages across multiple runs. Pocket states of 4 simulation runs of 1μs each for (C) AID in the α-state, (D) AID in the β-state, and (E) AID the β-state beginning in a closed conformation. The pocket states are registered as open (open), indeterminate (purple), or closed (red). The structure of AID α-state shown in (F) closed and (G) open conformations with cartoon, surface structures, and zinc as a sphere. Key residues are highlighted and shown as sticks such as the catalytic residues (purple), the R25 gatekeeper (red), T27 (deepteal), W84 and S85 (marine), Y114 (deepolive) and D89 (forest).

Inspection of individual simulation trajectories provided additional resolution. In the closed set, most simulations remained largely indeterminate with transient shifts to the open state for 10-20 ns, although one simulation remained predominantly open for ∼400 ns (Figures 7.D-E, S29–S32, and S37–D40). Two runs from the initially closed set were more likely to re-enter the closed state. Similarly, in the α-state simulations, one trajectory spent the majority of its time in the closed state, indicating an increased tendency toward closure compared to other conformations, which also were closed more often than the β-state (Figures 7.C and S33–S36).

Importantly, across all simulations, the same set of amino acids drove transitions between pocket states (Figure 7.F-G). These residues include the catalytic triad, the R25 gatekeeper, and the T27 hydroxyl group, with minor contributions from W84 and S85 of ℓ5. Notably, in the predominantly closed α-state simulation, α7 interacts with ℓ1, prompting it to shift position and enable R25 to cap the catalytic pocket, forming stable interactions with D89 of α4 and Y114 of ℓ7. These interactions appear to facilitate pocket closure in the α-state.

In summary, despite variability in the initial α7 positioning and starting pocket states, AID simulations exhibit consistent dynamics in the following key structural features: secondary structures, α7 motion, proton shuttle exposure, catalytic pocket volume, and gating residues. However, the α-state consistently promotes lower average pocket volumes and higher closure frequency than the β-state, and its greater structural stability may indicate that the α-state represents AID’s native conformation.

### 3.5. AID α7 truncations modify the assistant patch and substrate recognition loop ℓ7

To better understand the structural role of AID’s α7 helix, we simulated a series of truncations using 3D structural homology models based on both α-state and β-state conformations. We ran wild-type models in replicate alongside three truncations (terminating at residues 181, 183, and 188) derived from the α-state model (Figure 8.A). These simulations were designed to assess α7 stability and its interactions with surrounding structural elements. The wild-type α-state maintained a stable conformation throughout the simulations with α7 consistently interacting with α6 and ℓ7 (Figure 8.B). In contrast, the wild-type β-state rapidly became positionally unstable, losing its initial contacts after 252–300 ns. In one replicate, α7 migrated to interact with α4 and remained there, while in another, it cycled between contacts with α4, α6, and ℓ7, occasionally drifting into free solution before returning to the β-sheet region. In a third replicate, the β-state adopted a conformation resembling the α-state, although this mimic was less stable and did not fully insert into the α6/ℓ7 groove. Notably, this pseudo-α-state configuration disintegrated after 608 ns, suggesting that such mimicry lacks the stabilising interactions characteristic of true α-state binding.

**Figure 8.**
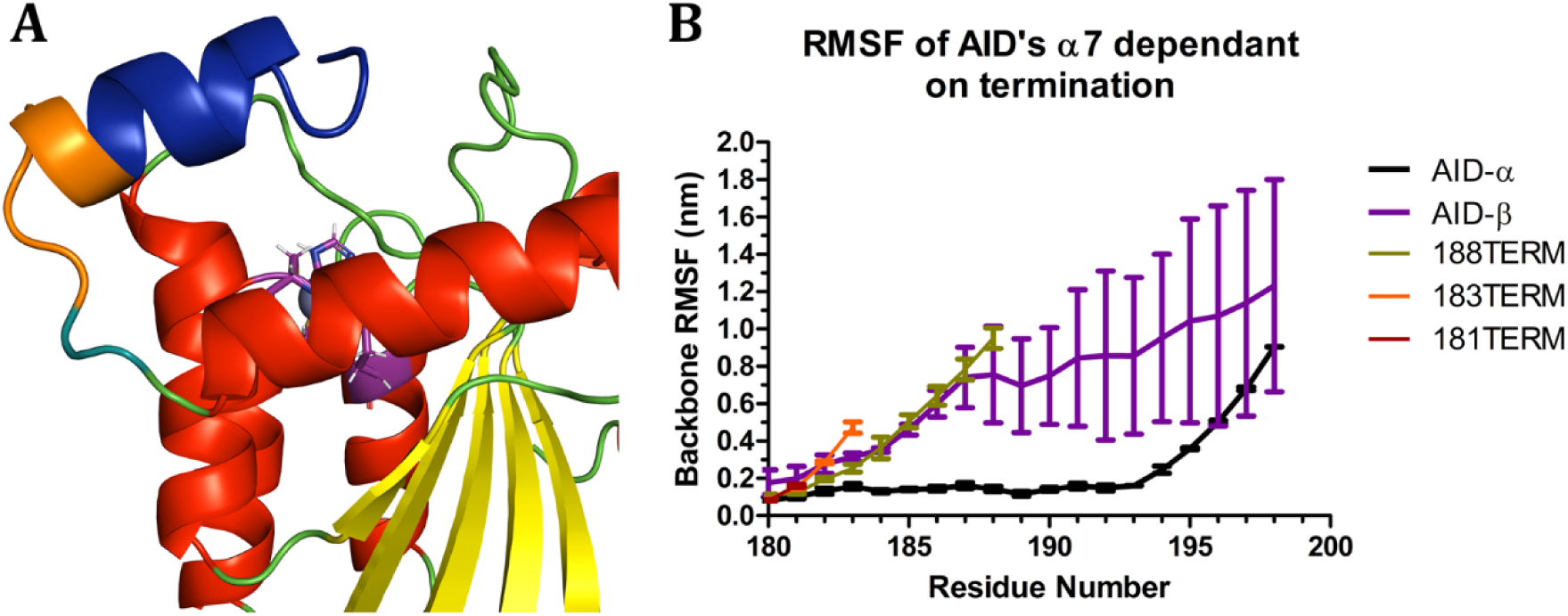
RMSF of AID α7 truncations. (A) Structure of AID coloured by secondary structure with the catalytic residues in purple, and highlighted regions that are terminated at residues 188 (blue), 183 (orange), or 181 (deepteal). (B) RMSF of multiple runs represented by error bars of AID proteins in the α-state, β-state, and the three terminations.

To further dissect α7’s role, we simulated truncations reported by Mu et al., 2012, focusing on the functional consequences of losing α7 segments [80]. We hypothesised that termination at residue 183 or 188 would disrupt the α6/ℓ7 interface, while truncation at 181 would destabilise α6 itself due to the proximity of the cut site. Indeed, neither the 181 nor 183 truncations maintained stable interactions with α6 or ℓ7. The 188 truncation initially contacted the α6/ℓ7 groove but rapidly lost stable positioning, leading to α7 secondary structure loss. However, the assistant patch consistently salt-bridged with D187 and D188. Truncations at 183 and 188 behaved similarly to the β-state, showing increased positional instability likely due to insufficient α6 support. This instability may explain the reported increase in activity of these truncations: the loss of a stable α-state α7 could reduce steric hindrance at the assistant patch, facilitating binding and promoting an open catalytic pocket conformation. Interestingly, the 181 truncation did not destabilise α6 as predicted, despite its location, and maintained moderate stability throughout. This finding suggests that factors beyond local structural disruption may contribute to the lowered activity observed in this mutant, though our current model does not fully explain the mechanism.

### 3.6. AID assistant patch mutations R171Y and R178D affect stability of α7

We next investigated how the assistant patch influences α7 stability. The assistant patch residues (R171, R174, R177, and R178) surround α6 in a configuration that enables binding to various conformations of α7, excluding fully solvent-exposed states (Figure 9.A–D). To assess the effects of disrupting these interactions, we introduced four point mutations: R174E and R178D, which may repel negatively charged α7 residues depending on initial conformation, and R171Y and R177A, which alter hydrophobic interactions. Simulations showed that R171Y significantly destabilised ℓ11 and α7 in both the α- and β-states (Figure 9.E, I). In the α-state, R171Y caused α7 to dislodge from its native position, leading to free solution behavior and transient interactions with α3 and α4. This was accompanied by either full unraveling or C-terminal unraveling of α7 within 130 ns. In the β-state, while RMSF values remained similar to wild-type, α7 again disintegrated, falling apart between 220–250 ns due to internal interactions with the introduced tyrosine. R174E exhibited a more nuanced effect. It had minimal impact on the β-state, which appeared slightly more stable (Figure 9.J), but in the α-state, R174E destabilised the C-terminal half of α7 almost immediately (<10 ns) while maintaining its position (Figure 9.F). The α-state mutation promoted salt bridge formation with α7’s R190 and R194, anchoring the disordered segment even as secondary structure was lost. The R177A mutation was less disruptive. In the α-state, only minor destabilisation was observed at the N-terminal end of α7, which began interacting with α6 (Figure 9.G). In the β-state, however, R177A led to gradual α7 unraveling and reorientation toward α4, occurring between 64–280 ns (Figure 9.K). The R178D mutation had the most dramatic effects. In the α-state, R178D caused full or partial detachment of α7, either shifting to interact with α3 and α4 or exhibiting N-terminal unraveling due to electrostatic repulsion from D187 and D188 (Figure 9.H). In the β-state, α7 adopted a novel conformation stabilised through new salt bridges: R190 and R194 were attracted to the introduced D178, while D187 and D188 were repelled and instead contacted α4 (Figure 9.L). Notably, the resulting orientation placed α7’s C-terminal end closer to α6, contrasting with R171Y, where the C-terminal end moved furthest from α6.

**Figure 9.**
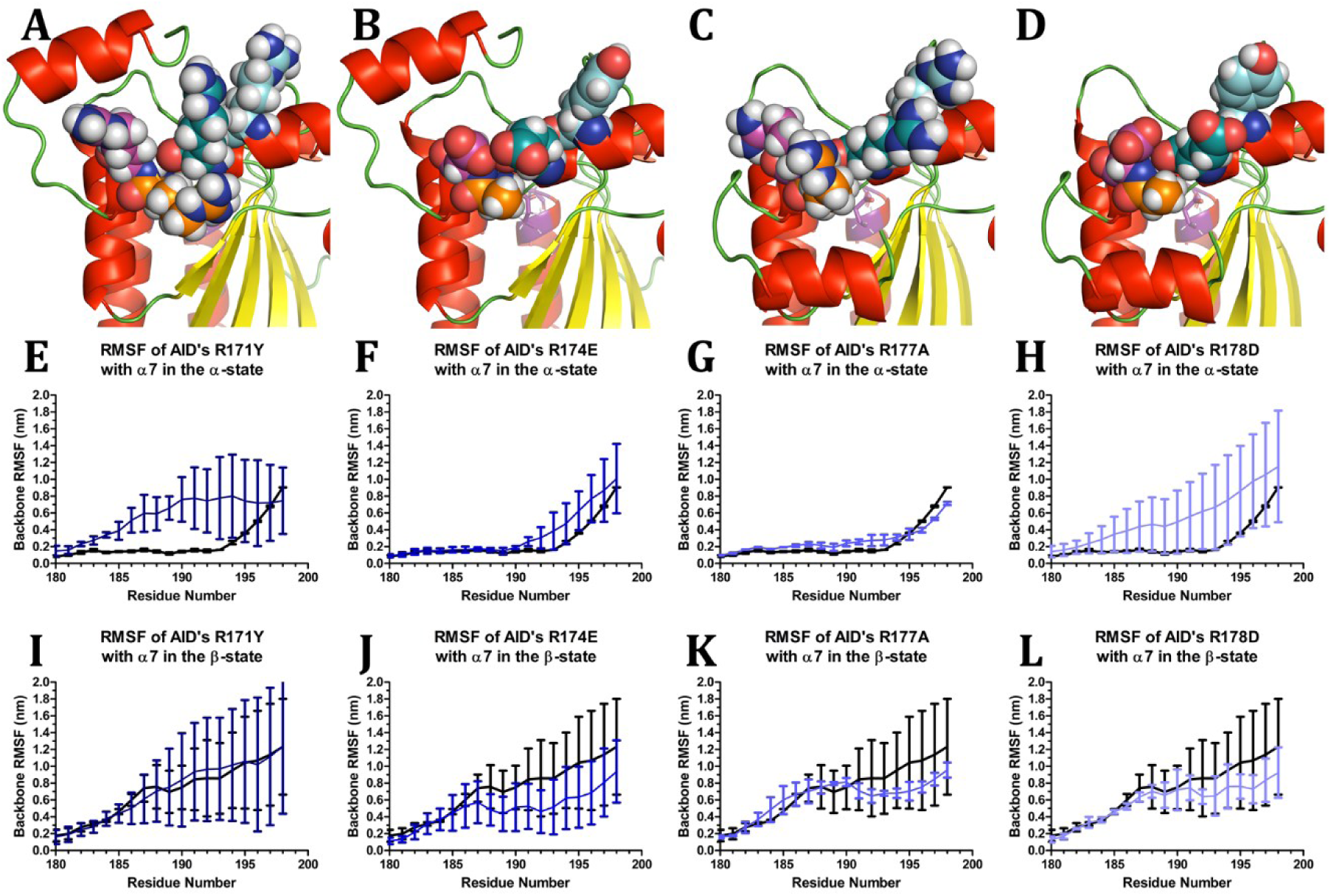
RMSF of AID α7 with α6 “assistant patch” mutations. Secondary structure of AID coloured by secondary structure shown in the (A-B) α-state, or (C-D) β-state. Residues are shown spheres for R171Y (cyan), R174E (deepteal), R177A (orange), and R178D (magenta), where (A,C) are shown as WT and (B,D) is shown in the modified form. RMSF graphs show (E-H) modifications of the α-state, and (I-L) modifications of the β-state, both compared to WT (black) where multiple runs are represented by error bars.

In summary, mutations like R171Y and R178D significantly destabilise the α7’s α-state, while R174E and R177A primarily affect secondary structure or specific structural contacts without displacing α7 (Figure 9.E–H). In the β-state, only R178D induces a fundamentally new conformation, suggesting R178D and, to a lesser extent, R171Y may better expose assistant patch regions compared to their α6 mutant counterparts (Figure 9.I-L).

### 3.7. AID α7 mutations R190D and R194A are capable of stabilising the β-state

To explore how α7 mutations influence their structural stability and positioning in both α- and β-states, we simulated mutations of key residues on α7: R190, F193, and R194 (Figure 10.A–D). In the α-state, R190 and R194 interact with ℓ7, while F193 contacts both ℓ7 and α6. In the β-state, R190 binds ℓ11 and α6, R194 is solvent-exposed, and F193 interacts solely with α6. Based on these roles, R190A and R194A were expected to disrupt salt bridges, R190D may alter charge-based interactions, and F193A may reduce stabilising planar-hydrophobic contacts. In simulations, R190A did not displace α7 from its α-state position, though its C-terminal half unravelled almost immediately (<10 ns), subsequently binding to ℓ7 residues Y114 and F115 (Figure 10.E). In the β-state, R190A had minimal effect, resembling the wild-type (Figure 10.I). To further probe R190’s role, we tested R190D, a charge-reversal mutation. In the α-state, R190D maintained binding to α6 and ℓ7 but formed weaker interactions with ℓ7 due to electrostatic repulsion from E117 and D118 (Figure 10.F). In contrast, in the β-state, R190D increased α7 stability, apparently forming new salt bridges with R177 and R178 (Figure 10.J). F193A behaved similarly to wild-type in terms of interaction sites in both states but consistently induced partial unraveling of α7 (Figure 10.G, K). In the α-state, this unraveling occurred early (∼5 ns), and loss of interactions with ℓ7’s Y114 and F115 appeared to destabilise residues 184–194. In the β-state, unraveling was slower (∼260 ns) but followed a similar trend. R194A did not markedly affect α7’s overall positioning in either state. Interestingly, in the α-state, residues 195–198 were slightly more stable than wild-type (Figure 10.H). In the β-state, however, α7 did not maintain its original position. Instead, it relocated to interact along a solvent-exposed face of α6, away from ℓ7 and the β-sheet. Here, R194A facilitated interactions between D191 and the R-group of R178 via its guanidinium group (Figure 10.L).

**Figure 10.**
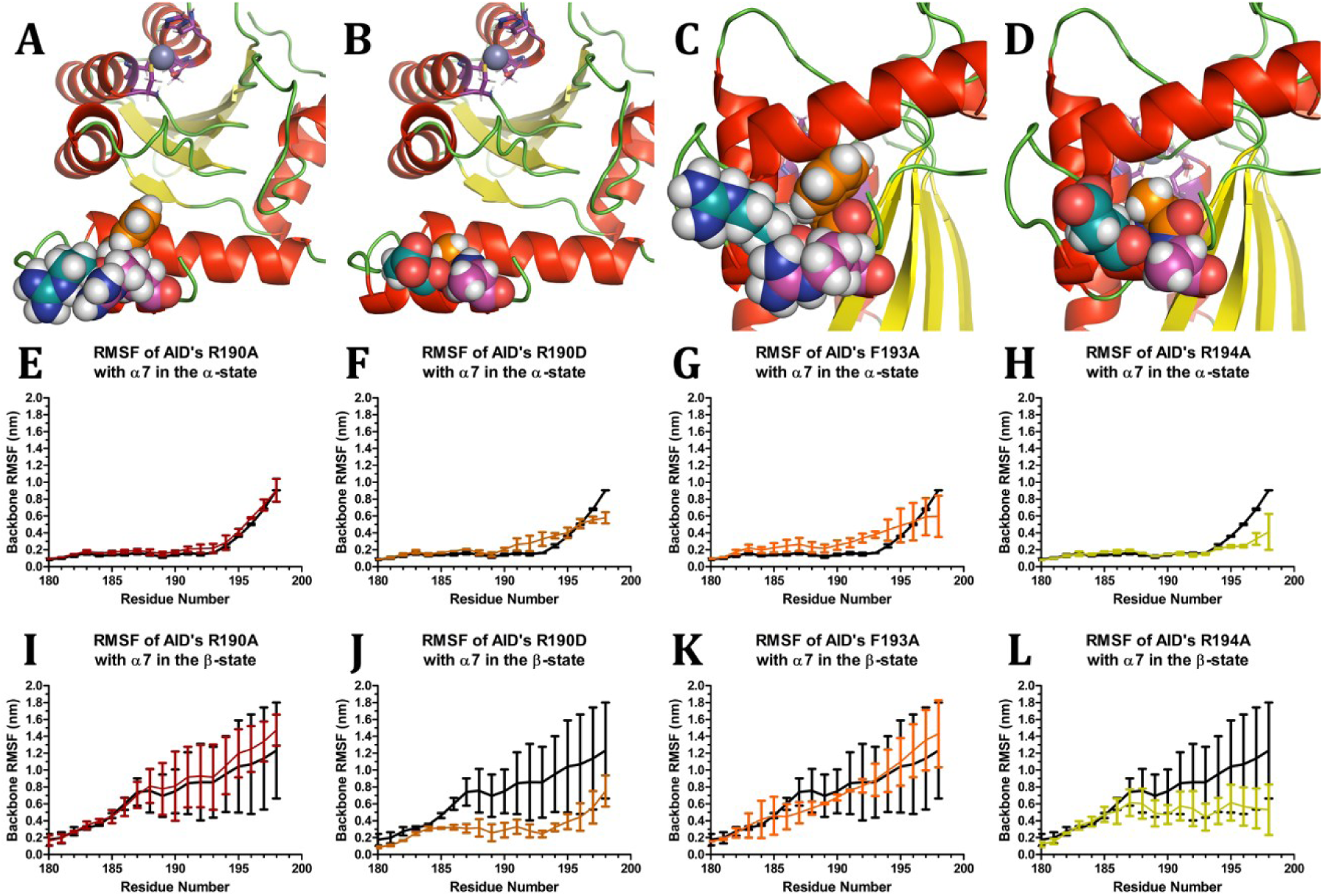
RMSF of AID α7 with mutations in α7. Secondary structure of AID coloured by secondary structure shown in the (A-B) α-state, or (C-D) β-state. Residues are shown spheres for R190D (deepteal), F193A (orange), and R194A (magenta), where (A,C) are shown as WT and (B,D) is shown in the modified form. RMSF graphs show (E-H) modifications of the α-state, and (I-L) modifications of the β-state, both compared to WT (black) where multiple runs are represented by error bars.

Overall, α7 in the α-state remained correctly positioned across all mutations, although R190A, R190D, and F193A disrupted specific stabilising interactions, leading to varying degrees of structural destabilisation (Figure 10.E–H). In the β-state, R190D notably increased conformational stability, and R194A relocated α7 and exposed the assistant patch, while R190A and F193A behaved more like wild-type, except for F193A’s tendency to induce unraveling (Figure 10.I–L). This suggests that although these mutations alter specific interactions, they do not destabilise the α-state; R190D and R194A can stabilise β-like conformations and nearby alternatives that expose functional surfaces.

### 3.8. AID mutations and PTMs of regions surrounding α7 affect binding but do not disrupt the α-state or completely stabilise β-state

We next examined a set of mutations and PTMs occurring outside of α7 but capable of influencing its structure and interactions. Specifically, we focused on residues H130, T140, Y184, and L198, which are located in α4, ℓ9, ℓ 11, and at the C-terminus, respectively (Figure 11.A–D). These positions were selected based on their proximity to α7 and their established or predicted roles in stabilizing regional structural elements. In the WT structure, H130 sits at a critical junction between β-strand 5, α6, and ℓ11, and can also contact α7 in the β-state. The H130P mutation eliminates a side chain capable of hydrogen bonding and removes a positive charge, likely disrupting local packing and flexibility. T140 interacts with α7 and the C-terminal residue in the β-state, while Y184 is involved with α4, α6, and α 7, as well as ℓ7 and 11, and β-strand 5. Phosphorylation of T140 (T140-p) and Y184 (Y184-p) alters their charge properties and interaction profiles, introducing repulsive or attractive effects that can influence the C-terminal residue, and ℓ11 or the α7 orientation, respectively. L198, the C-terminal residue, interacts with ℓ1, ℓ7, ℓ9, the β-sheet, α5, and α6. The L198S mutation introduces a polar side chain capable of hydrogen bonding, potentially affecting how α7 and the C-terminus engage with their surroundings.

**Figure 11.**
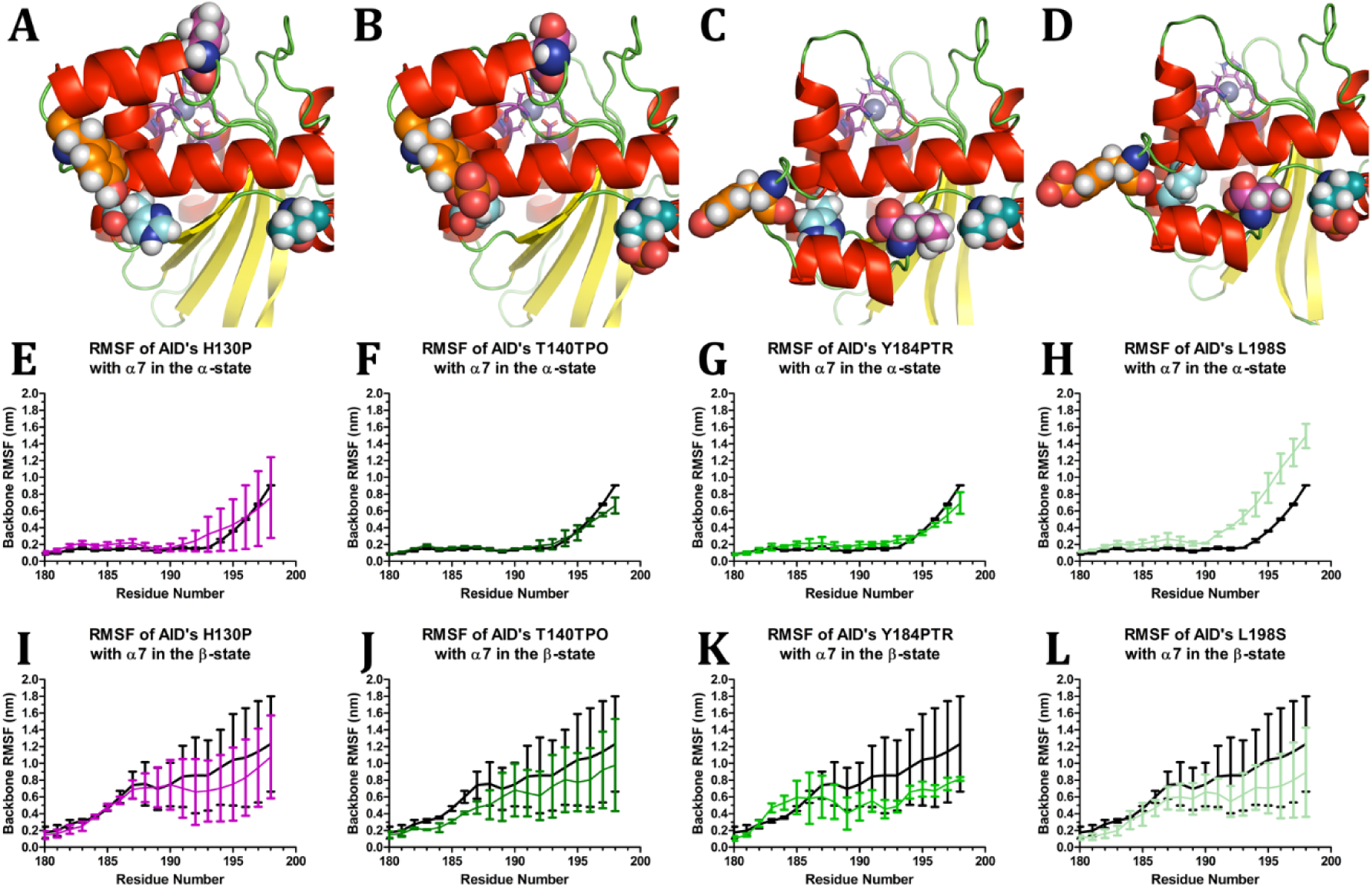
RMSF of AID α7 with mutations and PTMs across alternate regions. Secondary structure of AID coloured by secondary structure shown in the (A-B) α-state, or (C-D) β-state. Residues are shown spheres for H130P (cyan), T140TPO (deepteal), Y184PTR (orange), and L198S (magenta), where (A,C) are shown as WT and (B,D) is shown in the modified form. RMSF graphs show (E-H) modifications of the α-state, and (I-L) modifications of the β-state, both compared to WT (black) where multiple runs are represented by error bars.

H130P exhibited fluctuations similar to WT in both α and β conformations (Figure 11.E,I). However, in the α-state, the C-terminal half of α7 unravelled, likely due to weakened interactions with α6 and ℓ11. In the β-state, which was similar to WT, two trajectories diverged: in one, α7 bound to α4 and α6 via H130P-T195 or H130P-F193 interactions; in the other, α7 instead interacted with ℓ7 and α6, but α6 and α7 were positioned perpendicularly rather than in parallel. For T140-p, no change was observed in the α-state (Figure 11.F,J). However, in the β-state, the phosphate group repelled the C-terminal carboxylate, shifting α7 to bind α6 on its most solvent-exposed surface. Y184-p had minimal effect on the α7 fluctuations in the α-state, but in the β-state, it enhanced overall structural stability (Figure 11.G,K). In the α-state, although α7’s position is retained throughout, Y184-p is inserted between α6 and α7, attracted to assistant patch arginines which shifted α7 toward ℓ7 and caused the α7’s C-terminal half to dissolve. In the β-state, Y184-p increased fluctuations in residues 183–184 while stabilizing other interactions. The phosphorylated tyrosine often bound α4 or α6 instead of α7, resulting in α7 interacting more with the solvent-facing side of α6. L198S significantly reduced α7 stability in the α-state (Figure 11.H). The C-terminal portion of α7 unravelled and frequently shifted between its usual position and the solvent-exposed face of α6. In the β-state, stability was somewhat increased compared to WT but within expected limits (Figure 11.L). Here, α7 stabilised between α4 and α6’s solvent-facing surface. Notably, even when free in solution, the L198S residue often formed salt bridges with nearby regions, contributing to intermittent stabilization.

Overall, L198S caused the greatest destabilisation of α7 in the α-state and H130P disrupting α7’s C-terminal half, while T140-p and Y184-p primarily influenced structural interactions without disrupting α7 folding (Figure 11.E-H). The β-state remained unstable across all modifications, though Y184-p and L198S promoted partial stabilization via contacts with α4 and α6 (Figure 11.I–L). T140-p induced repulsion with the C-terminal residue, altering α7 binding, and H130P changed α7’s interaction partners. Collectively, these results indicate that while these residues modulate α7’s structural and interaction profile, none are sufficient to completely destabilise the α-state or stabilise the β-state.

## 4. Discussion

### 4.1. Universal structural dynamics of AID/APOBEC3

AID/APOBEC enzymes are members of a structurally conserved family characterised by the CDA αβα-fold [10, 12, 17, 30, 32, 53, 56, 58, 94, 108, 109]. Despite this shared fold, they exhibit functional diversity due to differences in cellular localization and substrate specificity. These enzymes deaminate dC to dU, contributing to a range of biological functions. They restrict the activity of human endogenous retroviruses and zoonotic viruses, induce mutations in antibody hypervariable and switch regions, and, when acting off-target, can promote mutagenesis in host genomes, contributing to various cancers. All AID/APOBECs have diverged from their ancestral tRNA adenosine deaminases of the TadA/ADAT lineage, evolving to act on single-stranded nucleic acids like ssDNA [2, 3, 12, 13, 15, 54, 58, 67, 110]. This adaptation involved the formation of a DNA-binding groove formed by secondary catalytic ℓ1, ℓ3, ℓ5, and ℓ7. These loops enable recognition and processive deamination of ssDNA [6, 8, 10, 69, 111–116]. While APOBEC3s often target linear or hairpin-form ssDNA, AID shows a preference for binding DNA bubble substrates and G4 structures. Within the binding groove, the DNA adopts a U-shaped conformation, passing around ℓ1, constrained by ℓ3, and extending over ℓ5 and ℓ7 [12, 23, 30, 54, 65, 66, 117, 118].

Each loop has a distinct structural and functional role. Loop ℓ3, for example, contributes to dimerization interfaces and shields the groove. Loop ℓ5 provides pocket stability, while ℓ7, although not a direct part of the catalytic pocket, controls substrate access with ℓ1 and specificity. Substrate preferences across ℓ7 vary among enzymes: AID favours WR<u>C</u> motifs (ℓ7: YFCE) [71, 79], A3G-CD2 targets CC<u>C</u> (YDDQ) [11, 16, 79, 119, 120], and other APOBEC3s—A3A, A3B-CD2, A3C, A3F-CD2, A3G-CD1, and A3H hap II—prefer TT<u>C</u> (ℓ7: YDYD, YYFQ, YYFW, and YYHW, respectively) [11, 13, 16, 65–67, 72, 73, 121–123].

Analysis of AID/APOBEC3 protein dynamics revealed that the highest fluctuations occurred in ℓ1, ℓ2, ℓ3, and ℓ10 (Figure 2). In contrast, ℓ5 and ℓ7 exhibited lower fluctuations likely due to partial structural occlusion by adjacent loops. Loop ℓ5 maintains the integrity of the catalytic pocket, while ℓ7’s role in substrate specificity may favour structural rigidity. An exception was observed in AID’s ℓ7: though equal in length to A3A and A3B-CD2, it exhibited greater fluctuations in the β-state, which were suppressed in the α-state by the proximity of α7. This dynamic variability may reflect AID’s flexible recognition of WR<u>C</u> motifs compared to the stricter preferences of APOBEC3s [11, 13, 16, 65–67, 72, 73, 121–123]. Loops ℓ1 and ℓ3, being longer and more solvent exposed, showed greater flexibility. Notably, A3H hap II’s ℓ1, the longest of all, had the highest RMSF, especially in a region known to bind dsRNA and facilitate dimerization via nucleic acid bridging [30, 62, 94]. Loop ℓ3 showed highest fluctuations in A3A, A3B-CD2, and A3G-CD2, likely due to their extended 13-residue length. These longer loops not only remain solvent exposed but also engage in groove shielding, dimerization, and interdomain interactions in double-domain enzymes [76, 77]. Interestingly, A3G-CD2’s ℓ3 residues H258 and H250 may coordinate an additional zinc atom not modelled in our simulations. In contrast, shorter ℓ3s in A3G-CD1 and AID contributed to different groove dynamics; either blocking it or exposing it further to solvent, respectively. Loop ℓ2’s fluctuations, while notable, are likely due to its distal location and may indirectly influence DNA binding by affecting CD1-CD2 orientation in double-domain enzymes. Loop ℓ10 also exhibited flexibility in its N-terminal region. This mobility may help mediate interactions with domain linkers or contribute to the surface environment where domain-domain contact occurs. Finally, ℓ8 of A3G-CD1 may influence interactions with CD2’s β-bulge or ℓ10, depending on the protein’s orientation, potentially modulating stability or interdomain communication.

### 4.2. Dynamics of the catalytic pocket state

AID/APOBECs, as members of the CDA superfamily, share a conserved core, yet their catalytic behavior diverges due to differences in secondary catalytic loops. To examine how these loops influence function, we analysed the SASA of the catalytic glutamate. Median accessibility was generally consistent across the family, except for A3C (lower) and A3G-CD2 (higher) (Figure 3.A-C). Each protein exhibited different peak patterns in these regions, while A3Z1 subtypes also showed a higher number of nearby water molecules within 4 Å. This suggests that although proton shuttling may be broadly conserved, water-mediated reactivation of the catalytic glutamate could differ across proteins.

We next evaluated catalytic pocket volumes and observed distinct clustering patterns (Figure 3.D-E). A3Z2s shared similar peak volumes, as did A3Z1s. AID’s β-state aligned with peaks in A3A and A3G-CD2 (A3Z1), while A3H hap II’s peak was intermediate with a long tail extending into A3Z1-like volumes. These trends reflect evolutionary conservation within Z-domain subtypes and highlight AID’s unexpectedly large pocket volume. Having mapped the range of catalytic pocket volumes, we examined which conformations allowed dC binding. Volume alone does not indicate catalytic readiness, so we utilised molecular docking of dC based on our understanding of the Schrödinger CATalytic pocket, previously described for AID, A3A, and A3B [55, 64, 66].

Using molecular dynamics, we extended this approach to most AID/APOBEC3s. Across all AID/APOBEC3s, the volume ranges corresponding to closed, indeterminate, and open states were consistent, indicating evolutionary conservation of the pocket and secondary catalytic loop architecture (Figure 4.A). Nonetheless, key differences emerged. A3G-CD1 showed severely restricted accessibility, highlighting how structural changes can limit pocket opening. In contrast, A3A and A3G-CD2 reached the highest volumes, AID was intermediate, and A3B-CD2, A3C, A3F-CD2, A3G-CD1, and A3H hap II formed a lower-volume group. Open state prevalence also varied: A3G-CD2 was open most often, followed by AID (β-state), A3A, A3H hap II, and A3B-CD2—all frequently in an indeterminate state (Figure 4.B). A3C and A3F-CD2 remained mostly closed, while A3G-CD1 never opened. This contrasted with our prior assumptions for AID based on Schrödinger’s model but was confirmed via docking and random validation.

We also assessed pocket dynamics across simulations. A3C, A3F-CD2, and A3G-CD1 showed little variability, tending to remain in a single state (Figure 4.C-J). In contrast, A3A, A3B-CD2, A3H hap II, and AID frequently shifted between pocket states with A3G-CD2 showing moderate fluctuation. This suggests that while all AID/APOBEC3s access similar pocket volumes, evolutionary divergence influences which state is favoured and how frequently transitions occur. To investigate the structural drivers of these differences, we compared open vs. closed pocket conformations across AID/APOBEC3s (Figure 4). As shown in earlier studies [55], residues in ℓ3, ℓ5, and ℓ7 (e.g., W84 in AID) shifted the “walls” of the pocket to control access. However, we found that the catalytic residues themselves played the most central role. The α2 histidine showed the greatest mobility, potentially triggering state transitions. The nearby glutamate proton-shuttle, cysteine thiolates in α3, and the Zn²⁺ ion itself were more constrained but operated in concert. A threonine or serine in ℓ1, positioned on the catalytic pocket “floor,” often acted as a gatekeeper. While most APOBEC3s used a combination of conserved secondary loop residues to aid pocket state regulation, A3G-CD1 was an exception. Even when partially open, its pocket geometry was too narrow for dC to access the catalytic site with the proper angle for deamination. This suggests a unique structural limitation of A3G-CD1 which could also be shared by other CD1 counterparts. Nonetheless, across the family, our findings support the idea that pocket closure is primarily driven by conserved residues, reinforcing the concept that Schrödinger’s CATalytic pocket is a shared structural bottleneck in AID/APOBEC3 enzymes [64].

### 4.3. AID-α7’s α-state is more stable but can be destabilised by R171Y and R178D

Although the expected trend of catalytic pocket restriction through open, indeterminate, and closed states was observed for AID, our data revealed a higher proportion of open or indeterminate pockets than previously anticipated [55, 64]. To understand this discrepancy, we expanded our simulation set to include additional starting conformations of AID, allowing us to compare the protein beginning from either an open or closed catalytic pocket and from α7 in either its α-state or β-state (Figure 6.A-B). As full-length AID has never been crystallised, current AID structures lack the C-terminal α7 helix [10, 79]. Past AID structural predictions [10, 55, 79, 124] suggest two dominant α7 conformations: (1) the β-state, where α7 binds on the protein’s backside opposite the catalytic pocket, and (2) the α-state, where α7 interacts with the assistant patch and ℓ7 on the front side. Comparing β-state simulations that began with open vs. closed catalytic pockets showed minimal differences in α7 stability; however, simulations starting from a closed pocket more often sampled lower pocket volumes and were slightly more prone to remain in a closed or indeterminate state than those starting open (Figure 6.G, Figure 7.B,D-E). In contrast, comparing the α7 α-state to the β-state revealed striking differences. The α-state exhibited significantly less RMSF fluctuation (Figure 6.C-D) and remained stably bound between ℓ7 and α6. Meanwhile, in β-state simulations, α7 frequently dissociated from its initial conformation and transiently interacted with several AID surface regions. Notably, some β-state simulations transitioned into α-state-like conformations, underscoring the greater stability of the α7 α-state. Critically, we found that when α7 adopts the α-state, it engages with both ℓ1 and ℓ7 to constrain the catalytic pocket. This led to a smaller average pocket volume and a higher frequency of closed conformations (Figure 7.C-D,F-G). Because α7 positioning so directly influences pocket accessibility, we propose that α7 conformational switching acts as a structurally-inherent bottleneck that suppresses excessive mutagenesis in AID. This mechanism could contribute to AID’s comparatively slow catalytic rate, as the α-state not only obstructs G4 interaction at the assistant patch but also occludes the catalytic pocket itself. Moreover, these dynamic shifts may be modulated in vivo by AID’s interactions with G4 structures.

To further probe α7’s role in pocket regulation, we simulated three well-characterised AID C-terminal truncation mutants: 188-termination, 183-termination, and 181-termination (Figure 8.A) [80]. These variants exhibit different catalytic efficiencies. The 188- and 183-terminations remove α7 partially or entirely with ℓ11, while 181-termination also truncates ℓ11 and α6. Experimentally, 188- and 183-termination enhance catalytic activity, whereas 181-termination decreases it; however, no structural rationale was offered [80]. Structurally, RMSF analysis of these truncations revealed highly mobile C-termini, resembling the instability of AID’s β-state (Figure 8.B). We hypothesise that removing α7 disinhibits the catalytic pocket, increasing its openness in 188- and 183-terminations. In contrast, partial deletion of α6 in 181-termination may disrupt the assistant patch, destabilise the protein and thus impair catalysis. Although it is beyond the scope of this study, 181-termination might alter oligomerization or promote degradation.

Because the α7 α-state binds directly to the assistant patch (Figure 9.A-D), we next assessed whether mutations in this region disrupt α7 conformation. The R171Y mutant destabilised both α7 α- and β-states, causing early unraveling (<250 ns; Figure 9.E,I). R174E destabilised the α-state via increased contacts with the α7 backbone, R190, and R194, but didn’t affect the β-state (Figure 9.F,J). R177A did not affect the α-state but caused β-state unraveling similar to wild-type (Figure 9.G,K). For R178D, electrostatic repulsion with D187 and D188 destabilised both α7 conformations, redirecting binding to an alternative surface. Together, these data show that assistant patch residues are critical for α-state stabilisation: mutations like R171Y and R178D severely disrupt this conformation, while the β-state is more resistant; only R178D showed major effects. These results suggest that α7 α-state is incompatible with DNA-bound AID, where the assistant patch is engaged, potentially explaining AID’s low basal activity. It is tempting to propose that AID naturally prefers the α7 α-state as a default repressive conformation, thereby limiting access to both the catalytic site and assistant patch and thus limiting mutagenesis.

To examine whether α7 mutations themselves influence its conformation, we simulated several point mutants predicted to alter α-state stability (R190A, R190D, F193A, and R194A) (Figure 10.A-D). R190A and R190D preserved the α-state, though R190D formed weaker interactions due to charge repulsion with ℓ7 residues E117 and D118 (Figure 10.A-B,E-F). F193A partially unravelled α7, weakening interactions with Y114 and F115, yet retained overall α-state conformation (Figure 10.A-B,G). Interestingly, R194A actually reinforced α7 α-state stability (Figure 10.A-B,H). No single point mutation abolished α-state formation, reinforcing its robustness. However, the β-state of R190D was stabilised through salt bridges with R177 and R178, while R194A shifted to solvent-exposed regions on α6, potentially increasing assistant patch accessibility once AID transitions out of the α-state (Figure 10.C-D,J,L).

Finally, we assessed the impact of known mutations and PTMs outside of α7 thought to affect α7 conformations (Figure 11.A-D). H130, which is known from Hyper-IgM patients, stabilises α7’s β-state and the surrounding β5, α6, and ℓ11 [19, 79, 125]. This variant abolishes catalysis and disrupts both α7 states, though it had limited effect on α7 RMSF; although, it lead to partial unraveling of the α-state (Figure 11.A-B,E,I). L198, the C-terminal residue of AID, is structurally important; its mutation (L198S) completely unravelled α7 in the α-state. We also examined two PTMs: phosphorylation of T140 and Y184 [126, 127]. T140-p is located near the C-terminal carboxylate and was predicted to destabilise the β-state. Indeed, T140-p had no impact on the α-state but displaced β-state α7 to a new position on α6 (Figure 11.B,F,J). Y184-p of ℓ11, in contrast, stabilised the β-state without affecting the α-state (Figure 11.G,K). These findings suggest PTMs may serve as regulatory switches: T140-p could limit catalysis by destabilising the open-access β-state, while Y184-p may enhance catalysis by stabilising it. Together, our results demonstrate that α7 conformational dynamics, governed by assistant patch contacts, truncations, mutations, and PTMs, serve as a central structural switch for regulating AID catalytic pocket accessibility and activity.

### 4.4. Conclusion

This study presents an extensive series of molecular dynamics simulations of AID/APOBEC3 family members, designed to uncover both their general conformational behaviors and the specific biochemical mechanisms underpinning their activity. A key achievement was the characterization of catalytic pocket volumes and conformational states (open, closed, or indeterminate) across multiple family members. We identified structural features responsible for these states, highlighting the importance of loop and helix positioning. Going forward, these findings can be expanded through simulations of additional APOBECs, such as A3B-CD1, A3F-CD1, A3D, various A3H haplotypes, and APOBECs 1, 2, and 4. It will also be valuable to examine double-domain A3 proteins in different conformational states, globular vs. dumbbell, to determine how interdomain interactions or cofactors like Zn²⁺ (e.g., in ℓ3 of A3G-CD2) influence pocket dynamics. Comparative simulations across species, point mutations, or PTMs could further clarify the evolutionary and structural constraints of these proteins. One of the most unexpected findings was the remarkable stability of the α7 α-state relative to the β-state, which favoured closed catalytic pocket conformations. Mutational studies provided insights into key residues stabilising these α7 conformations. Future work should investigate combinations of mutations and PTMs within α7 across evolutionary variants. Finally, this study demonstrated how pocket states are influenced by ℓ1 threonine/serine movement and core catalytic residues, alongside accessory dynamics in surrounding DNA-binding loops. Importantly, we uncovered how α7 conformations influence pocket accessibility and how they themselves are modulated by mutations.

## Supplementary Materials

The following supporting information is Table S1: RCSB Deaminases Overview; Table S2: RCSB-PDB References of Deaminases across all types and species.

## Author Contributions

Conceptualization, D.N.G.H. and M.L.; software, D.N.G.H.; investigation, D.N.G.H.; writing—original draft preparation, D.N.G.H.; writing—review and editing, D.N.G.H, J.J.K, and M.L..; visualization, D.N.G.H.; supervision, M.L..; project administration, D.N.G.H.; funding acquisition, M.L. All authors have read and agreed to the published version of the manuscript.

## Institutional Review Board Statement

Not applicable.

## Informed Consent Statement

Not applicable.

## Acknowledgments

The authors are grateful to the aid of Gladdale Huebert as well as the other members of the Larijani Lab for editing this manuscript.

## Supplemental Data

**Figure S1.**
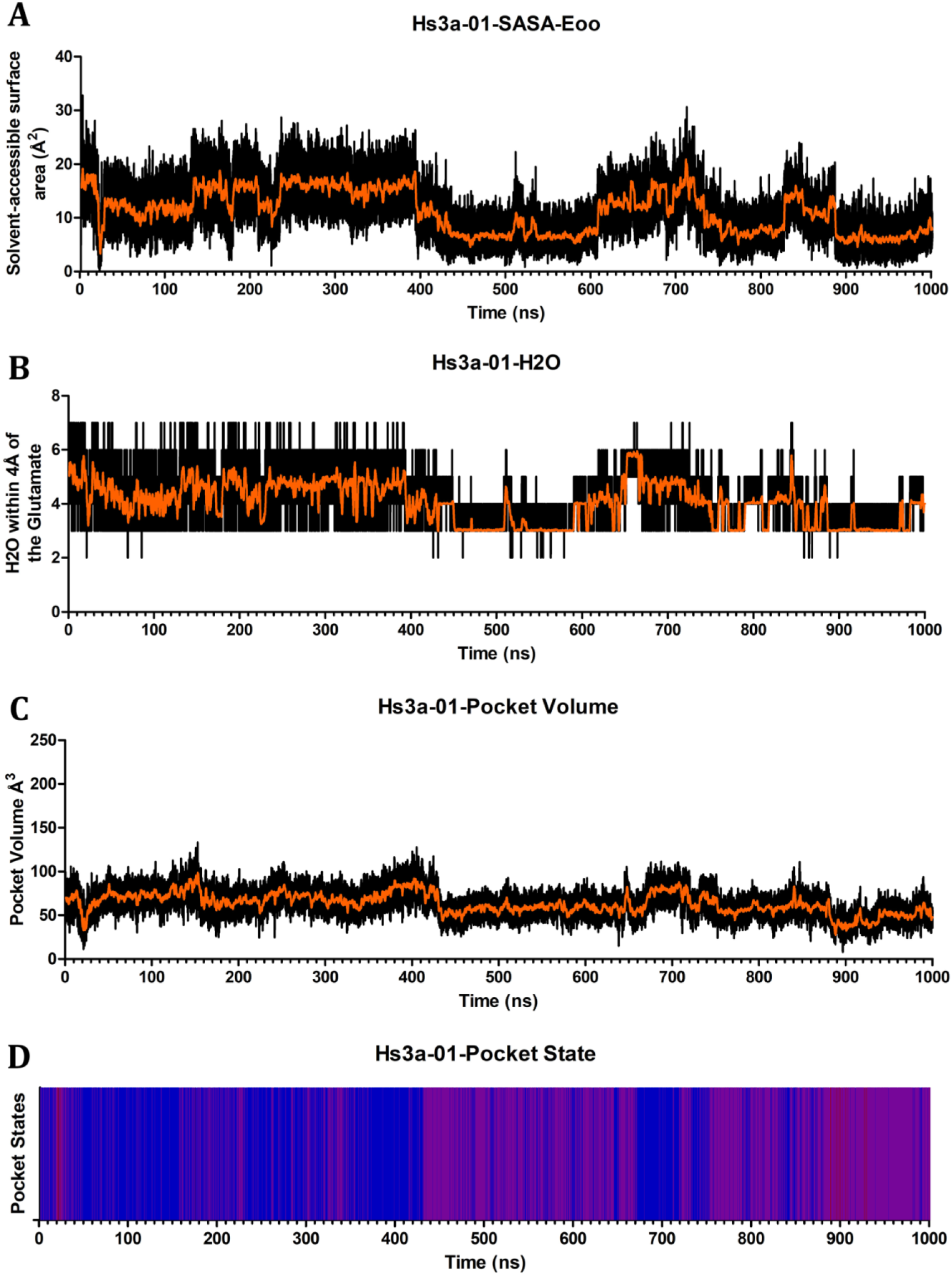
Pocket Dynamics of Hs3a-01. Graphs depict Hs3a-01 across 1µs simulation showing the (A) Solvent-Accessible Surface Area of the catalytic glutamates oxygens, (B) number of water molecules within 4Å of the catalytic glutamates oxygens, (C) pocket volume in Å^3^, and (D) pocket state coloured as open (blue), transient (purple), and closed (red).

**Figure S2.**
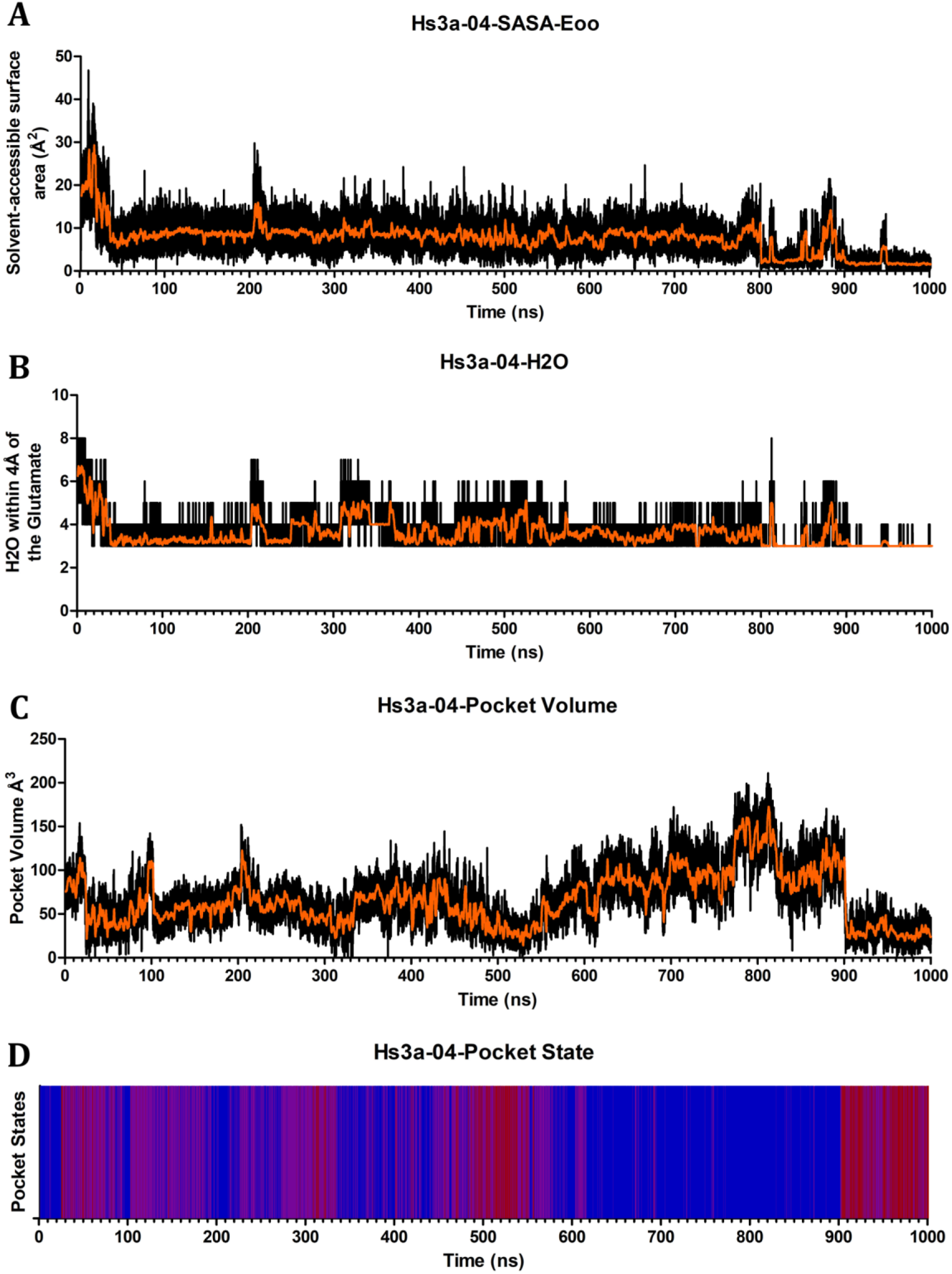
Pocket Dynamics of Hs3a-04. Graphs depict Hs3a-04 across 1µs simulation showing the (A) Solvent-Accessible Surface Area of the catalytic glutamates oxygens, (B) number of water molecules within 4Å of the catalytic glutamates oxygens, (C) pocket volume in Å^3^, and (D) pocket state coloured as open (blue), transient (purple), and closed (red).

**Figure S3.**
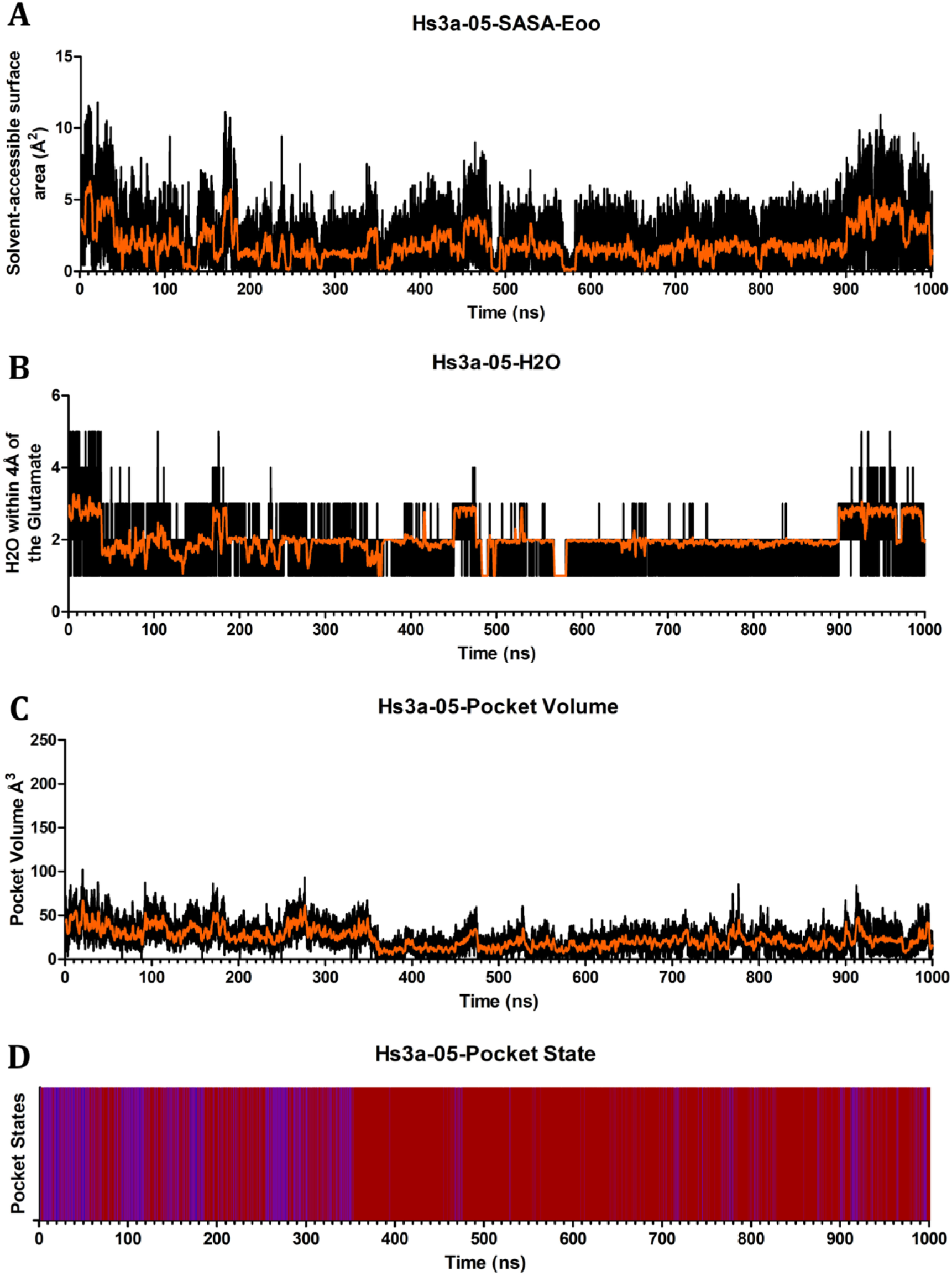
Pocket Dynamics of Hs3a-05. Graphs depict Hs3a-05 across 1µs simulation showing the (A) Solvent-Accessible Surface Area of the catalytic glutamates oxygens, (B) number of water molecules within 4Å of the catalytic glutamates oxygens, (C) pocket volume in Å^3^, and (D) pocket state coloured as open (blue), transient (purple), and closed (red).

**Figure S4.**
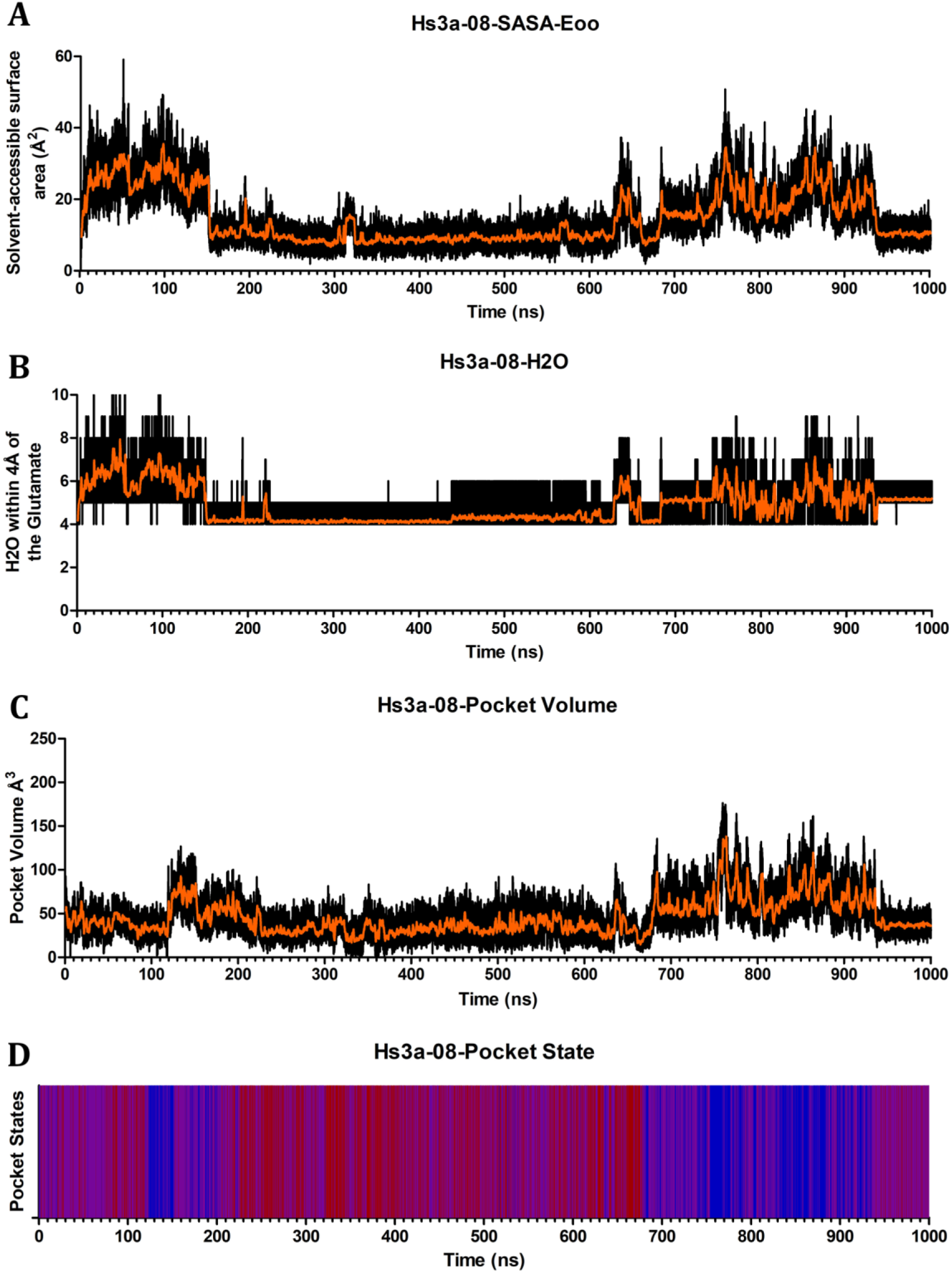
Pocket Dynamics of Hs3a-08. Graphs depict Hs3a-08 across 1µs simulation showing the (A) Solvent-Accessible Surface Area of the catalytic glutamates oxygens, (B) number of water molecules within 4Å of the catalytic glutamates oxygens, (C) pocket volume in Å^3^, and (D) pocket state coloured as open (blue), transient (purple), and closed (red).

**Figure S5.**
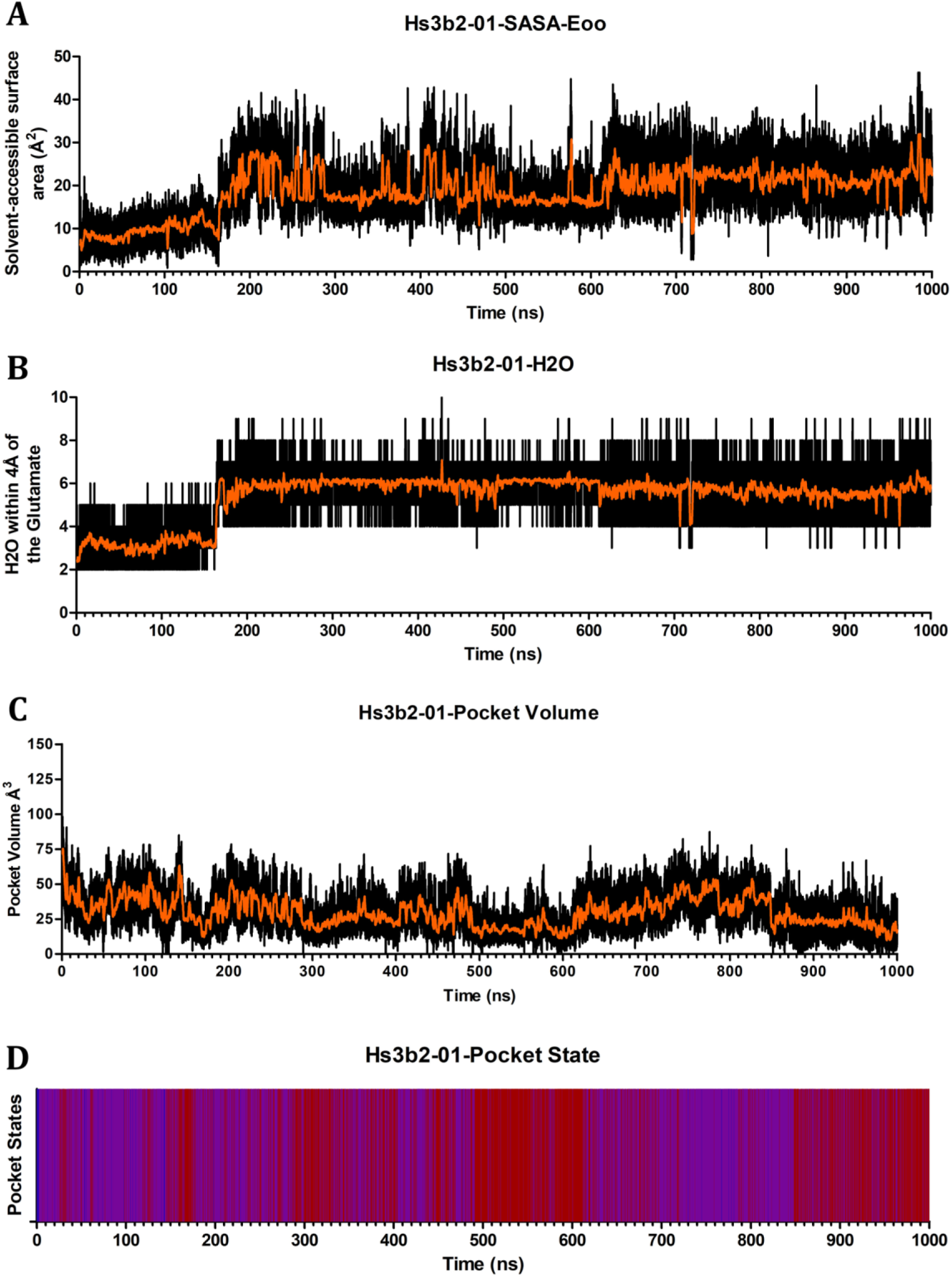
Pocket Dynamics of Hs3b2-01. Graphs depict Hs3b2-01 across 1µs simulation showing the (A) Solvent-Accessible Surface Area of the catalytic glutamates oxygens, (B) number of water molecules within 4Å of the catalytic glutamates oxygens, (C) pocket volume in Å^3^, and (D) pocket state coloured as open (blue), transient (purple), and closed (red).

**Figure S6.**
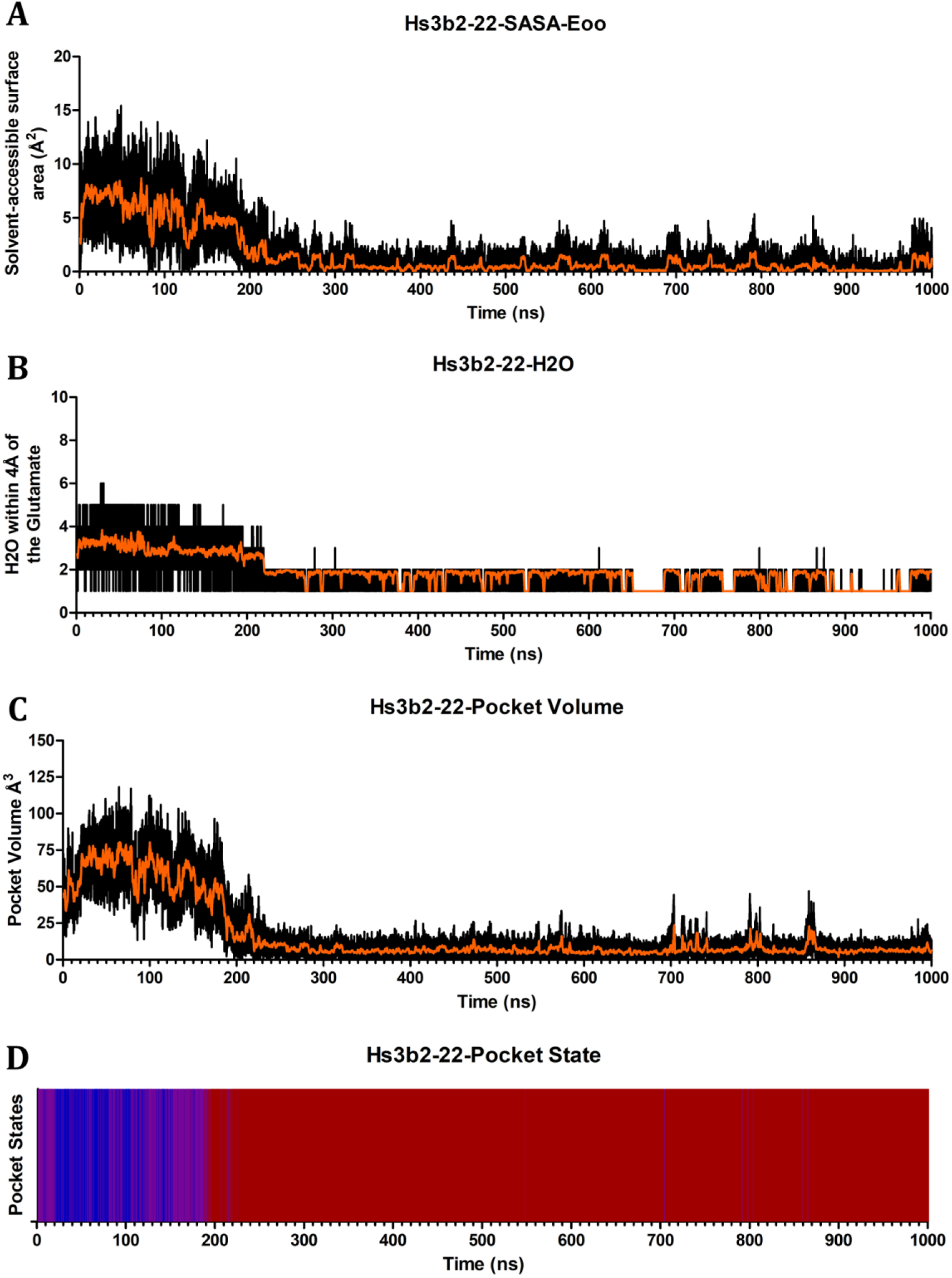
Pocket Dynamics of Hs3b2-22. Graphs depict Hs3b2-22 across 1μs simulation showing the (A) Solvent-Accessible Surface Area of the catalytic glutamates oxygens, (B) number of water molecules within 4Å of the catalytic glutamates oxygens, (C) pocket volume in Å^3^, and (D) pocket state coloured as open (blue), transient (purple), and closed (red).

**Figure S7.**
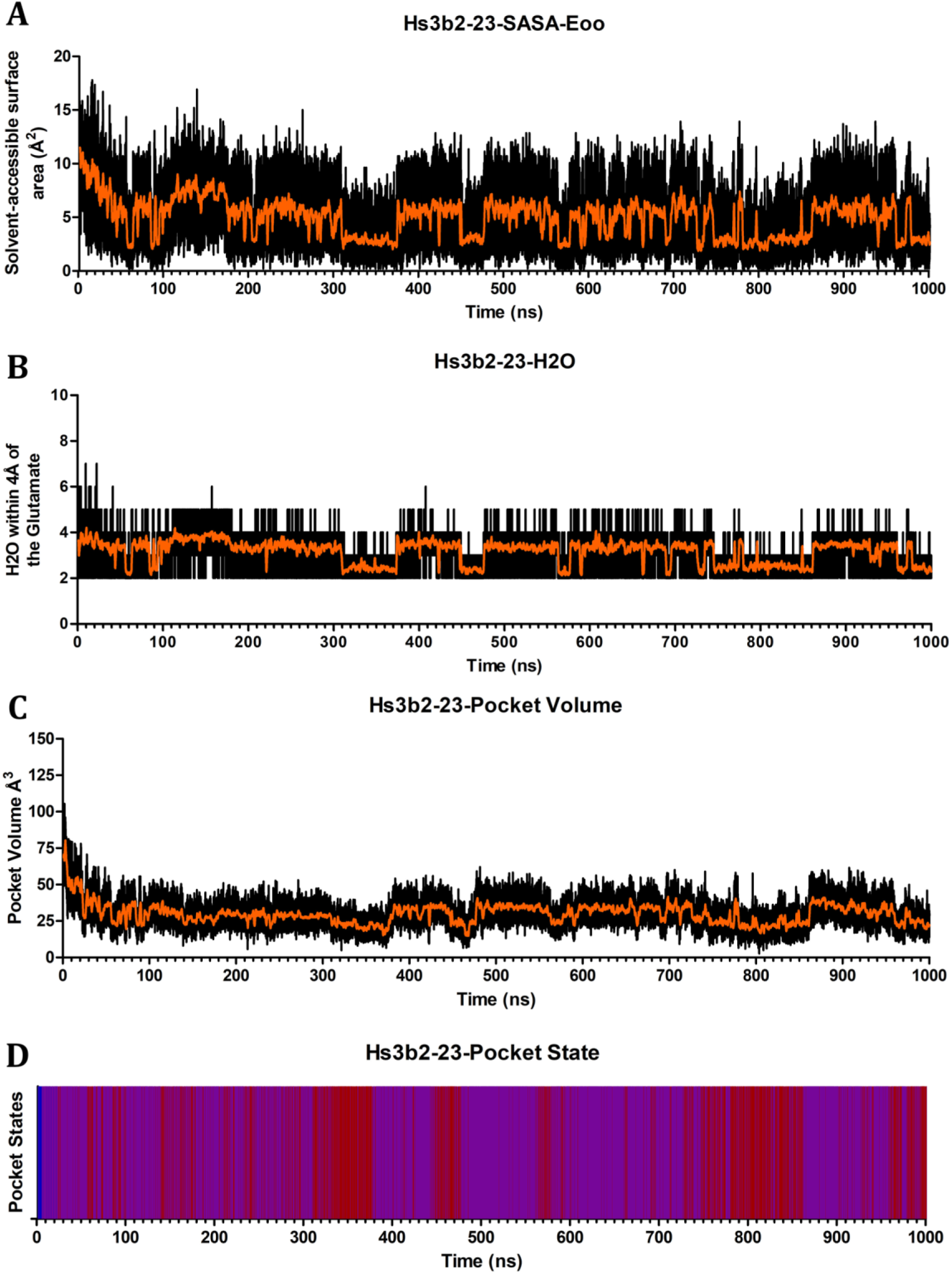
Pocket Dynamics of Hs3b2-23. Graphs depict Hs3b2-23 across 1μs simulation showing the (A) Solvent-Accessible Surface Area of the catalytic glutamates oxygens, (B) number of water molecules within 4Å of the catalytic glutamates oxygens, (C) pocket volume in Å^3^, and (D) pocket state coloured as open (blue), transient (purple), and closed (red).

**Figure S8.**
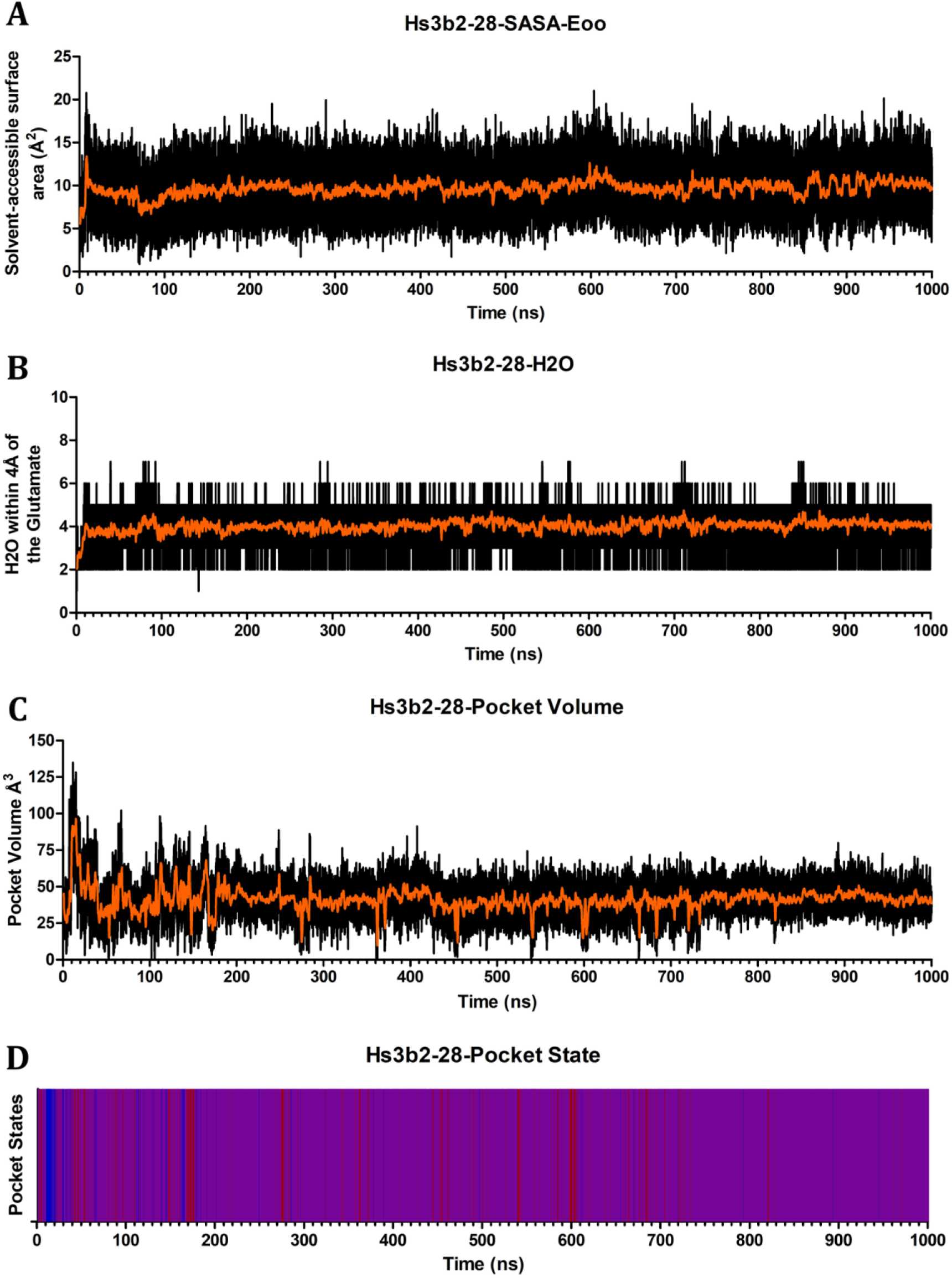
Pocket Dynamics of Hs3b2-28. Graphs depict Hs3b2-28 across 1μs simulation showing the (A) Solvent-Accessible Surface Area of the catalytic glutamates oxygens, (B) number of water molecules within 4Å of the catalytic glutamates oxygens, (C) pocket volume in Å^3^, and (D) pocket state coloured as open (blue), transient (purple), and closed (red).

**Figure S9.**
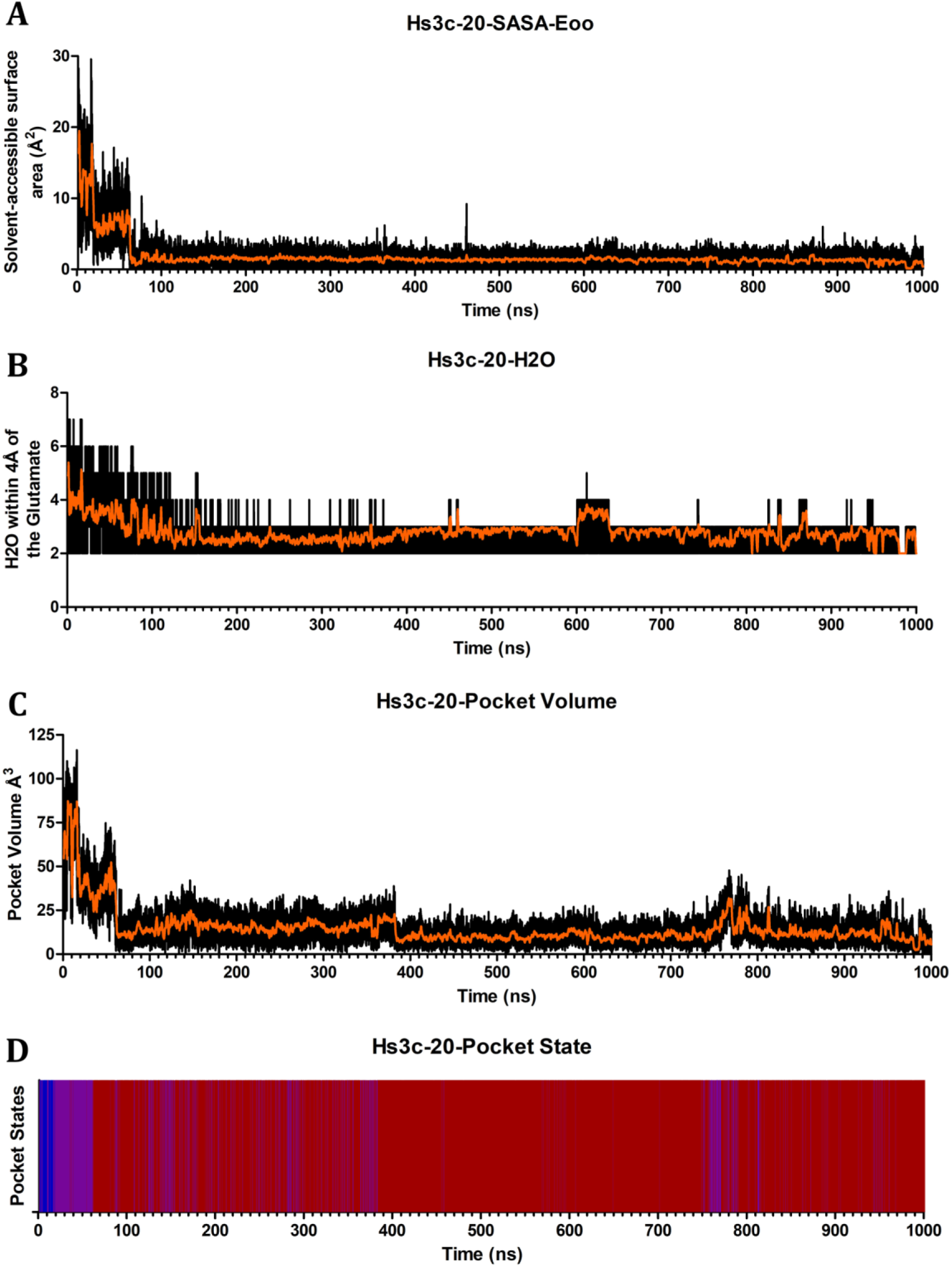
Pocket Dynamics of Hs3c-20. Graphs depict Hs3c-20 across 1μs simulation showing the (A) Solvent-Accessible Surface Area of the catalytic glutamates oxygens, (B) number of water molecules within 4Å of the catalytic glutamates oxygens, (C) pocket volume in Å^3^, and (D) pocket state coloured as open (blue), transient (purple), and closed (red).

**Figure S10.**
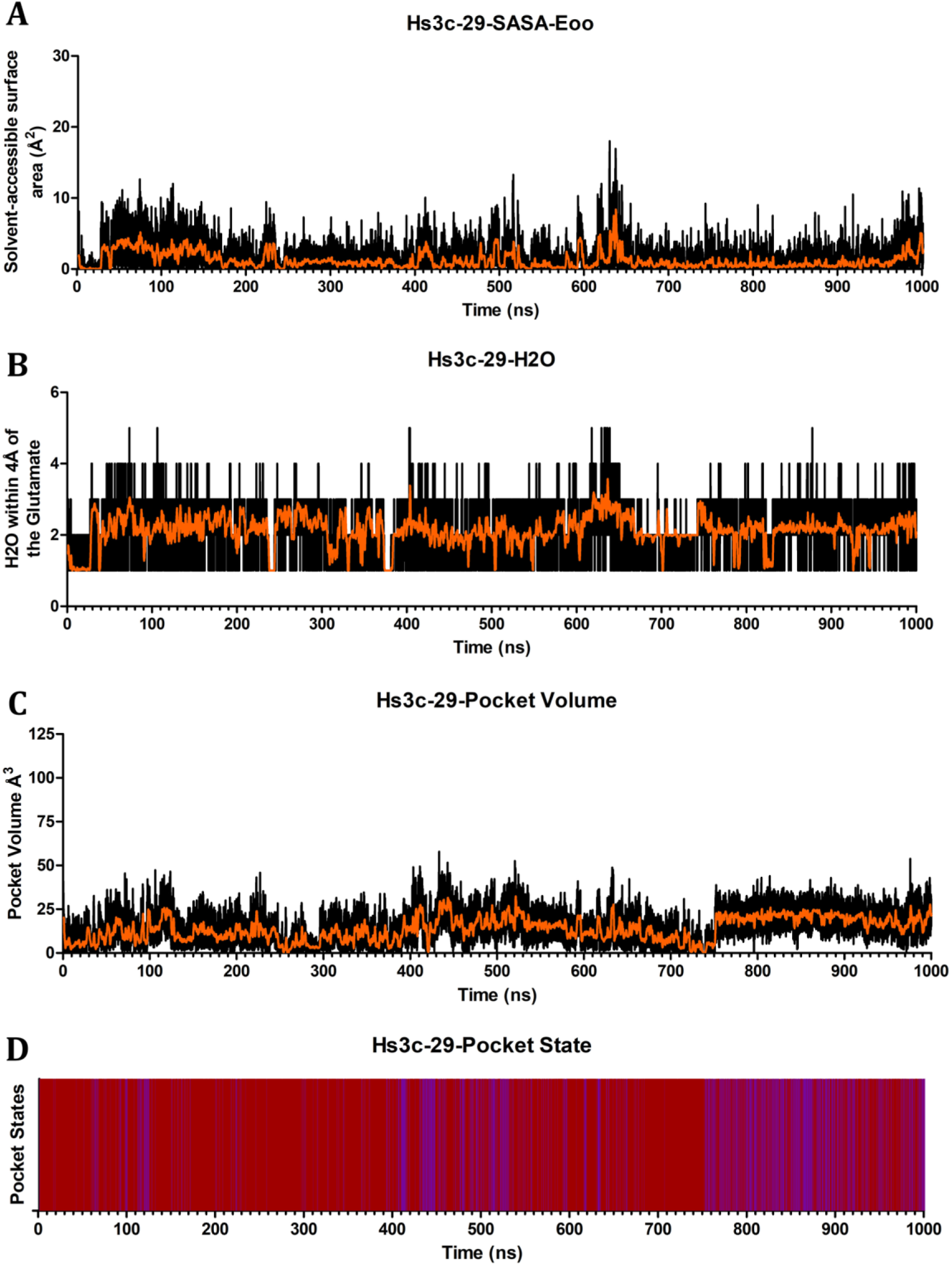
Pocket Dynamics of Hs3c-29. Graphs depict Hs3c-29 across 1μs simulation showing the (A) Solvent-Accessible Surface Area of the catalytic glutamates oxygens, (B) number of water molecules within 4Å of the catalytic glutamates oxygens, (C) pocket volume in Å^3^, and (D) pocket state coloured as open (blue), transient (purple), and closed (red).

**Figure S11.**
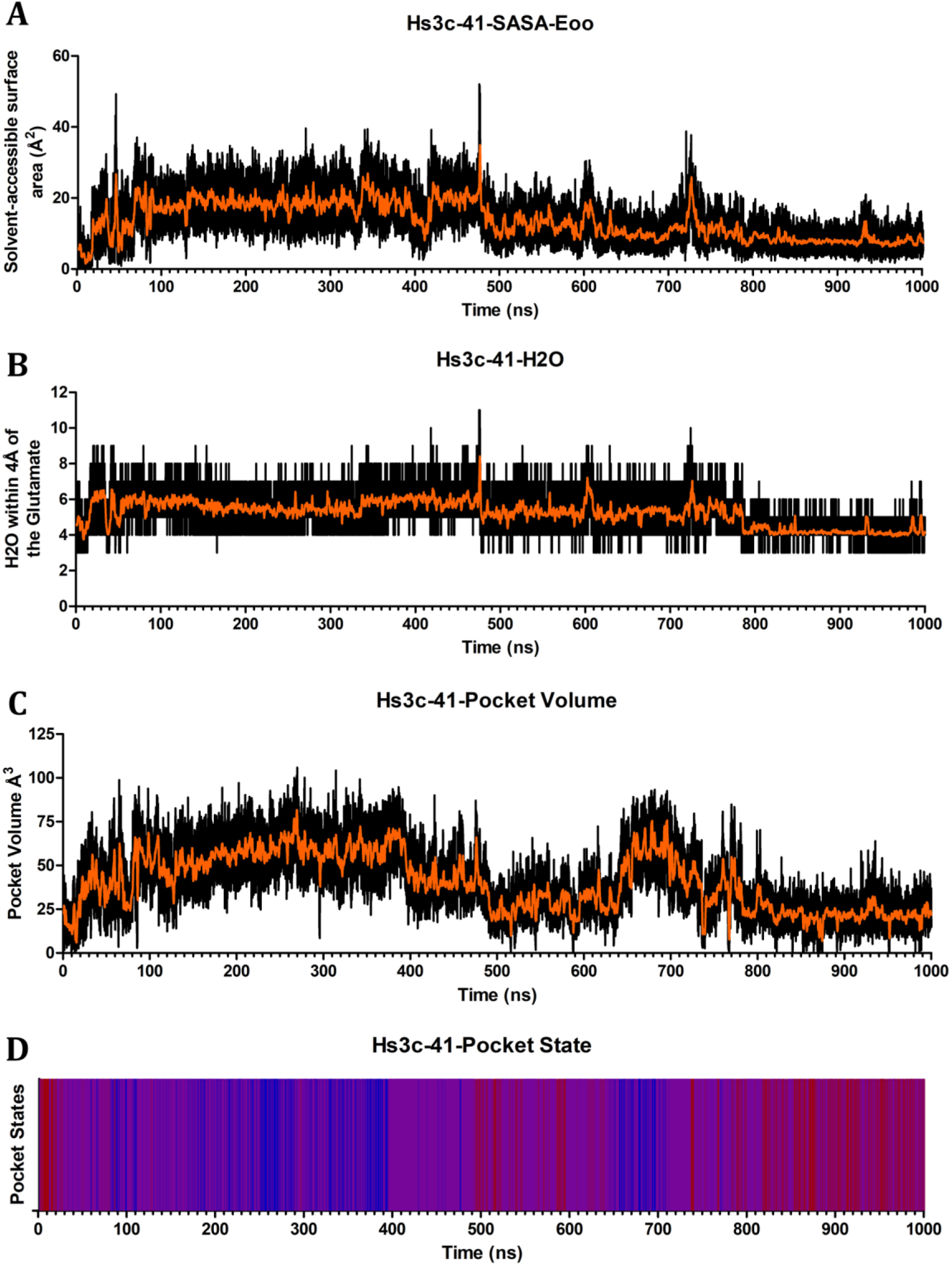
Pocket Dynamics of Hs3c-41. Graphs depict Hs3c-41 across 1μs simulation showing the (A) Solvent-Accessible Surface Area of the catalytic glutamates oxygens, (B) number of water molecules within 4Å of the catalytic glutamates oxygens, (C) pocket volume in Å^3^, and (D) pocket state coloured as open (blue), transient (purple), and closed (red).

**Figure S12.**
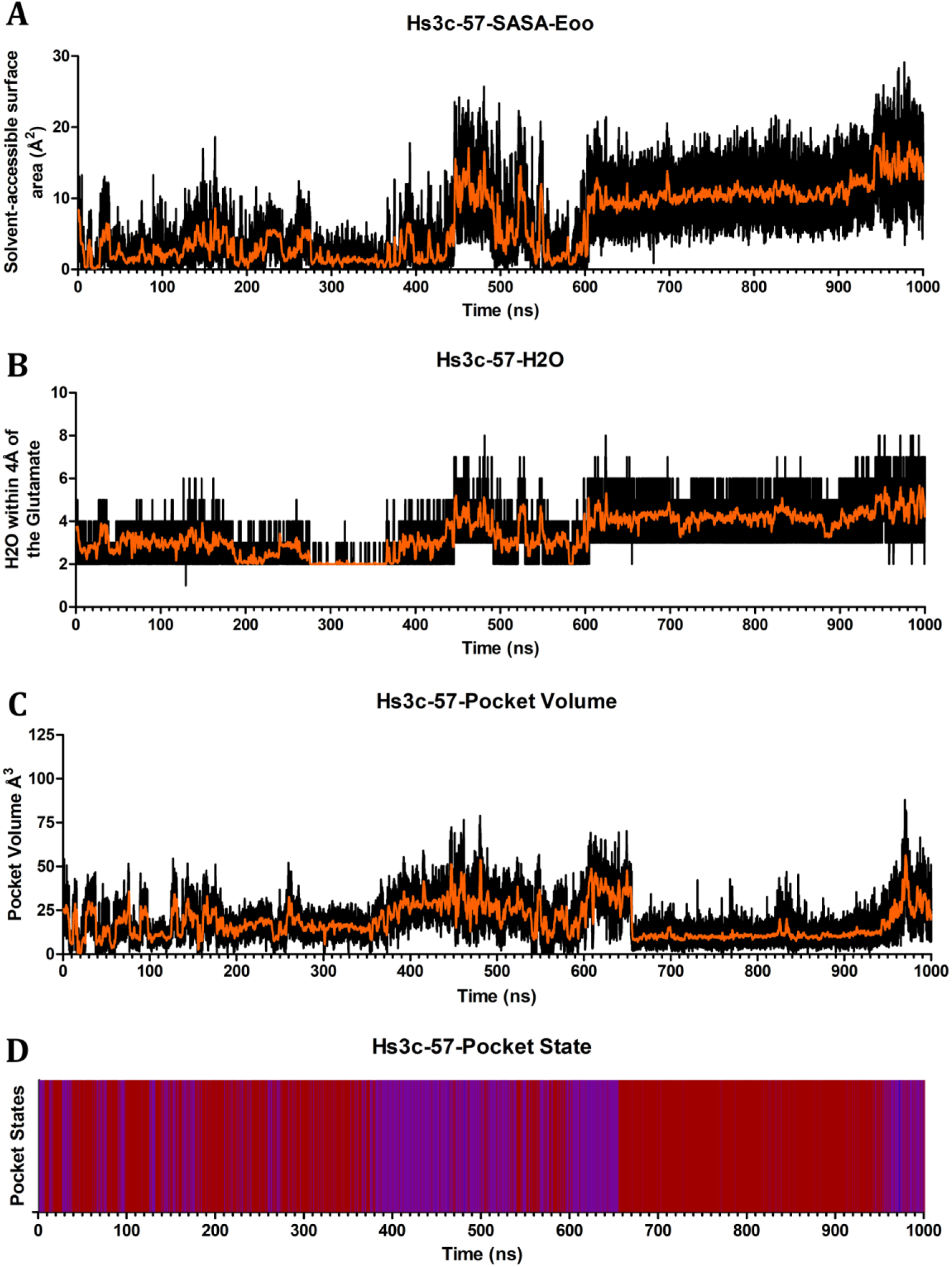
Pocket Dynamics of Hs3c-57. Graphs depict Hs3c-57 across 1μs simulation showing the (A) Solvent-Accessible Surface Area of the catalytic glutamates oxygens, (B) number of water molecules within 4Å of the catalytic glutamates oxygens, (C) pocket volume in Å^3^, and (D) pocket state coloured as open (blue), transient (purple), and closed (red).

**Figure S13.**
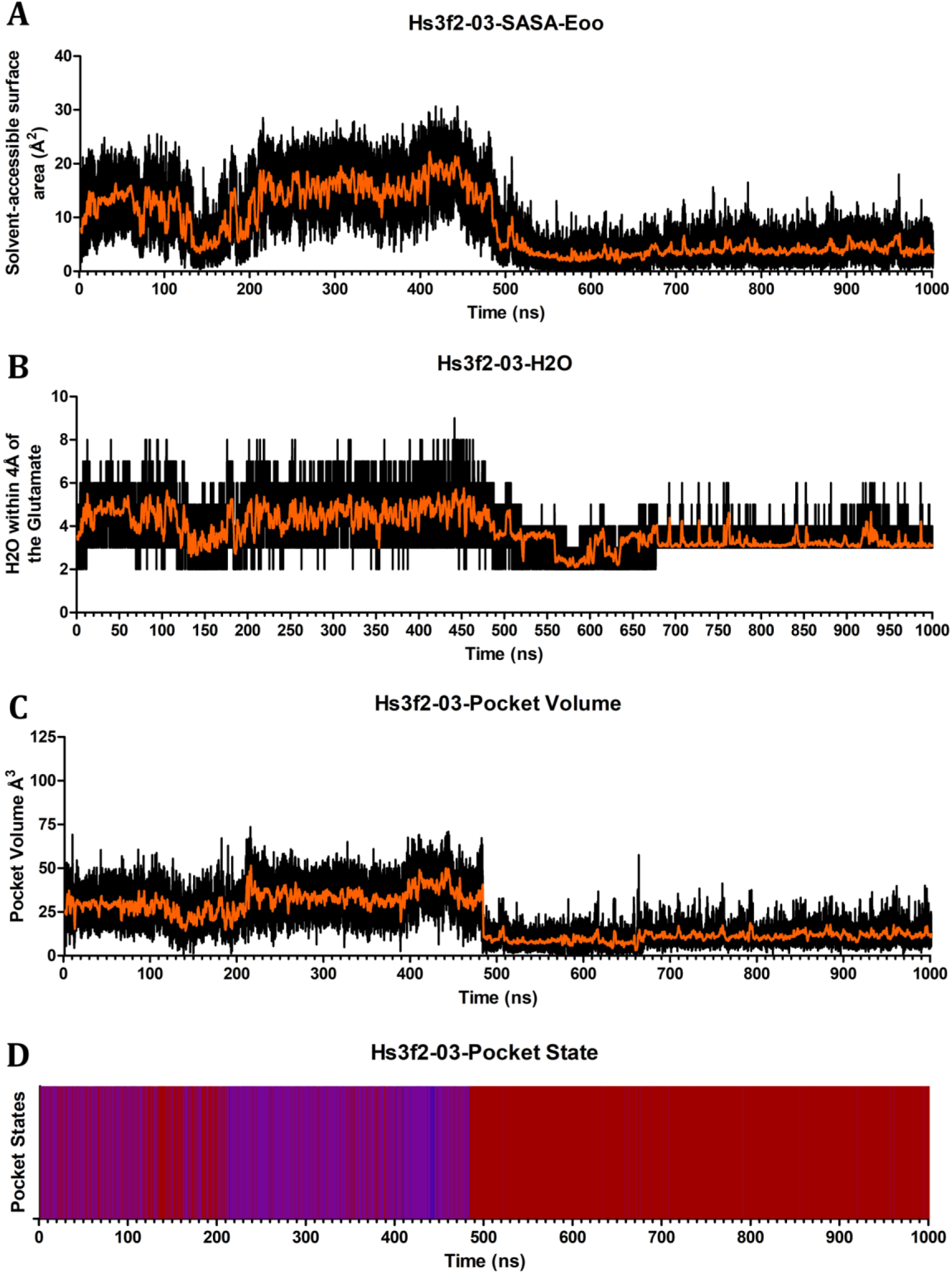
Pocket Dynamics of Hs3f2-03. Graphs depict Hs3f2-03 across 1μs simulation showing the (A) Solvent-Accessible Surface Area of the catalytic glutamates oxygens, (B) number of water molecules within 4Å of the catalytic glutamates oxygens, (C) pocket volume in Å^3^, and (D) pocket state coloured as open (blue), transient (purple), and closed (red).

**Figure S14.**
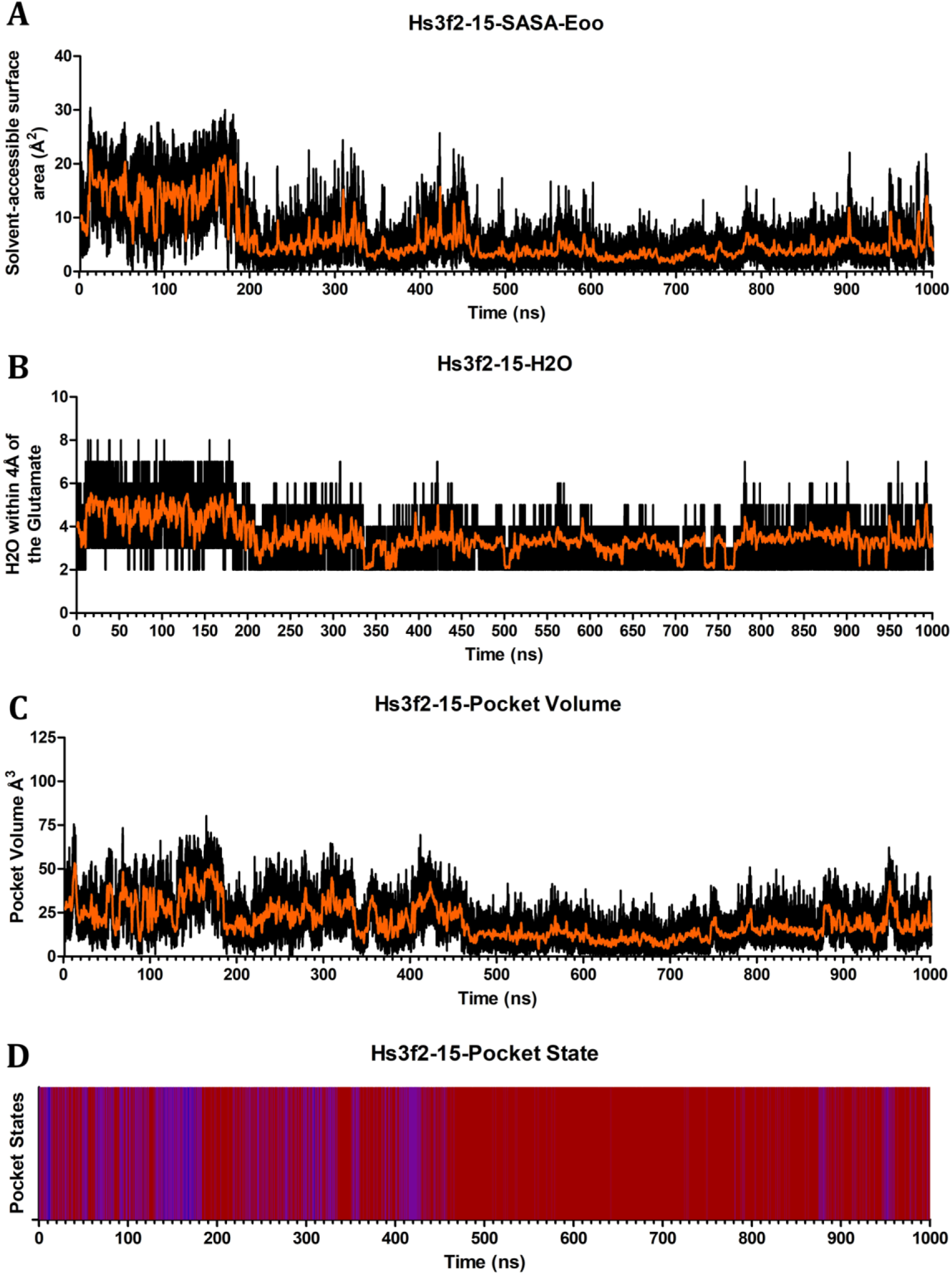
Pocket Dynamics of Hs3f2-15. Graphs depict Hs3f2-15 across 1μs simulation showing the (A) Solvent-Accessible Surface Area of the catalytic glutamates oxygens, (B) number of water molecules within 4Å of the catalytic glutamates oxygens, (C) pocket volume in Å^3^, and (D) pocket state coloured as open (blue), transient (purple), and closed (red).

**Figure S15.**
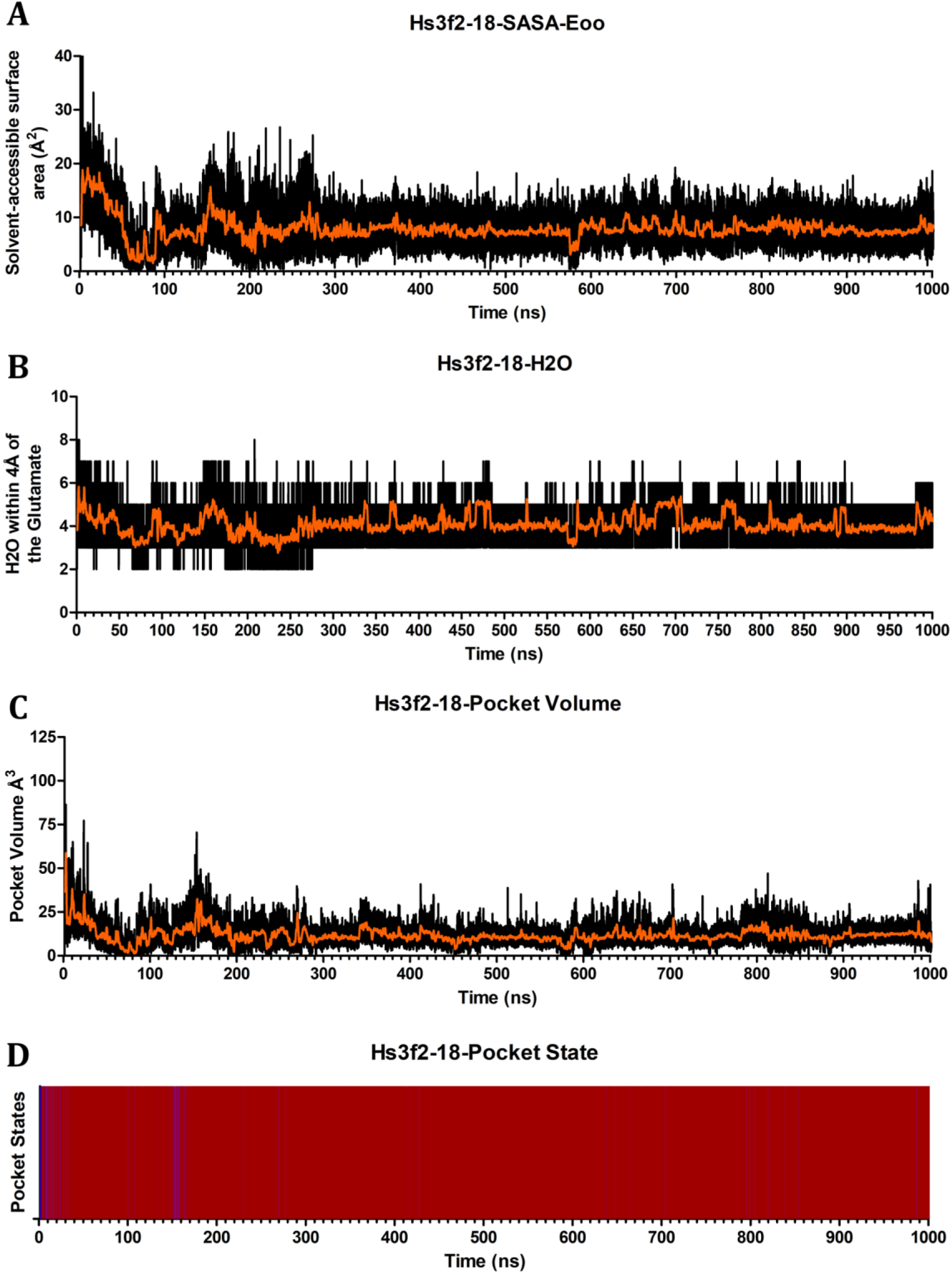
Pocket Dynamics of Hs3f2-18. Graphs depict Hs3f2-18 across 1μs simulation showing the (A) Solvent-Accessible Surface Area of the catalytic glutamates oxygens, (B) number of water molecules within 4Å of the catalytic glutamates oxygens, (C) pocket volume in Å^3^, and (D) pocket state coloured as open (blue), transient (purple), and closed (red).

**Figure S16.**
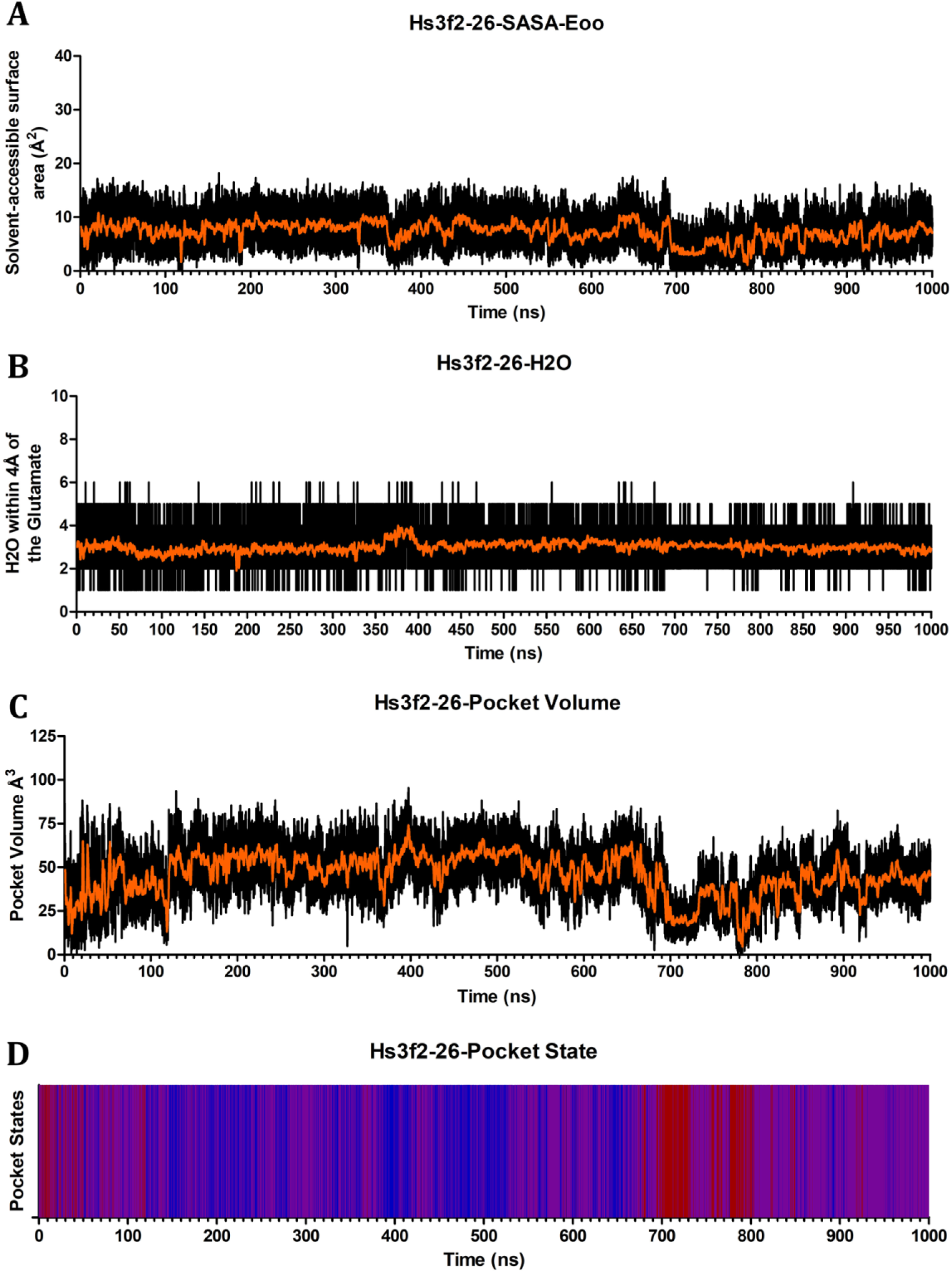
Pocket Dynamics of Hs3f2-26. Graphs depict Hs3f2-26 across 1μs simulation showing the (A) Solvent-Accessible Surface Area of the catalytic glutamates oxygens, (B) number of water molecules within 4Å of the catalytic glutamates oxygens, (C) pocket volume in Å^3^, and (D) pocket state coloured as open (blue), transient (purple), and closed (red).

**Figure S17.**
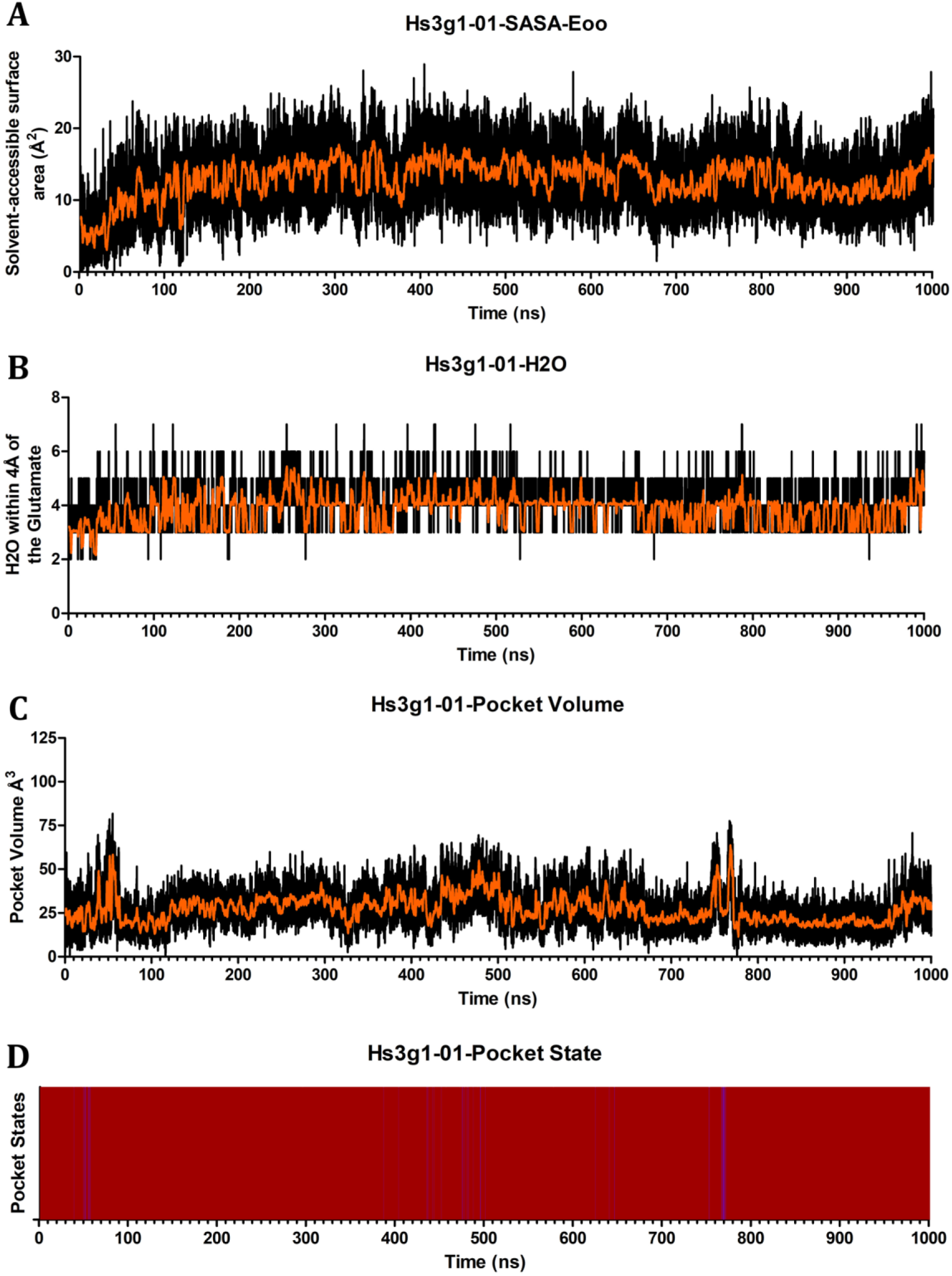
Pocket Dynamics of Hs3g1-01. Graphs depict Hs3g1-01 across 1μs simulation showing the (A) Solvent-Accessible Surface Area of the catalytic glutamates oxygens, (B) number of water molecules within 4Å of the catalytic glutamates oxygens, (C) pocket volume in Å^3^, and (D) pocket state coloured as open (blue), transient (purple), and closed (red).

**Figure S18.**
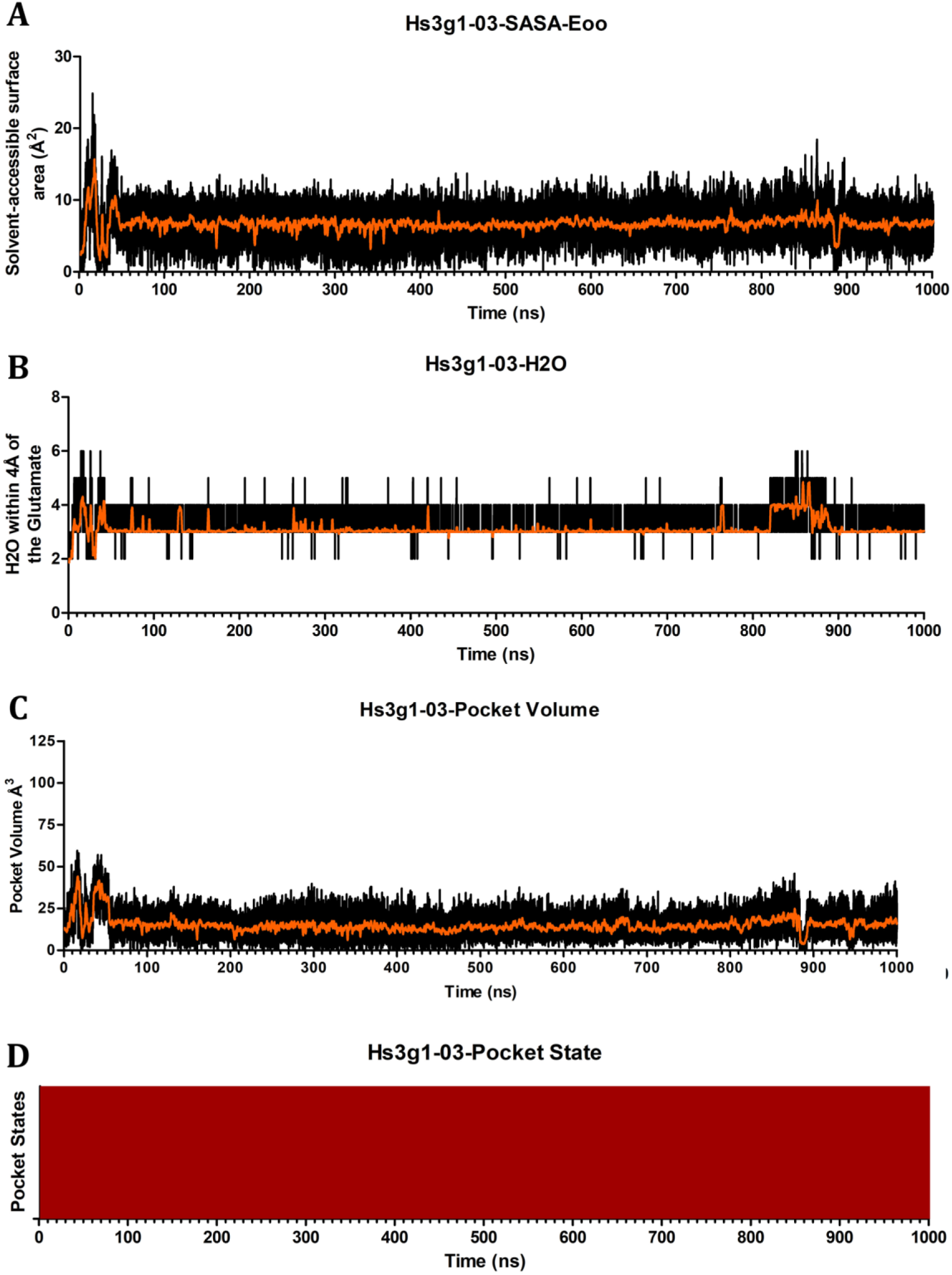
Pocket Dynamics of Hs3g1-03. Graphs depict Hs3g1-03 across 1μs simulation showing the (A) Solvent-Accessible Surface Area of the catalytic glutamates oxygens, (B) number of water molecules within 4Å of the catalytic glutamates oxygens, (C) pocket volume in Å^3^, and (D) pocket state coloured as open (blue), transient (purple), and closed (red).

**Figure S19.**
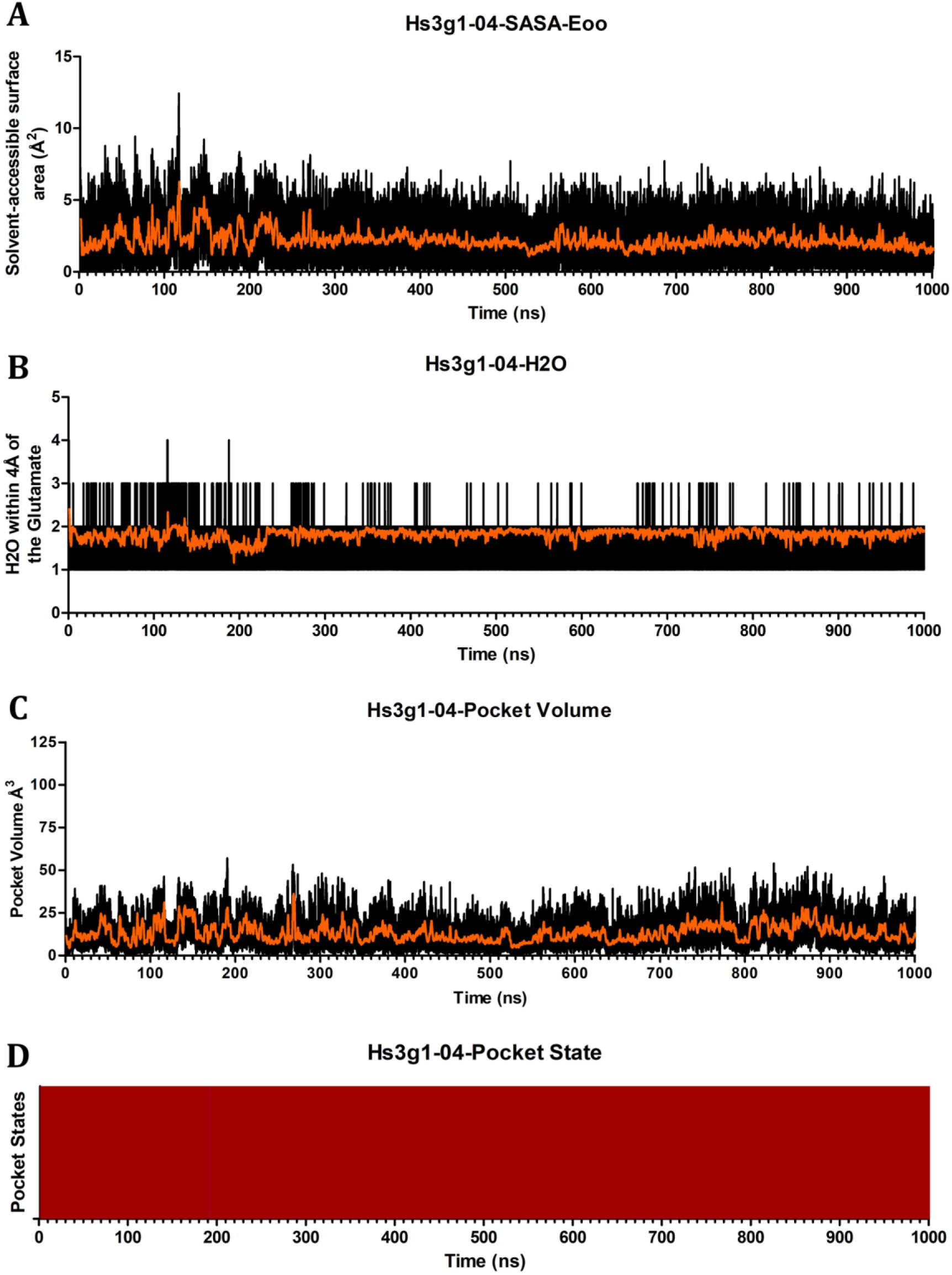
Pocket Dynamics of Hs3g1-04. Graphs depict Hs3g1-04 across 1μs simulation showing the (A) Solvent-Accessible Surface Area of the catalytic glutamates oxygens, (B) number of water molecules within 4Å of the catalytic glutamates oxygens, (C) pocket volume in Å^3^, and (D) pocket state coloured as open (blue), transient (purple), and closed (red).

**Figure S20.**
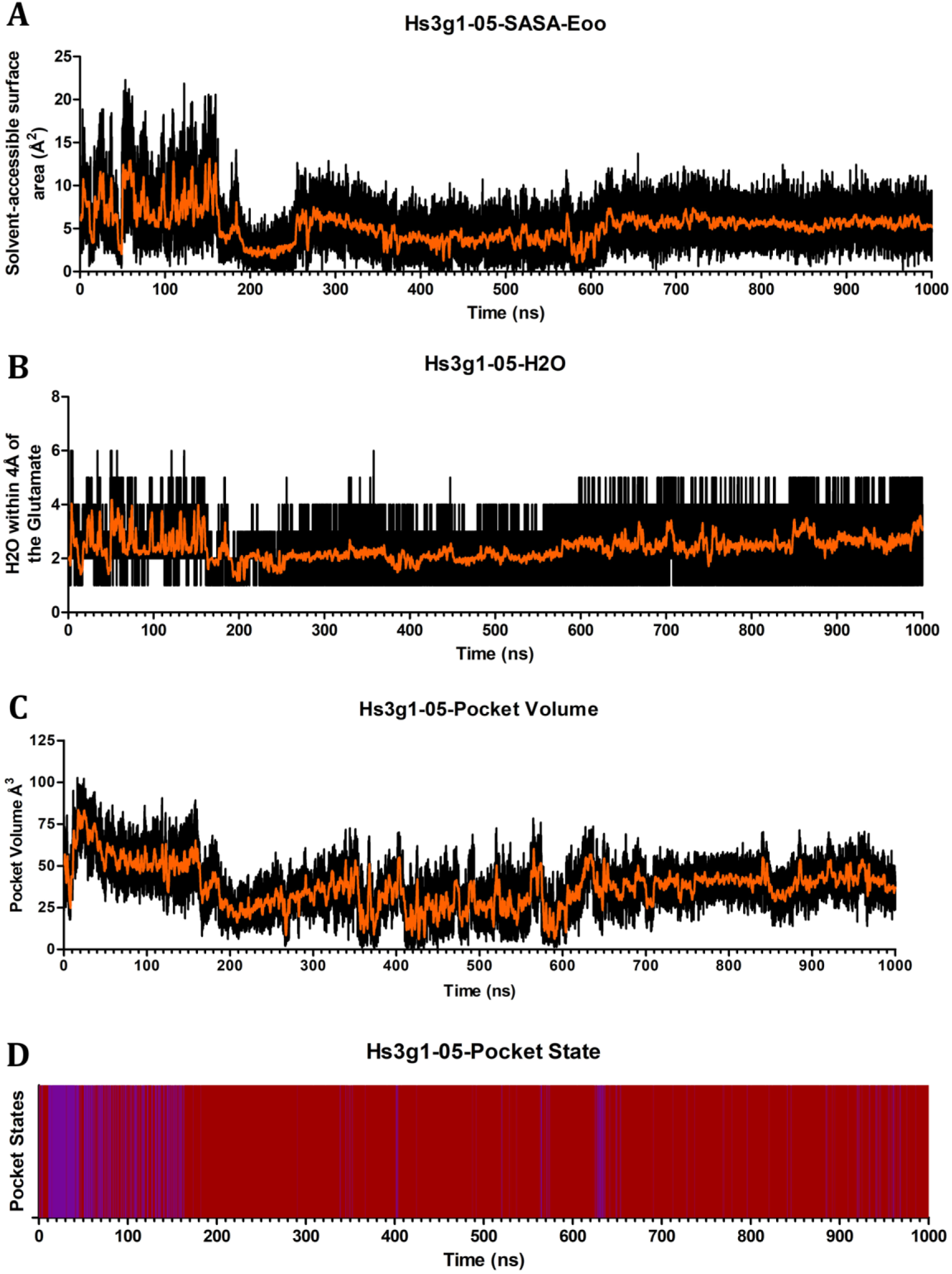
Pocket Dynamics of Hs3g1-05. Graphs depict Hs3g1-05 across 1μs simulation showing the (A) Solvent-Accessible Surface Area of the catalytic glutamates oxygens, (B) number of water molecules within 4Å of the catalytic glutamates oxygens, (C) pocket volume in Å^3^, and (D) pocket state coloured as open (blue), transient (purple), and closed (red).

**Figure S21.**
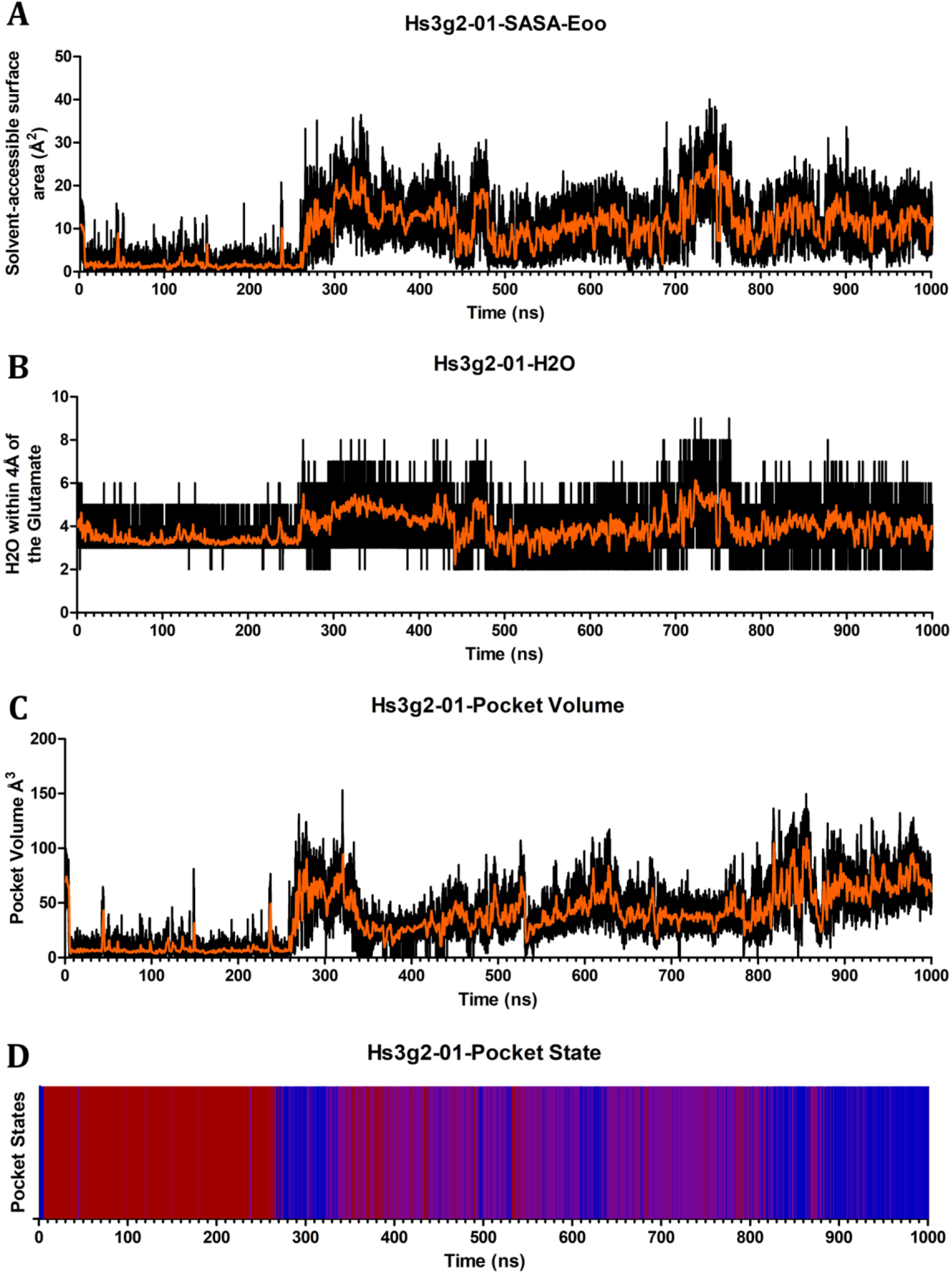
Pocket Dynamics of Hs3g2-01. Graphs depict Hs3g2-01 across 1μs simulation showing the (A) Solvent-Accessible Surface Area of the catalytic glutamates oxygens, (B) number of water molecules within 4Å of the catalytic glutamates oxygens, (C) pocket volume in Å^3^, and (D) pocket state coloured as open (blue), transient (purple), and closed (red).

**Figure S22.**
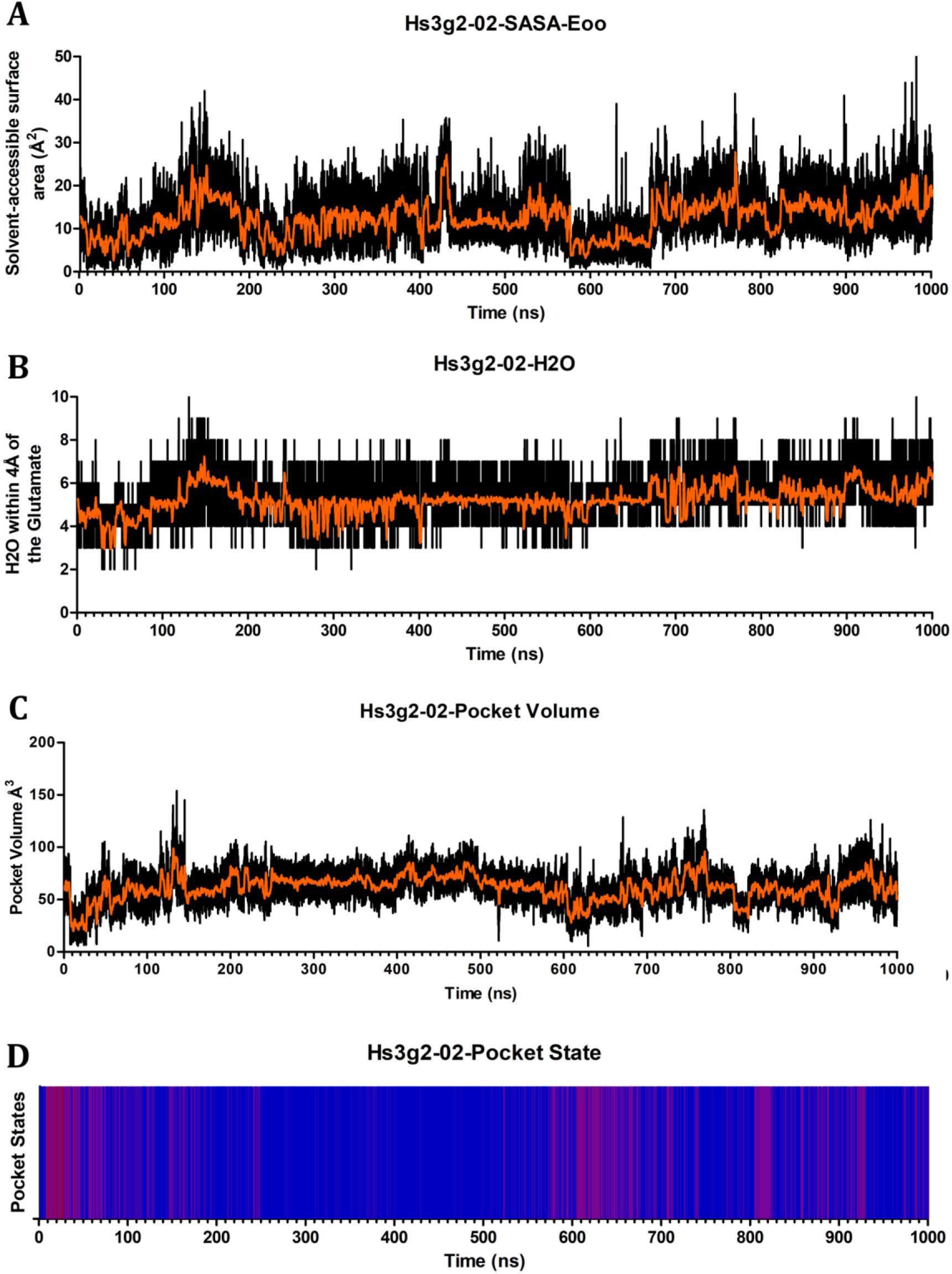
Pocket Dynamics of Hs3g2-02. Graphs depict Hs3g2-02 across 1μs simulation showing the (A) Solvent-Accessible Surface Area of the catalytic glutamates oxygens, (B) number of water molecules within 4Å of the catalytic glutamates oxygens, (C) pocket volume in Å^3^, and (D) pocket state coloured as open (blue), transient (purple), and closed (red).

**Figure S23.**
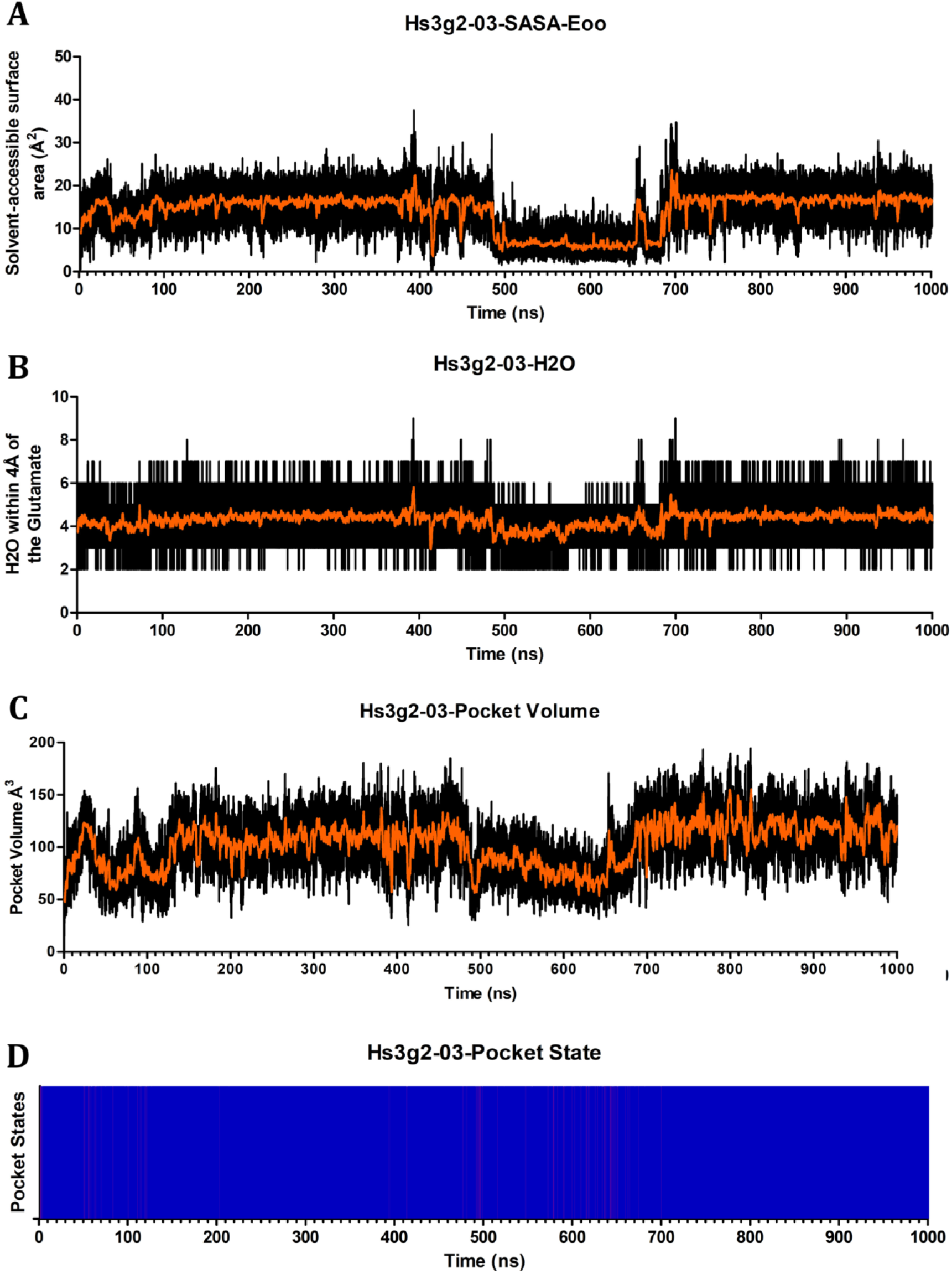
Pocket Dynamics of Hs3g2-03. Graphs depict Hs3g2-03 across 1μs simulation showing the (A) Solvent-Accessible Surface Area of the catalytic glutamates oxygens, (B) number of water molecules within 4Å of the catalytic glutamates oxygens, (C) pocket volume in Å^3^, and (D) pocket state coloured as open (blue), transient (purple), and closed (red).

**Figure S24.**
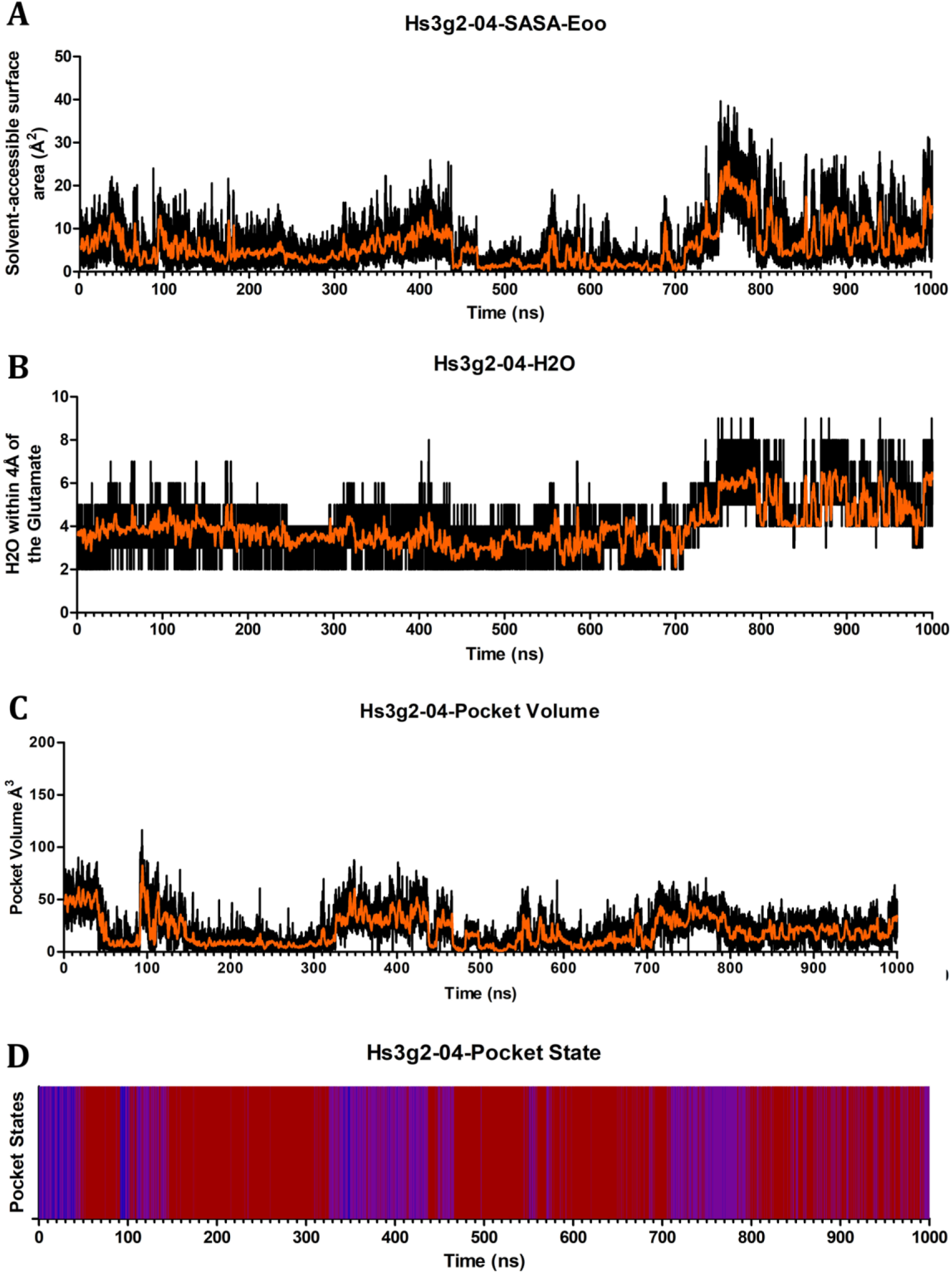
Pocket Dynamics of Hs3g2-04. Graphs depict Hs3g2-04 across 1μs simulation showing the (A) Solvent-Accessible Surface Area of the catalytic glutamates oxygens, (B) number of water molecules within 4Å of the catalytic glutamates oxygens, (C) pocket volume in Å^3^, and (D) pocket state coloured as open (blue), transient (purple), and closed (red).

**Figure S25.**
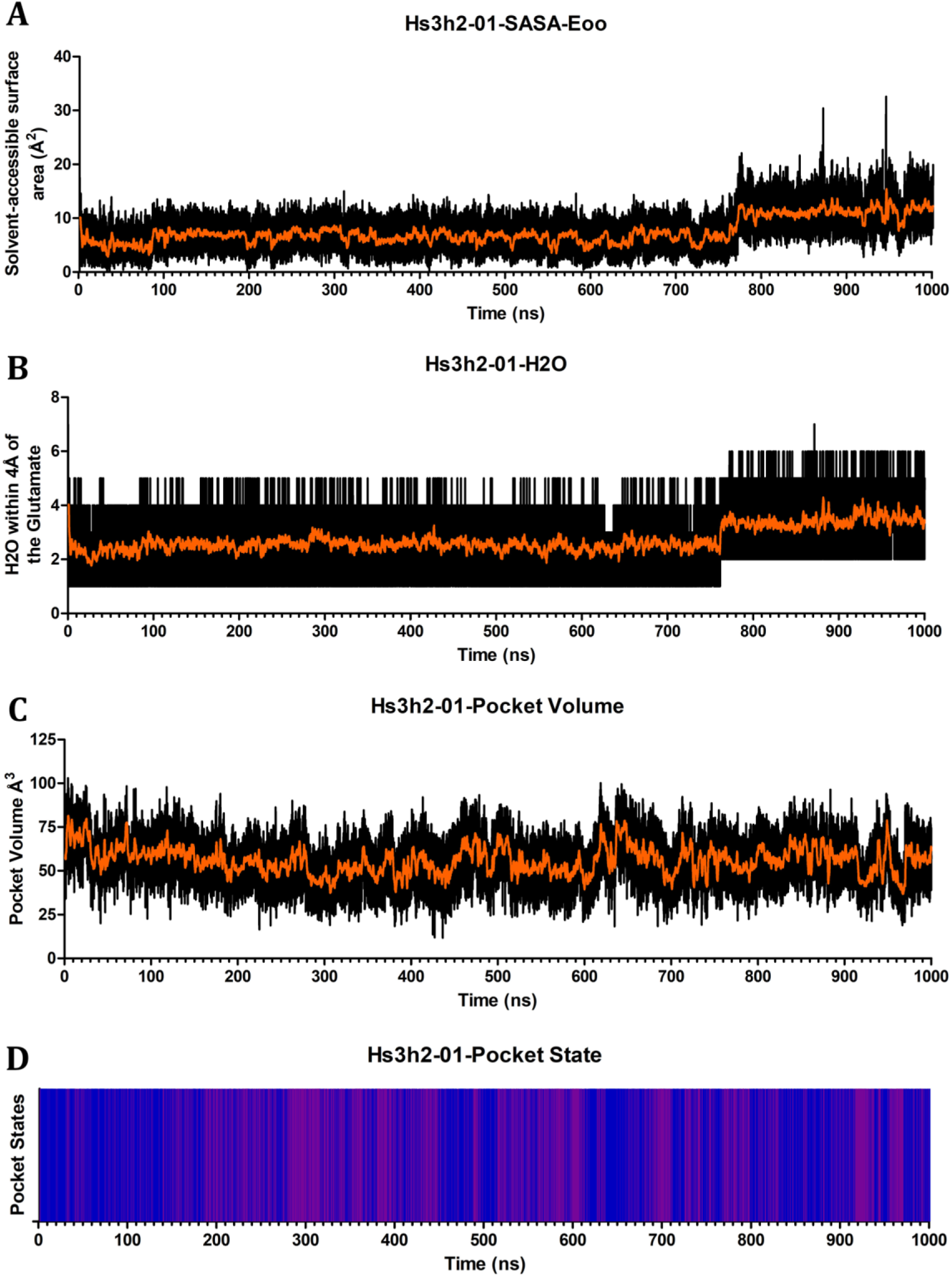
Pocket Dynamics of Hs3h2-01. Graphs depict Hs3h2-01 across 1μs simulation showing the (A) Solvent-Accessible Surface Area of the catalytic glutamates oxygens, (B) number of water molecules within 4Å of the catalytic glutamates oxygens, (C) pocket volume in Å^3^, and (D) pocket state coloured as open (blue), transient (purple), and closed (red).

**Figure S26.**
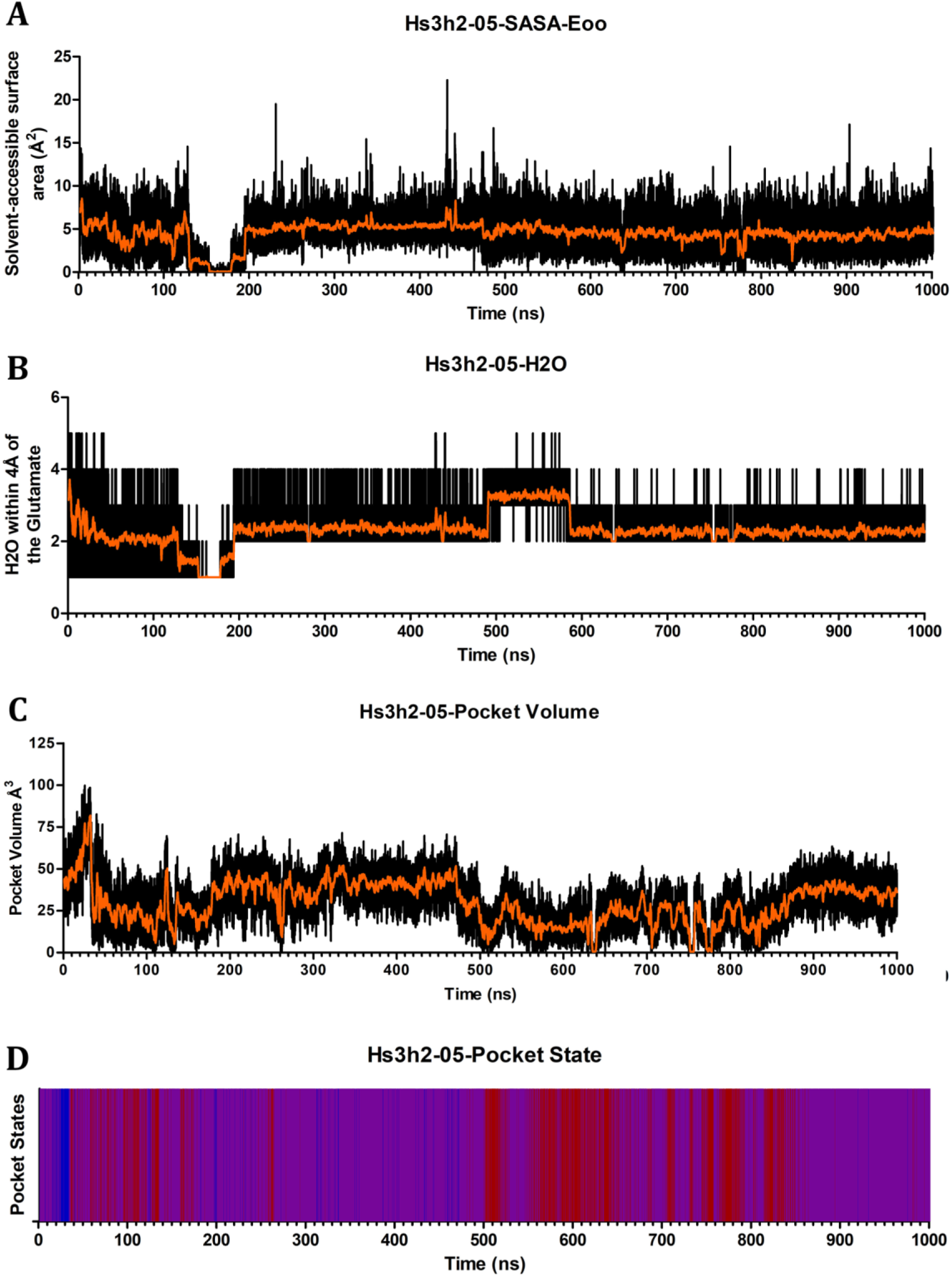
Pocket Dynamics of Hs3h2-05. Graphs depict Hs3h2-05 across 1μs simulation showing the (A) Solvent-Accessible Surface Area of the catalytic glutamates oxygens, (B) number of water molecules within 4Å of the catalytic glutamates oxygens, (C) pocket volume in Å^3^, and (D) pocket state coloured as open (blue), transient (purple), and closed (red).

**Figure S27.**
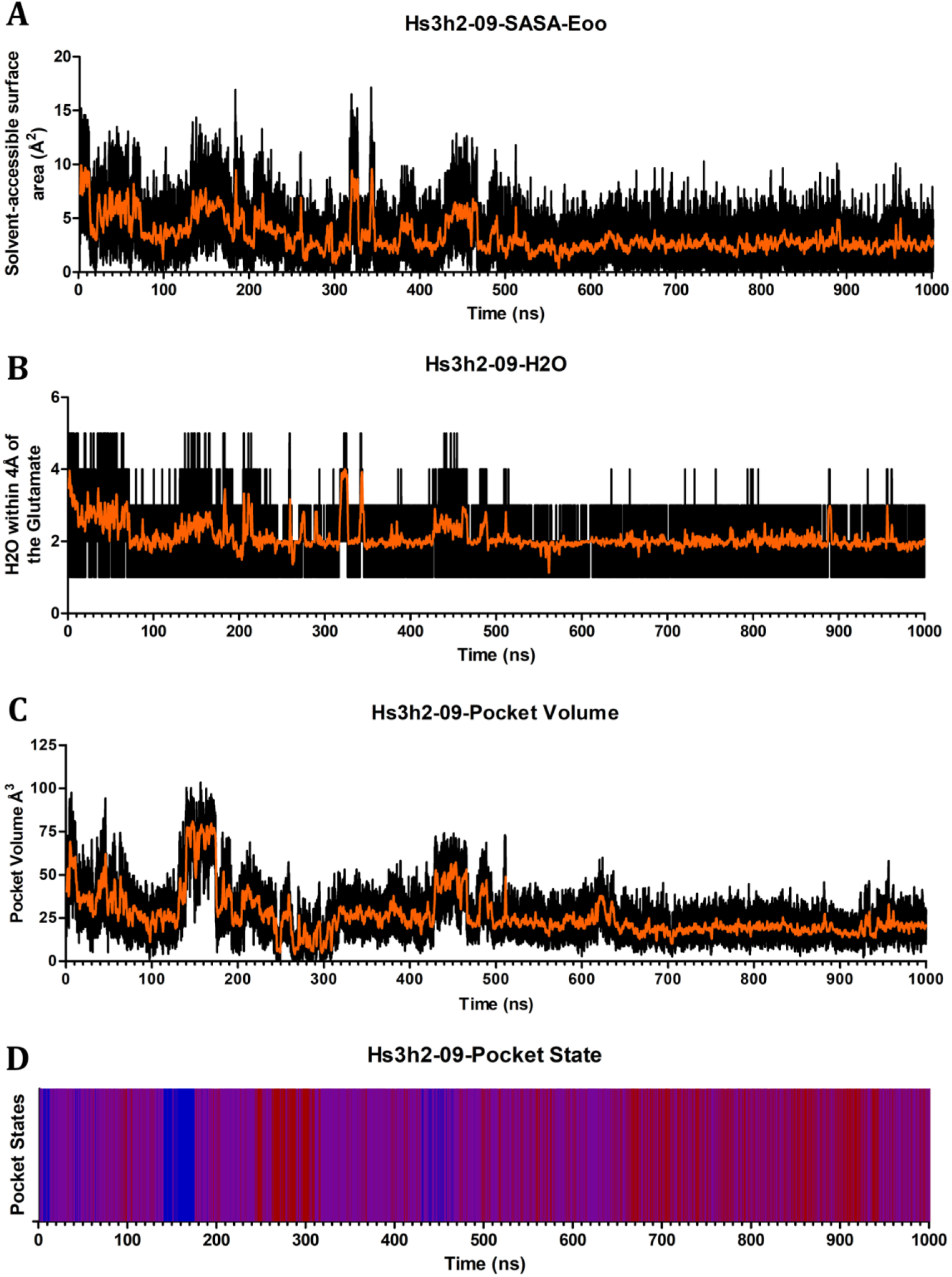
Pocket Dynamics of Hs3h2-09. Graphs depict Hs3h2-09 across 1μs simulation showing the (A) Solvent-Accessible Surface Area of the catalytic glutamates oxygens, (B) number of water molecules within 4Å of the catalytic glutamates oxygens, (C) pocket volume in Å^3^, and (D) pocket state coloured as open (blue), transient (purple), and closed (red).

**Figure S28.**
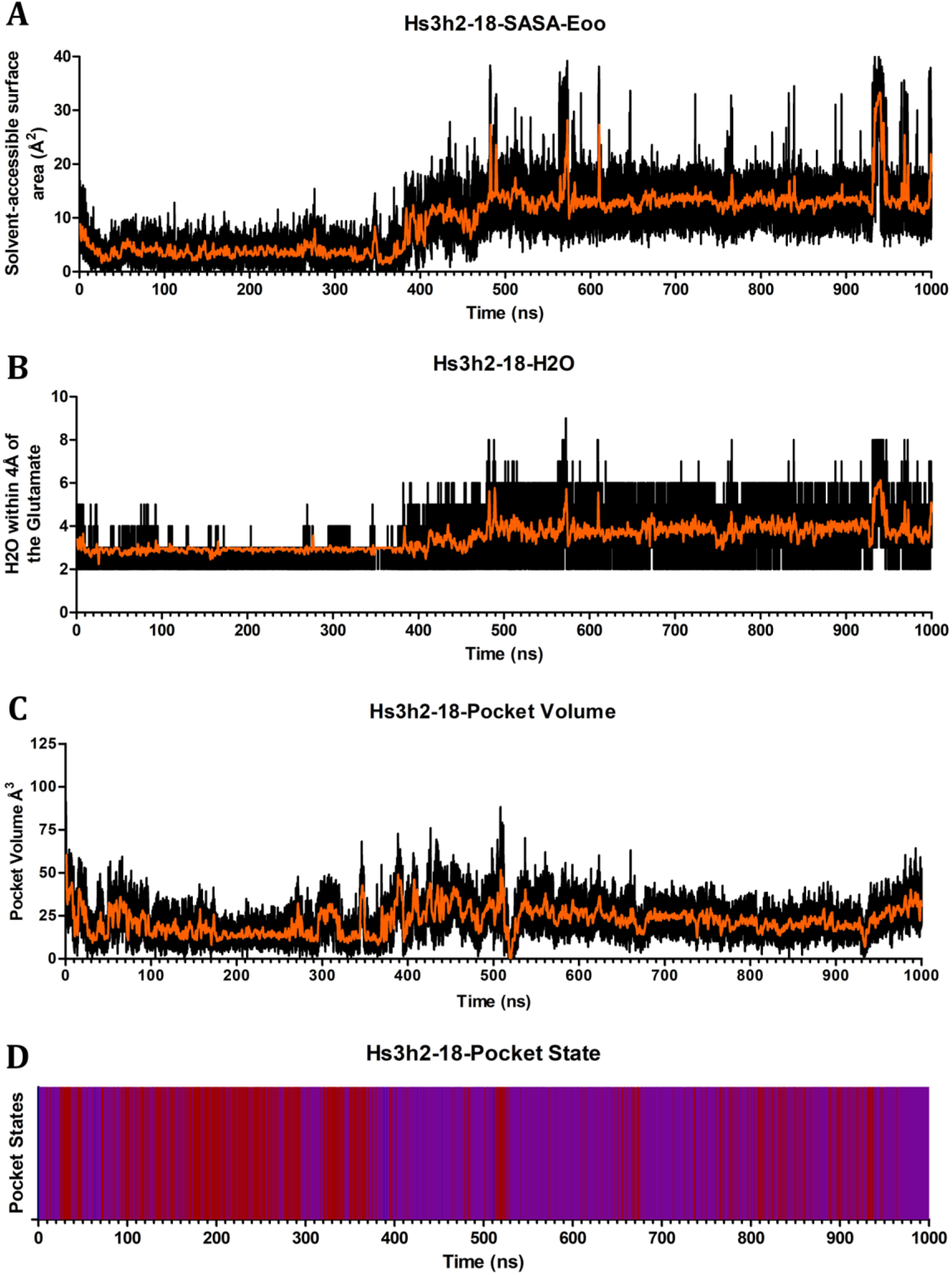
Pocket Dynamics of Hs3h2-18. Graphs depict Hs3h2-18 across 1μs simulation showing the (A) Solvent-Accessible Surface Area of the catalytic glutamates oxygens, (B) number of water molecules within 4Å of the catalytic glutamates oxygens, (C) pocket volume in Å^3^, and (D) pocket state coloured as open (blue), transient (purple), and closed (red).

**Figure S29.**
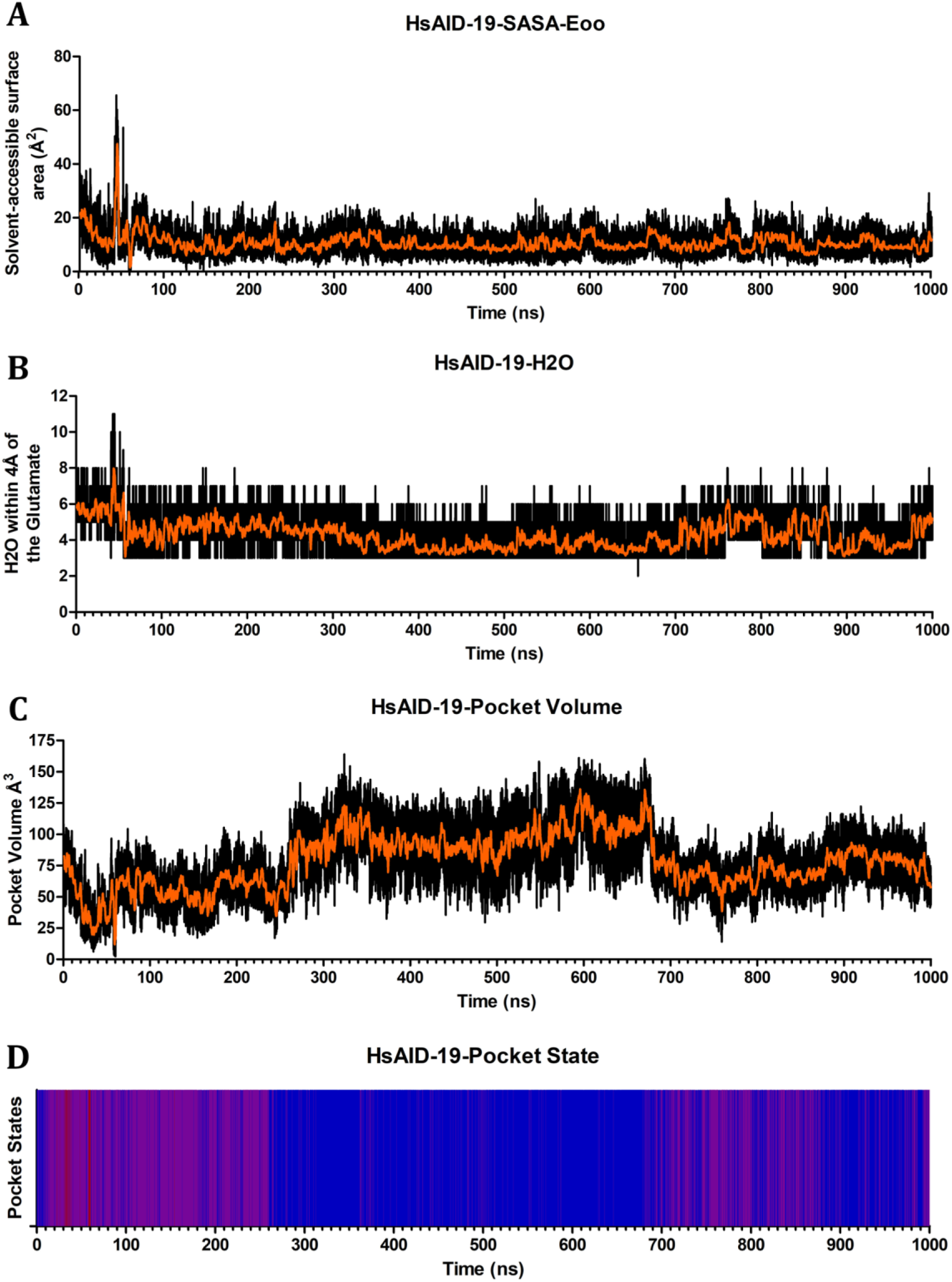
Pocket Dynamics of HsAID-β-state-19. Graphs depict HsAID-β-state-19 across 1μs simulation showing the (A) Solvent-Accessible Surface Area of the catalytic glutamates oxygens, (B) number of water molecules within 4Å of the catalytic glutamates oxygens, (C) pocket volume in Å^3^, and (D) pocket state coloured as open (blue), transient (purple), and closed (red).

**Figure S30.**
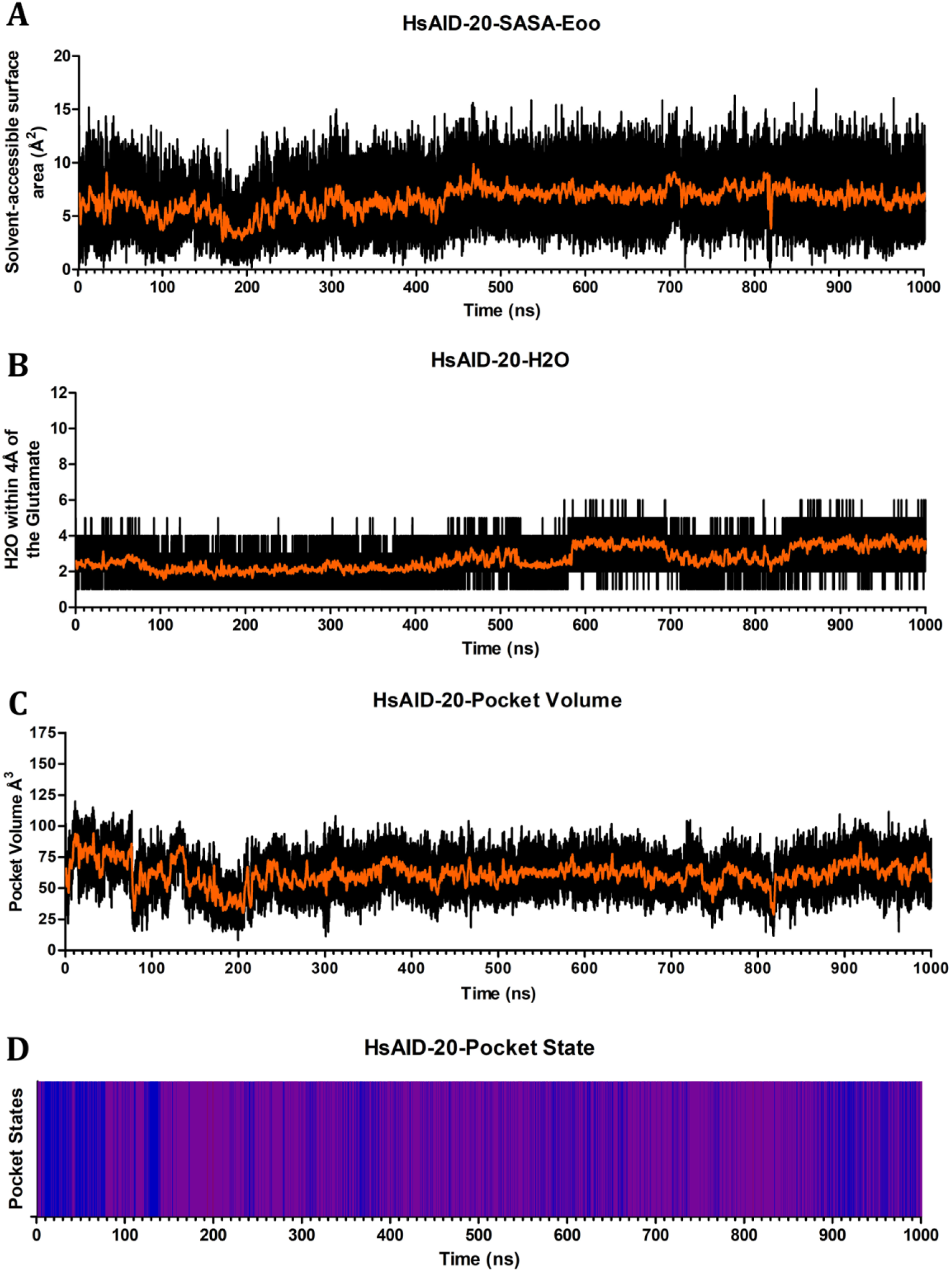
Pocket Dynamics of HsAID-β-state-20. Graphs depict HsAID-β-state-20 across 1μs simulation showing the (A) Solvent-Accessible Surface Area of the catalytic glutamates oxygens, (B) number of water molecules within 4Å of the catalytic glutamates oxygens, (C) pocket volume in Å^3^, and (D) pocket state coloured as open (blue), transient (purple), and closed (red).

**Figure S31.**
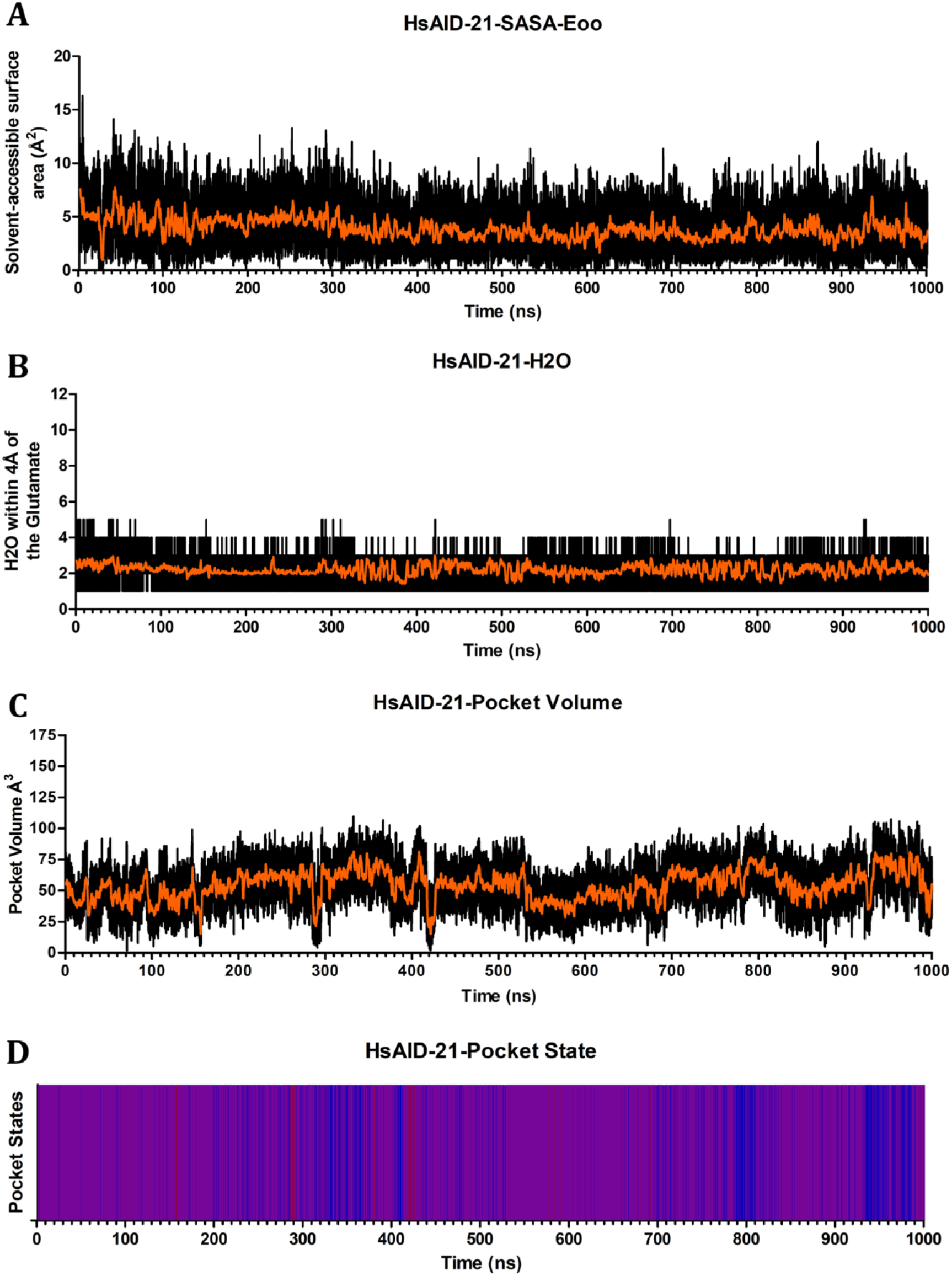
Pocket Dynamics of HsAID-β-state-21. Graphs depict HsAID-β-state-21 across 1μs simulation showing the (A) Solvent-Accessible Surface Area of the catalytic glutamates oxygens, (B) number of water molecules within 4Å of the catalytic glutamates oxygens, (C) pocket volume in Å^3^, and (D) pocket state coloured as open (blue), transient (purple), and closed (red).

**Figure S32.**
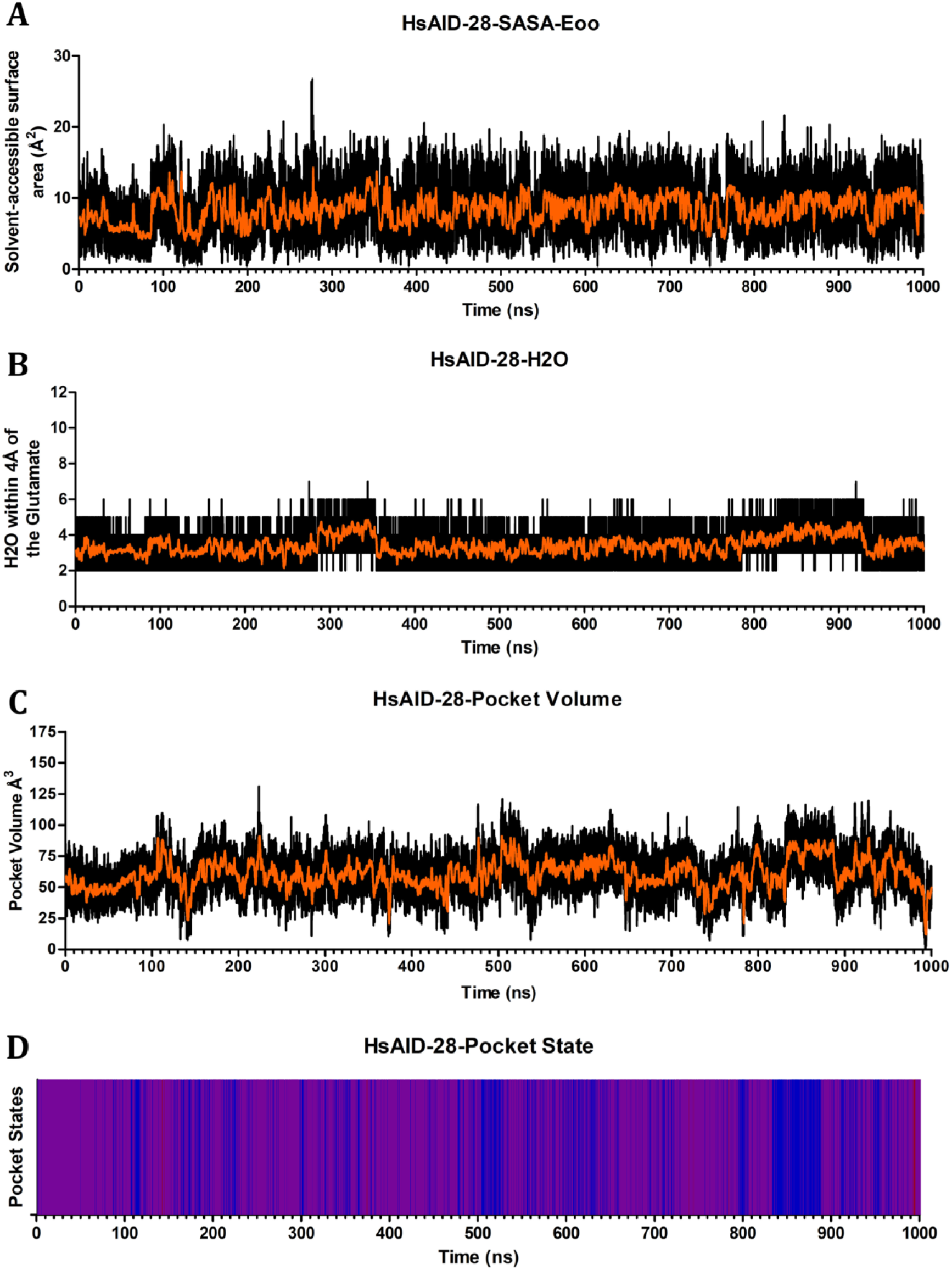
Pocket Dynamics of HsAID-β-state-28. Graphs depict HsAID-β-state-28 across 1μs simulation showing the (A) Solvent-Accessible Surface Area of the catalytic glutamates oxygens, (B) number of water molecules within 4Å of the catalytic glutamates oxygens, (C) pocket volume in Å^3^, and (D) pocket state coloured as open (blue), transient (purple), and closed (red).

**Figure S33.**
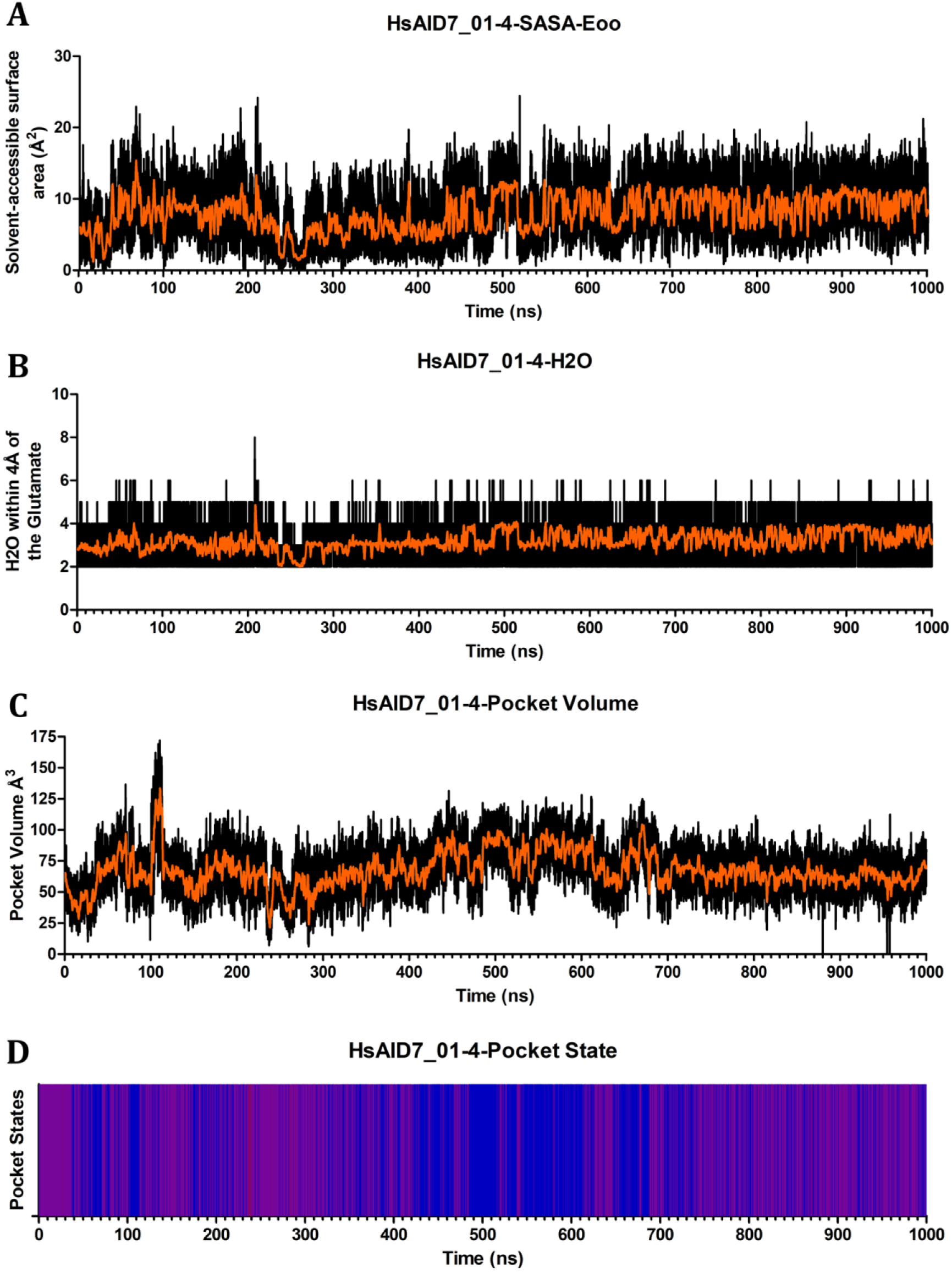
Pocket Dynamics of HsAID-α-state-7_01-4. Graphs depict HsAID-α-state-7_01-4 across 1μs simulation showing the (A) Solvent-Accessible Surface Area of the catalytic glutamates oxygens, (B) number of water molecules within 4Å of the catalytic glutamates oxygens, (C) pocket volume in Å^3^, and (D) pocket state coloured as open (blue), transient (purple), and closed (red).

**Figure S34.**
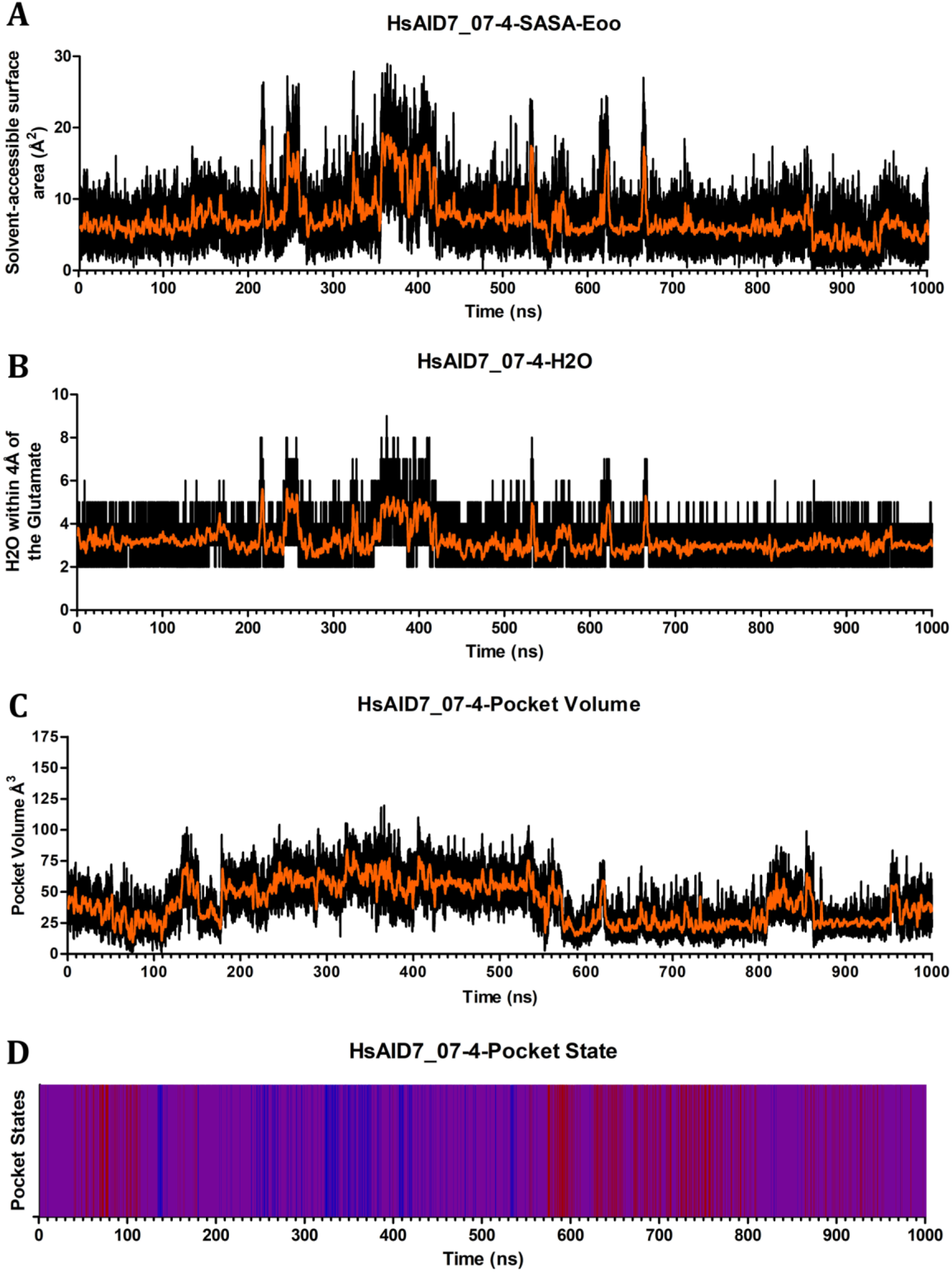
Pocket Dynamics of HsAID-α-state-7_07-4. Graphs depict HsAID-α-state-7_07-4 across 1μs simulation showing the (A) Solvent-Accessible Surface Area of the catalytic glutamates oxygens, (B) number of water molecules within 4Å of the catalytic glutamates oxygens, (C) pocket volume in Å^3^, and (D) pocket state coloured as open (blue), transient (purple), and closed (red).

**Figure S35.**
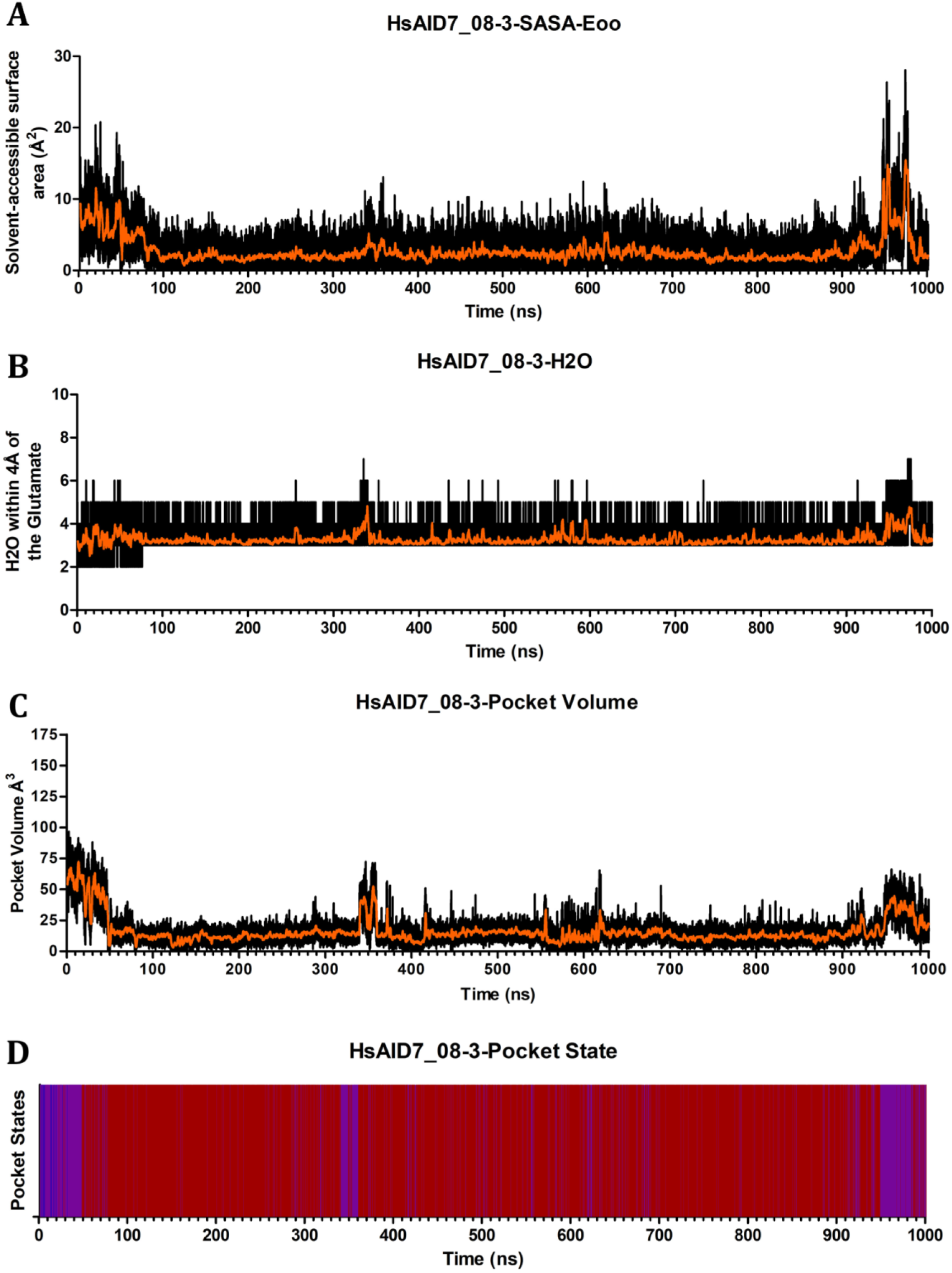
Pocket Dynamics of HsAID-α-state-7_08-3. Graphs depict HsAID-α-state-7_08-3 across 1μs simulation showing the (A) Solvent-Accessible Surface Area of the catalytic glutamates oxygens, (B) number of water molecules within 4Å of the catalytic glutamates oxygens, (C) pocket volume in Å^3^, and (D) pocket state coloured as open (blue), transient (purple), and closed (red).

**Figure S36.**
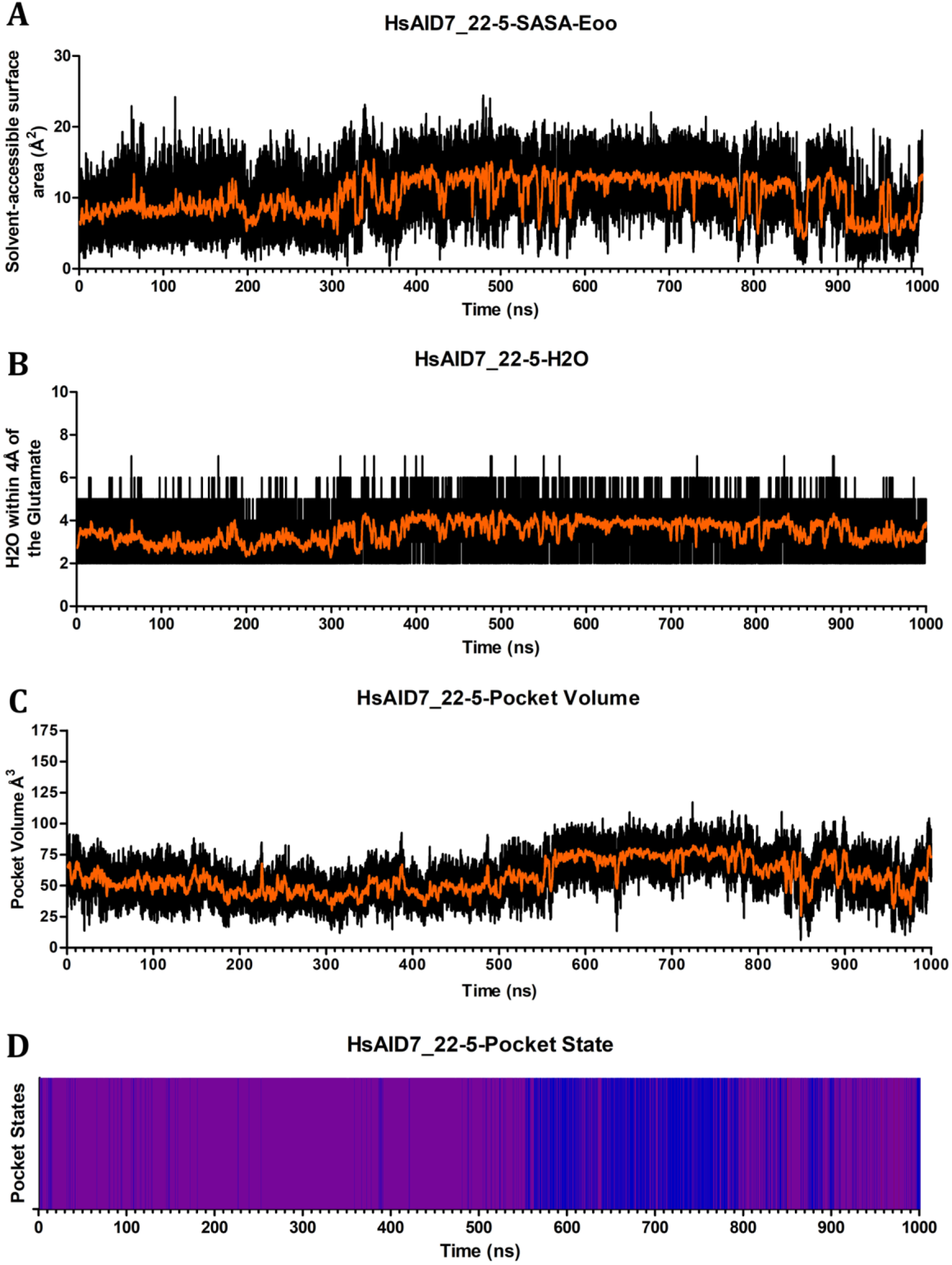
Pocket Dynamics of HsAID-α-state-7_22-5. Graphs depict HsAID-α-state-7_22-5 across 1μs simulation showing the (A) Solvent-Accessible Surface Area of the catalytic glutamates oxygens, (B) number of water molecules within 4Å of the catalytic glutamates oxygens, (C) pocket volume in Å^3^, and (D) pocket state coloured as open (blue), transient (purple), and closed (red).

**Figure S37.**
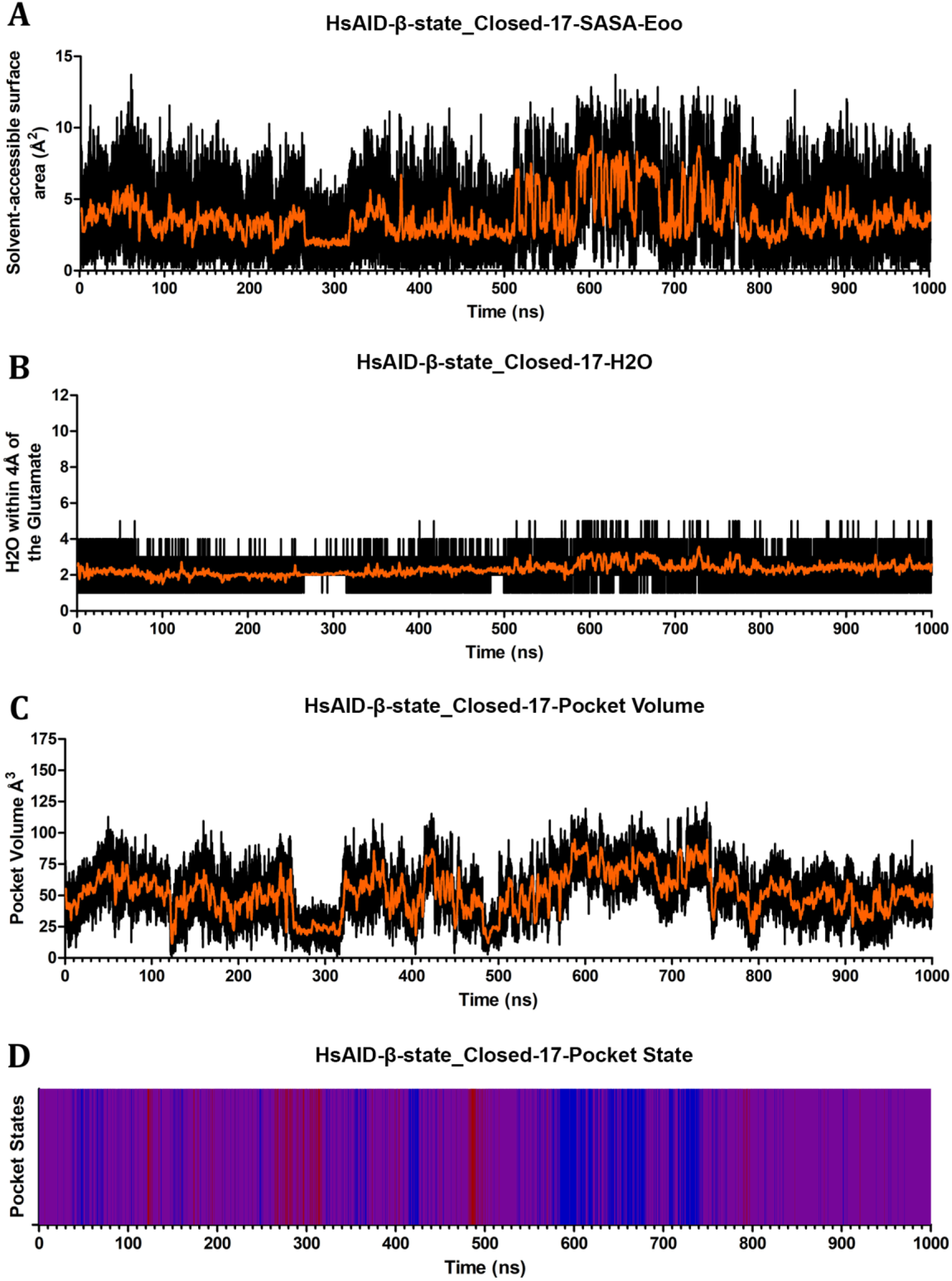
Pocket Dynamics of HsAID-β-state_Closed-17. Graphs depict HsAID-β-state_Closed-17 across 1μs simulation showing the (A) Solvent-Accessible Surface Area of the catalytic glutamates oxygens, (B) number of water molecules within 4Å of the catalytic glutamates oxygens, (C) pocket volume in Å^3^, and (D) pocket state coloured as open (blue), transient (purple), and closed (red).

**Figure S38.**
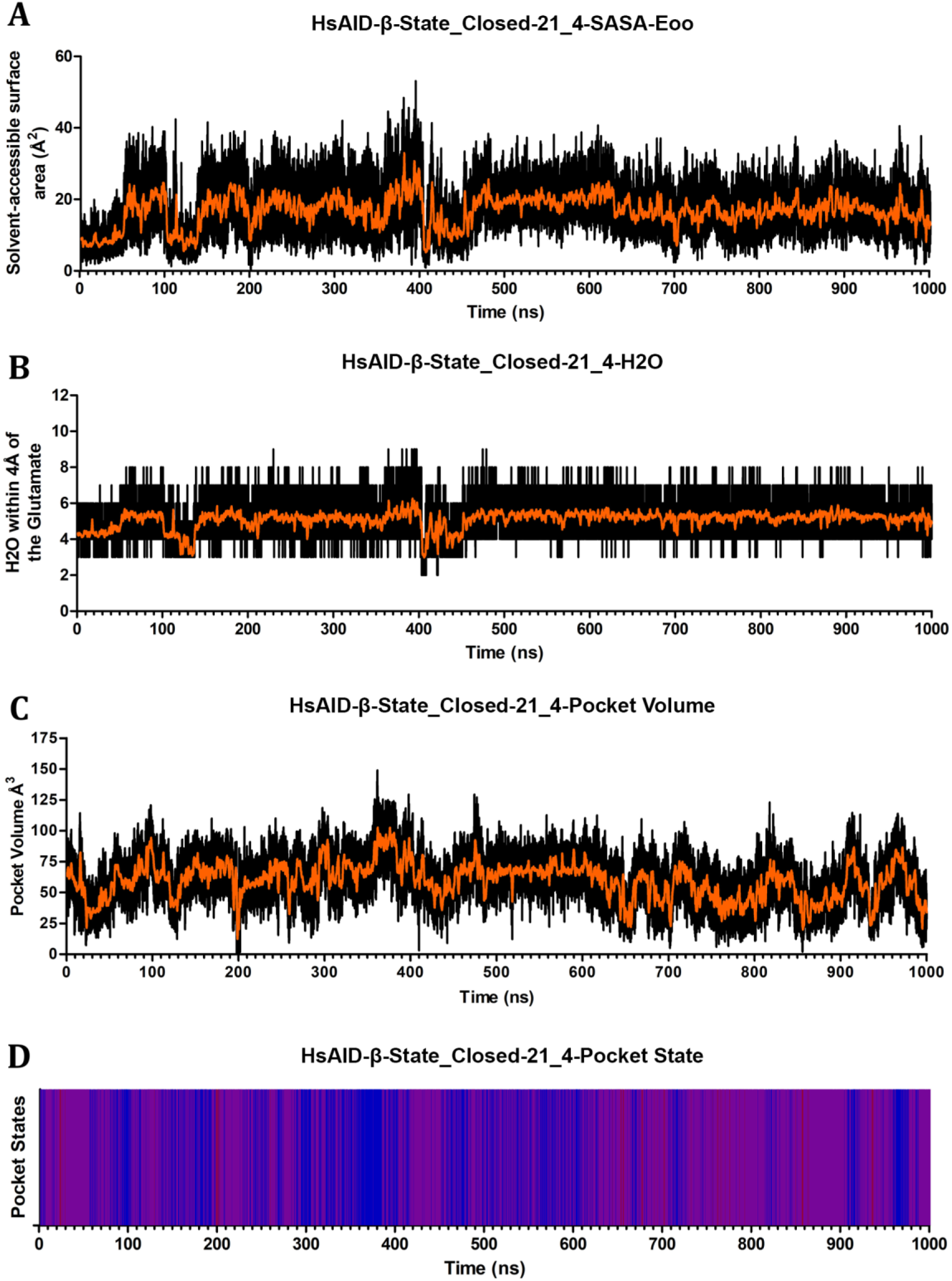
Pocket Dynamics of HsAID-β-state_Closed-21_4. Graphs depict HsAID-β-state_Closed-21_4 across 1µs simulation showing the (A) Solvent-Accessible Surface Area of the catalytic glutamates oxygens, (B) number of water molecules within 4Å of the catalytic glutamates oxygens, (C) pocket volume in Å^3^, and (D) pocket state coloured as open (blue), transient (purple), and closed (red).

**Figure S39.**
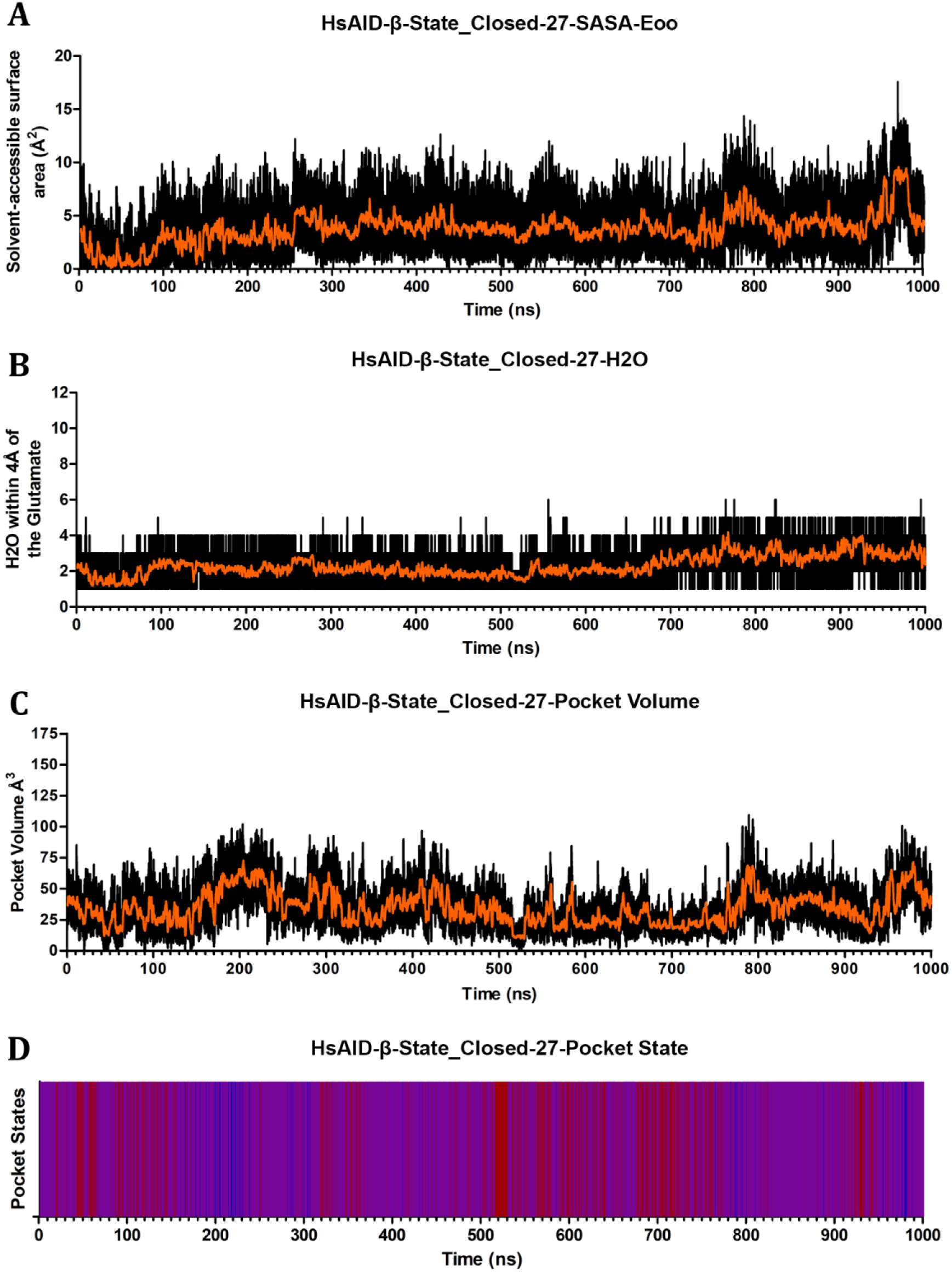
Pocket Dynamics of HsAID-β-state_Closed-27. Graphs depict HsAID-β-state_Closed-27 across 1µs simulation showing the (A) Solvent-Accessible Surface Area of the catalytic glutamates oxygens, (B) number of water molecules within 4Å of the catalytic glutamates oxygens, (C) pocket volume in Å^3^, and (D) pocket state coloured as open (blue), transient (purple), and closed (red).

**Figure S40.**
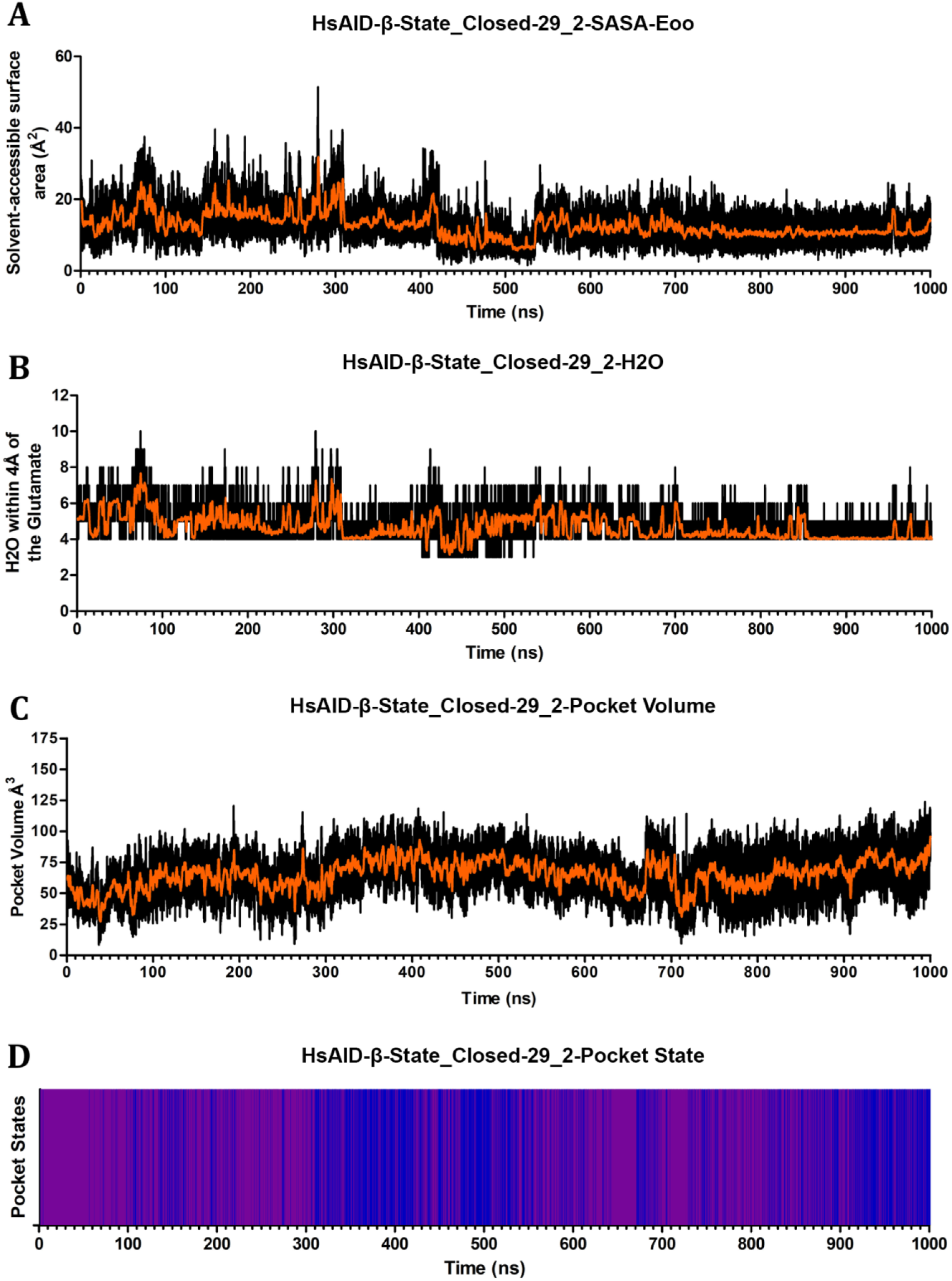
Pocket Dynamics of HsAID-β-state_Closed-29_2. Graphs depict HsAID-β-state_Closed-29_2 across 1µs simulation showing the (A) Solvent-Accessible Surface Area of the catalytic glutamates oxygens, (B) number of water molecules within 4Å of the catalytic glutamates oxygens, (C) pocket volume in Å^3^, and (D) pocket state coloured as open (blue), transient (purple), and closed (red).

